# Fast and Ultra-Capable Protein Design: Advancing the Frontier Through Atomistic SE(3)-Equivariance with Genie 3

**DOI:** 10.64898/2026.05.01.722168

**Authors:** Yeqing Lin, Minji Lee, Aakarsh Vermani, Ellena Jiang, Sebastiaan De Cooman, Matej Špeťko, Mohammed AlQuraishi

## Abstract

Despite the breakneck pace of progress in protein design methodology, frontier problems remain challenging, with leading methods struggling to design high-affinity binders, scaffold multiple functional motifs, or stabilize large multi-domain proteins. Recent research efforts have focused on two areas: improving model reasoning when generating active sites or binding interfaces, and improving concordance between the design process and the *in silico* oracle used to select promising designs. In addressing the first, the field has shifted towards all-atom models that capture sidechain conformations in atomistic detail by eschewing data-efficient SE(3)-equivariance, mirroring the evolution of AlphaFold2 to AlphaFold3. In addressing the second, recent work has focused on replacing generative models employing diffusion or flow-matching with hallucination approaches that directly optimize the oracle in sequence space; this improves success rates but reduces computational efficiency. Here, we close and surpass the generation-hallucination gap by revisiting SE(3)-equivariance using a branched polymer treatment of protein structures. The resulting diffusion model, Genie 3, achieves state-of-the-art performance on binder design, motif scaffolding, and unconditional generation, while being significantly faster than the best existing methods. We use Genie 3 to design a nanomolar binder of Nipah Glycoprotein G, a tetramer with minimal structural or biophysical characterization, as part of the Adaptyv Bio Nipah Competition, achieving a 12.5% success rate. Taken together, our results present a new frontier in protein design capability and a reexamination of the role of SE(3)-equivariance in molecular modeling.

## 1. Introduction

Recent advances in generative modeling of protein structures have revolutionized protein design, yielding *de novo* proteins with increasingly complex functions. One important application is the design of high-affinity binders that can act as therapies, regulate enzymes, and modulate cellular signaling. With AlphaFold’s achievement of highly accurate protein structure prediction, one approach to binder design, so-called hallucination-based, iteratively optimizes a protein sequence to maximize its AlphaFold-predicted binding to a target protein using a weighted combination of confidence metrics, interface contacts, and biophysical properties. This approach is exemplified by BindCraft (Pacesa et al., 2024), the current state-of-the-art method for binder design. Despite their success, hallucination-based approaches exhibit two major limitations: first, sequence optimization is performed using an *in silico* oracle—typically AlphaFold— which does not necessarily reflect experimental reality; second, hallucination requires iteratively running a computationally expensive structure prediction model, resulting in possibly intractable generation times for difficult targets.

An alternative to hallucination-based design is generative modeling via diffusion or flow matching, where a model of protein structure space is learned through iterative denoising. In this setting, binder design is performed by conditioning on the structure of the bound protein and, optionally, its interface residues (“hotspots”). Among generative methods for binder design, Proteina-Complexa represents the current state-of-the-art, exhibiting improved performance over BindCraft under matched compute budgets. The performance of generative methods depends not only on the quality of the learned distribution but also the sampling and selection strategies employed. When the effects of these components are disentangled—a contribution of this work—success rates for the underlying generative models, including Proteina-Complexa, fall far behind hallucination methods such as BindCraft. This indicates a gap between the two approaches: generative models can achieve strong overall performance when coupled with scaled inference-time sampling, but their learned distributions remain less concentrated on regions of structure space that map to successful binders. Here, we seek to bridge this gap by building a generative model whose learned distribution better reflects the space of foldable protein structures.

Recent generative models have shifted towards non-SE(3)-equivariant architectures, where proteins are represented in a manner dependent on their absolute orientation and position, a non-physical property. This shift, reflected by the evolution of AlphaFold2 (Jumper et al., 2021) to AlphaFold3 (Abramson et al., 2024), has been driven by the desire to capture atomistic details of sidechain conformations and, in some instances, to support modeling of non-polymeric small molecules. Non-equivariance requires data augmentation with random rotations and translations during training, incurring higher computational costs during model training. In this work, we revisit this design choice, by representing proteins as branched polymers, enabling atomistic reasoning over sidechain conformations while maintaining SE(3)-equivariance.

We build upon Genie 2, a structure diffusion model that employs asymmetric representations in the forward and backward diffusion processes. We introduce representational, architectural, and sampling modifications to enable the model to reason in an all-atom fashion. The resulting model, Genie 3, substantially outperforms Genie 2, achieving state-of-the-art performance on long monomer generation, motif scaffolding, and binder design. The latter represents a new functional capability for the Genie series of models, one that outperforms BindCraft’s binder design capabilities while being significantly faster. We furthermore show that Genie 3 and BindCraft explore different regions of protein structure, yielding highly complementary binder designs, suggesting a fundamental complementarity between hallucination- and generation-based approaches. To demonstrate Genie 3’s applicability to a real-world application, we use it to design a nanomolar binder of the Nipah Glycoprotein G after only eight attempts at *de novo* binder design.

## 2. Related Work

### Diffusion models

Diffusion models (Ho et al., 2020; Song et al., 2020b) are a class of generative models that map a predefined, easy-to-sample distribution, such as an isotropic Gaussian, to a complex data distribution such as the space of natural images, or, as in our case, the space of protein structures. In this work, we focus on diffusion in Euclidean space. Let **x** = [**x**^1^, **x**^2^, *…*, **x**^*N*^] denote an ordered set of coordinates of length *N*. The forward diffusion process iteratively injects isotropic Gaussian noise using a predefined variance schedule *β* = [*β*_1_, *β*_2_, *…, β*_*T*_], where *T* is the number of diffusion steps:

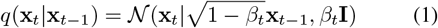

If the noise magnitude is sufficiently small at each step, we can approximate the corresponding reverse diffusion process using a Gaussian distribution:

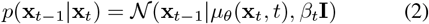

where

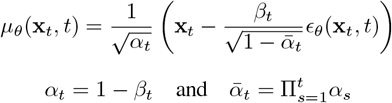

and *ϵ*_*θ*_ is a neural network trained to predict the noise injected at every time step *t*.

### Protein structure diffusion

The initial wave of protein structure diffusion methods focused on modeling protein backbones. FrameDiff (Yim et al., 2023b), FrameFlow (Yim et al., 2023a), Proteus (Wang et al., 2024), and RFDiffusion perform diffusion or flow matching in the SE(3) space of protein backbone frames, maintaining SE(3)-equivariance in the denoising process. These models rely on AlphaFold2-inspired architectures, with RFDiffusion being a directly fine-tuned variant of RosettaFold (Baek et al., 2021), which is itself adapted from AlphaFold2. Deviating from this recipe, Chroma (Ingraham et al., 2023) employs a correlated diffusion process that attempts to respect statistical properties of natural proteins, such as radius-of-gyration. Genie 1 and 2 use protein representations that are decoupled between the forward and backward diffusion processes; the former, point clouds, and the latter, reference frames. This enables rich SE(3)-equivariant reasoning in the backward process while retaining a very simple, non-equivariant noising process. Conversely, Proteina (Geffner et al., 2025b) performs diffusion in the Euclidean space of C_*α*_ atoms but use a non-equivariant transformer for denoising. Ambient (Daras et al., 2025) adopts Genie 2’s architecture while optimizing the training set through geometric clustering and careful integration of low-confidence *in silico* data.

More recent efforts have focused on all-atom diffusion inclusive of protein sidechain atoms. For structure-only models, this introduces a complication, as atom counts and identities depend on the amino acids comprising the protein sequence. RFDiffusion 2 (Ahern et al., 2025) adapts the RF-AllAtom (Krishna et al., 2024) architecture to a partial atomization scheme, where *a priori* known segments of a protein design are encoded using all amino acid atoms, while unknown segments are encoded using reference frames representing only backbone atoms. RFDiffusion 3 (Butcher et al., 2025) takes a different approach by modeling every residue using the largest canonical amino acid (tryptophan, with 4 backbone atoms and 10 sidechain atoms), effectively employing a redundant representation where extra atoms are collapsed onto the *C*_*β*_ atom. This is similar to Protpardelle-1c (Chu et al., 2024; Lu et al., 2025), a joint sequence-structure model, except the latter represents each residue as a “superposition” of all 20 possible amino acids (4 backbone atoms and 33 sidechain atoms); this non-physical “superposed” residue is then collapsed into a canonical amino acid once its identity is fixed during sampling. Continuing in this sequence-structure co-generation vein, La-Proteina (Geffner et al., 2025a) and Proteina-Complexa (Didi et al., 2025) use an autoencoder to encode sequence and atomistic sidechain information into a latent vector of fixed dimensionality, then train a flow matching model in the joint space of latent vectors and explicit *C*_*α*_ atoms. Proteina-Complexa introduces additional latent conditioning mechanisms and is trained on datasets of protein complexes, namely Teddymer and PLINDER (Durairaj et al., 2024), to support binder design against proteins and small molecule targets. Most all-atom models (except RFDiffusion 2) use a non-SE(3)-equivariant model for score (diffusion) or vector field (flow matching) prediction.

### Motif scaffolding

Different design methods expose different capabilities. Motif scaffolding, where the goal is to design proteins encompassing one or more functional substructures (“motifs”; *e*.*g*., enzyme active sites (Butcher et al., 2025), is one such capability. Wu et al. (2023) tackle motif scaffolding by proposing the Twisted Diffusion Sampler, a sequential Monte Carlo method that approximates the motif-conditional distribution by reweighting samples from an unconditional diffusion process. Similarly, Yim et al. (2024) start from an unconditional flow-based model and apply gradient-based guidance with respect to a structural loss between the target and generated motif, thereby biasing sampling trajectories toward motif-conditioned outputs. An alternative and more commonly adopted approach to conditional guidance is conditional training, as in Lin et al. (2024); Geffner et al. (2025a); Lu et al. (2025) and the motif-amortization approach of Yim et al. (2024), where the model is explicitly trained to scaffold randomly sampled motif segments. Recent all-atom models (Geffner et al., 2025a; Lu et al., 2025) encode motif sidechain atoms, with Lu et al. (2025) demonstrating that atomistic conditioning can be crucial to successful motif scaffolding.

### Binder design

Another important capability is the design of high-affinity binders to a protein of interest. Recent efforts can be grouped into two camps. The first, so-called hallucination approach, iteratively optimizes the binder sequence by maximizing its predicted binding to the target protein according to an *in silico* oracle such as AlphaFold. In practice, a weighted combination of AlphaFold-based confidence metrics, interface contacts, and predicted biophysical properties are used, and optimized sequences are further refined using an inverse folding model. BindCraft is a hallucination approach that uses AlphaFold-Multimer (AF2M) (Evans et al., 2021) as the oracle, while BoltzDesign1 (Cho et al., 2025) uses Boltz-1 (Wohlwend et al., 2025), an open-source derivative of AlphaFold3.

The second camp (Watson et al., 2023; Zambaldi et al., 2024; Stark et al., 2025; Didi et al., 2025) relies on generative models, where training (or fine-tuning) is explicitly performed using binder design tasks; for instance, by generating binders based on a target structure and prespecified hotspot residues corresponding to the target’s binding interface. Empirically, hallucination methods have so far outperformed generative models in binder design, at the cost of greater compute cost. It has been hypothesized that this may in part be due to the use of the *in silico* oracle as the optimization objective and the filtering criterion (after inverse folding), whereby the latter is made to directly align with the former. Note that this hypothesis is independent of the question of whether the oracle accurately predicts experimental success.

## 3. Methods

In this section, we describe the Genie 3 model architecture, training regimen, and training datasets. Where applicable, we also delineate differences from Genie 2.

### 3.1. Architecture verview

Genie 3 represents proteins as clouds of atoms in the forward process and as Frenet-Serret (FS) frames in the reverse process. It uses a standard forward diffusion process (Section 2) that adds isotropic Gaussian noise through a cosine variance schedule of *T* = 1000 diffusion steps. An SE(3)-equivariant denoiser, *ϵ*_*θ*_, reasons over reference frames to predict the noise injected during the forward process. This denoiser consists of (i) an input embedder that encodes noisy structure *x*_*t*_ and diffusion timestep *t* into residue-level (single) and residue-residue (pair) representations, (ii) a latent transformer that reasons over these representations, and (iii) an SE(3)-equivariant decoder that predicts denoised structures (Figure 1).

**Figure 1.**
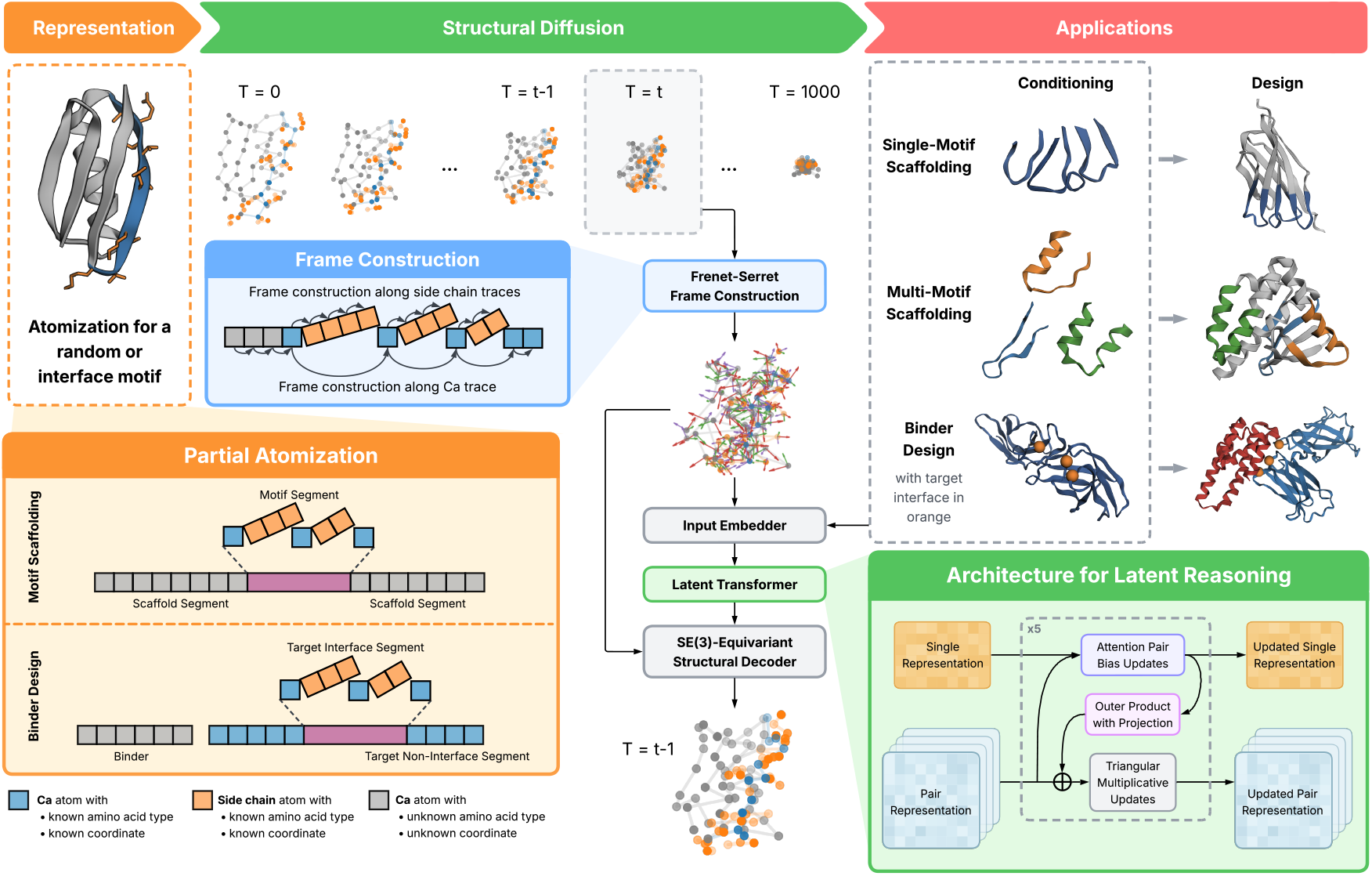
The Genie 3 Architecture. Proteins are represented as branched polymers with atomistic detail for sidechains when the sequence is known, such as in scaffolded motifs (orange inset). During the structural diffusion process, partially atomized proteins are represented as point clouds in the forward process and as frame clouds in the reverse process. Frames are computed using the Frenet-Serret construction (blue inset) along the backbone *C*_*α*_ trace and along each sidechain heavy atom trace (atom ordering is based on atom14 convention (Qu et al., 2024)). During each reverse diffusion step *t*, the denoiser predicts the injected noise at step *t* and the denoised structure at step *t−*1. The denoiser comprises three components: (1) an SE(3)-invariant input embedder that supports optional sequence and structure conditions depending on use case; (2) a latent transformer that iteratively operates on single and pair representations (see 3.1) to facilitate downstream structure decoding (green inset); and (3) an SE(3)-equivariant structural decoder that uses the mature single and pair representations from (2) along with the initial frames to predict a denoised structure for the next reverse diffusion step. During training, varying inputs are passed to the denoiser, to enable Genie 3 to support multiple downstream applications, including single- and multi-motif scaffolding and binder design.

The latent transformer in Genie 3 is substantially revised from Genie 2. In the latter, information flow between the single and pair representations occurs once at the beginning and end of the module. In Genie 3, information flow is bidirectional and occurs at every layer: pair representations are now updated by single representations via an outer product operation (Algorithm 3); single representations are updated by pair representations via their self-attention mechanism, by biasing this attention and injecting new values into the residual stream (Algorithm 2). We also introduce global tokens to the latent representation to promote global reasoning. The tokens are concatenated to the end of the single representations and removed prior to structural decoding. Algorithm 1 describes the overall latent reasoning module in detail.

For the structural decoder, Genie 3 uses Invariant Point Attention (IPA) (Jumper et al., 2021) to update single representations that are in turn used to update input reference frames. Final noise vectors are computed as the displacement between the translation component of the updated frames and that of the input frames. For more details refer to Lin et al. (2024); Lin & AlQuraishi (2023).

### 3.2. Motif atomization

As previously noted, an atomistic treatment of sidechains has been shown to be valuable for protein design models to achieve high-fidelity motif scaffolding. Backbone-only models can generate motifs with low *C*_*α*_ but high all-atom RMSD with respect to the target motif. In Genie 3, we adopt a partial atomization approach, using all-atom representations for known segments (*i*.*e*., motif residues) while operating exclusively over backbones for scaffold residues. To enable an SE(3)-equivariant treatment of the entire protein, we retain the frame-based representation of Genie 2 but generalize it to the branched polymer nature of proteins. For the backbone, we anchor Frenet-Serret (FS) frames at *C*_*α*_ atoms in relation to the FS frames of preceding *C*_*α*_ atoms, as in Genie 2. For sidechains, we start with the backbone FS frame then construct subsequent sidechain FS frames using the predefined ordering of the atom14 system (Qu et al., 2024). During training, we occasionally mask motif sidechain atoms and require the model to reconstruct these atoms given the motif sequence and, optionally, coordinates of the motif’s *C*_*α*_ atoms (see Appendix A.3.1).

### 3.3. Multimeric training

To equip Genie 3 with the ability to reason over multimeric complexes—necessary for binder design—we train it on monomeric and multimeric structural data. For monomeric structures, as in Genie 2, we use the Foldseek-clustered version (Barrio-Hernandez et al., 2023) of the AlphaFold Protein Structure Database (Varadi et al., 2022), filtering it for confidently predicted structures (average pLDDT *≥*80). We do not use any experimentally determined monomeric structures.

For multimers, we use the Pinder dataset, which comprises protein dimers extracted from the Protein Databank (PDB) and clustered by their interface structure similarity (Kovtun et al., 2024). During training, we first sample interface clusters, then sample protein complexes within clusters, to ensure equal representation at the interface level. For each chosen complex, we randomly treat one protein as the target and the other as the binder, and randomly mask half of the interface residues. Interfaces are defined as the set of residues whose *C*_*β*_ atom is within 8Å of a *C*_*β*_ atom of the other protein (*C*_*α*_ atoms are substituted for glycines).

We retain Genie 2’s input featurization except to add chain identifiers and binding interface mask tokens. To preserve representational expressiveness of the binding interface while ensuring computational feasibility, we only atomize interface residues. To promote atomistic reasoning at interfaces more generally, as in motif scaffolding, we occasionally mask out the coordinates of interface sidechain atoms and task the model with reconstructing them (Appendix A.3.2).

### 3.4. Heuristics for defining the binding interface

An algorithmic design choice subtlety arises when defining the target’s binding interface in training versus generation. During the latter, the target’s interface is generally treated to be a few target residues (known as hotspot residues). This is inconsistent or at least underspecified with respect to the interface definition used to condition the model during training (*C*_*β*_-*C*_*β*_ threshold of 8Å, as previously described), which typically spans larger regions, thus resulting in a distributional shift in the conditioner. To mitigate this issue, we explore two heuristic methods for expanding the set of interface residues during generation, given a set of starting hotspot residues. The first heuristic (termed “extended hotspots”) treats any target surface residue within 6Å of any hotspot residue as part of the interface (based on *C*_*β*_-*C*_*β*_ distances). We define surface residues to be ones with a Relative Solvent Accessibility (RSA) *≥* 0.25 and Solvent-Accessible Surface Area (SASA) 10Å^2^. The second heuristic (termed “convergent hotspots”) starts with the initial, user-specified hotspot residues as the interface, then adds interface residues that are found in the intersection of all successful designs from one or more previous design cycles. In this work, we use extended hotspots for monomeric targets and convergent hotspots for multimeric targets; for the latter, we start with a Genie 3-based design round of 100 structures to define the initial hotspots.

### 3.5. Low temperature sampling via directional scaling

Generative protein design models, including Genie 2, typically use lower sampling temperatures relative to training. Empirically, this improves the designability of generated proteins but introduces a discrepancy between training- and inference-time behavior. It is also a peculiarity specific to protein models, as generative models for other modalities, including vision, do not exhibit this phenomenon. Why this occurs has not been definitively resolved, but one possible reason is that many structures in the PDB (and AFDB) are too flexible—owing to their evolved origins—to be reliably designed by current generative models, as they rely on *in silico* folding models that in turn cannot reliably predict structures of natural proteins from single sequences. Lowering sampling temperatures effectively select for more rigid, and thus more designable (and predictable), proteins.

Technically, this is achieved by downscaling the noise at each reverse sampling step, resulting in lower noise variance levels than the corresponding diffusion time step *t* during training. Since *t* is passed as input to the denoising model, the expected noise level differs from the actual noise level in structure *x*_*t*_, with potentially detrimental effects on generative quality. To address these shortcomings in Genie 3, we pursue an alterative heuristic for low-temperature sampling by reformulating the reverse step (Equation 2) using the general approach of Denoising Diffusion Implicit Models (Song et al., 2020a):

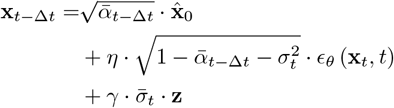

where

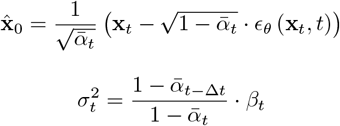

*η* is the direction scale and *γ*, Δ*t* = 10, is the noise scale. We set 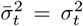, and the noise scale *γ* to 1. The reverse sampling process is moderated by the direction scale *η*, which controls the mean of the reverse Gaussian distribution at time step *t*. This is in contrast to Genie 2, where 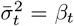, Δ*t* = 10, and the direction scale *η* is set to 1; the reverse sampling process is moderated by the noise scale *γ*, which controls noise variance in the reverse Gaussian distribution. Note that by setting Δ*t* = 10 in Genie 3, we reduce the number of sampling steps from 1,000 to 100, and thus reduce sampling time by 10x with negligible impact on generative quality.

## 4. Results

We tested Genie 3 and several leading protein design methods on unconditional generation, motif scaffolding, and binder design. To assess these methods at a scale sufficiently large to demonstrate statistical significance, we relied on *in silico* evaluation pipelines that test for self-consistency between generated structures and their corresponding predicted structures from AlphaFold-based models; this approach permits computational testing of thousands of designs. For targeted validation of Genie 3’s binder design capabilities, we experimentally tested eight designed binders of the Nipah Glycoprotein G (NiV-G).

### 4.1. Unconditional generation and motif scaffolding

To evaluate unconditional and motif scaffolding capabilities, we employed a widely used self-consistency pipeline (Watson et al., 2023; Lin et al., 2024; Geffner et al., 2025a) that inverse folds generated structures into novel sequences using ProteinMPNN (Dauparas et al., 2022), predicts the structures of these sequences using ESMFold (Lin et al., 2023), and tests for concordance between the two (for more details, see Appendix B.1). Highly concordant designs are considered successful. For unconditional generation, we consider designs ranging in length from 50 to 250 residues for short monomers and from 300 to 800 residues for long monomers, and generate 100 structures per sequence length per method tested. We remove all unsuccessful designs, then structurally cluster the remaining ones using a TM-score threshold of 0.5 to yield the final number of unique structural clusters. This approach quantifies not only the designability of generated proteins but their structural diversity as well. Genie 3 achieves comparable or better short monomer generation (*<*300 residues) performance relative to leading design models such as Ambient and La-Proteina (Appendix B.3), and exceeds their performance on long monomer generation (*>*300 residues) (Figure 2A). Genie 3 is only trained on structures up to 256 residues in length; its long monomer performance thus demonstrates out-of-domain generalization.

**Figure 2.**
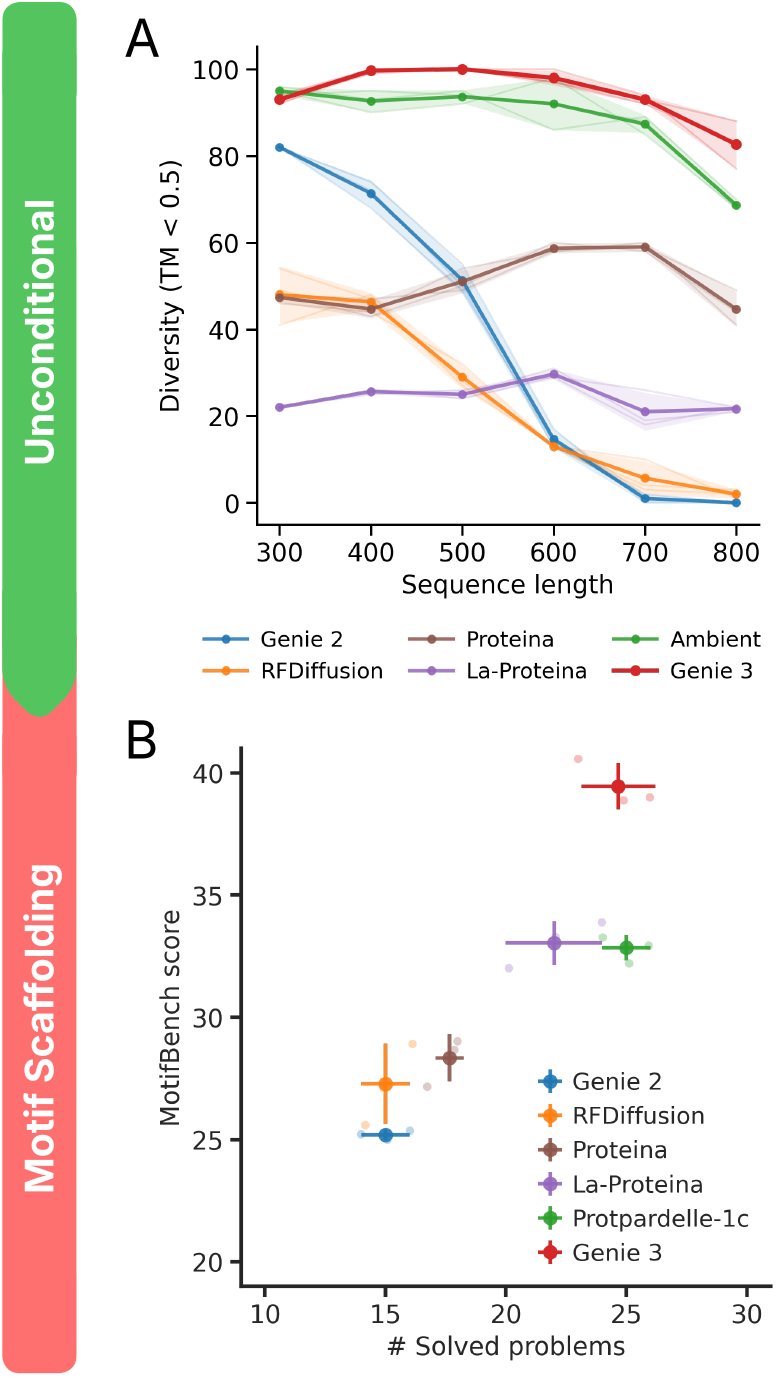
Comparative evaluation of unconditional generation and motif scaffolding. (A) Unconditional generation performance is assessed as a function of protein length (x-axis) using the number of structurally unique designed clusters (per 100 trials) as the evaluation criterion (y-axis). Plot shows mean diversity of three repeated samplings with shaded region indicating the extent of one standard deviation. (B) Motif scaffolding performance is assessed using MotifBench score (y-axis) as a function of the number of solved MotifBench problems (x-axis). Light points show three independent sampling runs. Dark points and error bars show the mean and standard deviation, respectively.

For motif scaffolding, we assessed performance using MotifBench (Zheng et al., 2025), a standardized benchmark comprising 30 distinct challenges. For each challenge, MotifBench computes a score as 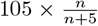, where *n* is the number of unique solutions generated by a design method. The overall MotifBench score is then computed as the average score across all challenges. This formulation results in diminishing returns as the number of solutions increase; the first few unique solutions yield a larger score increase than later solutions, favoring models that solve harder problems. Genie 3 solves as many problems as the best other method (Protpardelle-1c) while achieving a higher Motif-Bench score (Figure 2B). The problems solved by Genie 3 are also not the same as those solved by other methods, suggesting complementarity (details in Appendix C.3).

### 4.2. *In silico* binder evaluation pipeline

To evaluate binder design capabilities, we adapted the *in silico* evaluation pipeline of AlphaProteo (Zambaldi et al., 2024). An advantage of this pipeline is that its filtering power has been retrospectively assessed using the Cao et al. (2022) dataset, which comprises experimentally validated designs (both positives and negatives) for 11 different binder problems (See Appendix D.1 for details). Using these tested designs as ground truths, Zambaldi et al. (2024) quantified the precision of their evaluation approach. While imperfect, with high false positive rates and applied in the context of classically designed Rosetta-based *de novo* proteins, the AlphaProteo pipeline nonetheless quantifies the oracular power of the *in silico* oracle.

Starting with a designed binder-target complex, the pipeline uses AlphaFold3 to predict the structure using the multimeric sequence as the sole sequence input (without a multiple sequence alignment), together with the groundtruth target structure as template. The predicted and designed structures are then compared. The design is considered successful if it is a high-quality binder (pTM *>* 0.8 and minimum interaction pAE *<* 1.5Å) and is in close agreement with the predicted structure (complex RMSD *<* 2.5Å). We term this evaluation pipeline “AF3 Benchmark” to indicate its dependency on AlphaFold3. Its ability to select successful binders across the 11 aforementioned binder design tasks is summarized in Figure 3A.

**Figure 3.**
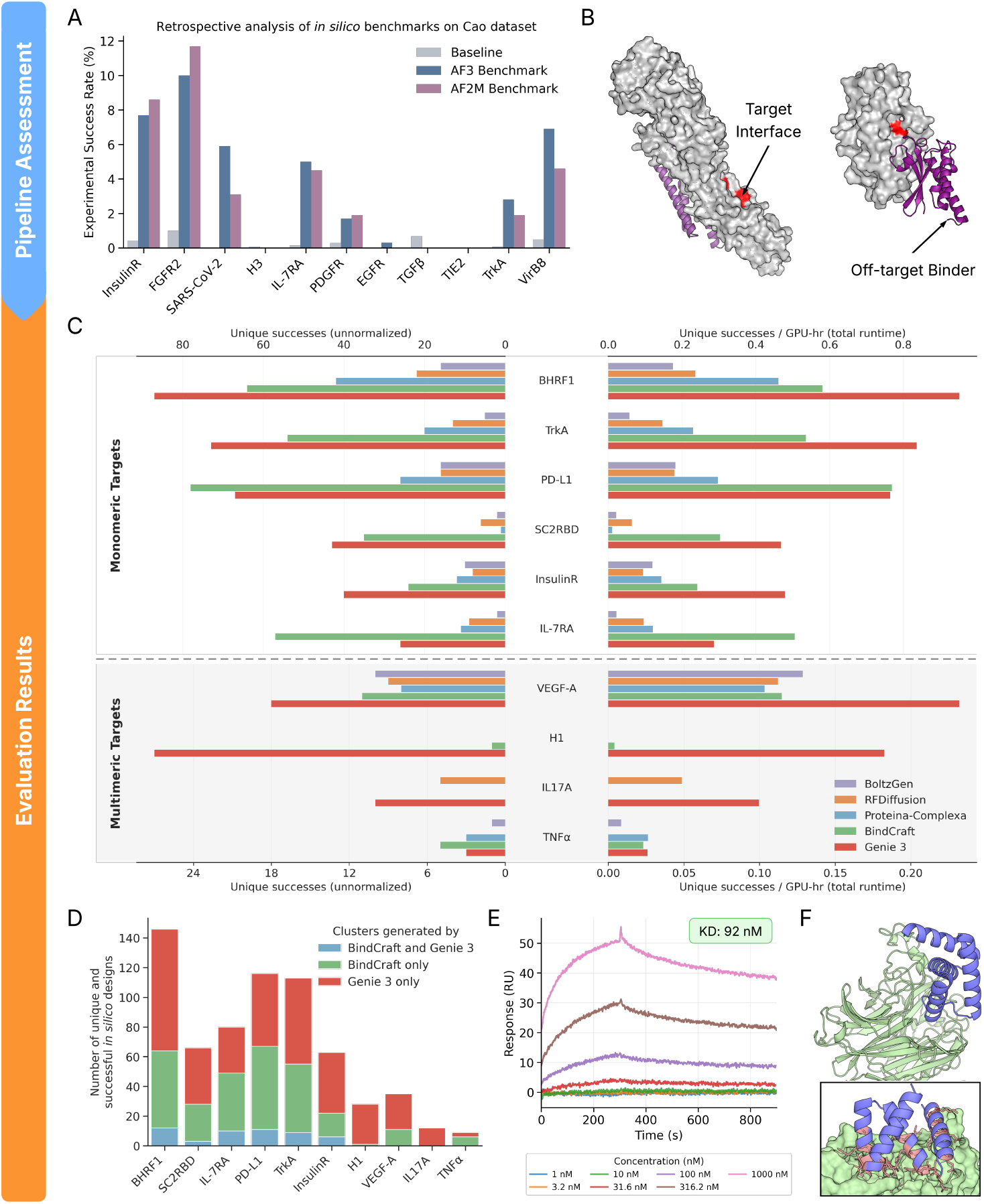
Comparative evaluation of binder design performance. (A) Assessment of the filtering power of multiple *in silico* binder evaluation pipelines (see Section 4.2). AF3 Benchmark and AF2M Benchmark were evaluated using the Cao dataset, which consists of 640K experimentally validated binders and non-binders across 11 design problems. We report the precision for every problem/benchmark combination, computed as the fraction of experimentally validated binders from the pool of designs deemed to be successful by an *in silico* benchmark. (B) Predicted structures of designed binders (violet) that pass the AF2M Benchmark criteria but fail to bind to the user-specified binding interface (yellow). (C) Number of unique (TM-score *≤* 0.6) and successful designs (as assessed by AF2M+ Benchmark) out of 200 attempts per problem for Genie 3 and four leading binder design methods across 6 monomeric and 4 multimeric binder design problems (left: unnormalized; right: normalized by number of A100 GPU hours used for generation and evaluation). (D) Successful designs unique to Genie 3, unique to BindCraft, and common to both as assessed by structural clustering at TM-score *≤* 0.6. (E)Binding response curves of Nipah Glycoprotein G to Genie-designed protein (*K*_*D*_ *≈* 92 nM). (F) AF2M prediction of Genie protein (blue) in complex with Nipah protein (green). Inset shows zoomed-in view of the binding interface.

Due to AlphaFold3’s restrictive license and current limitations of its existing reproductions, we adapt the pipeline to use AlphaFold-Multimer (AF2M) instead. We perform predictions with the 5 different AF2M models (each using up to 20 recycles with default early stopping) and select the top ranked prediction; rank score is computed as 0.8*·*ipTM+0.2*·*pTM. We term this pipeline “AF2M Benchmark” and find it be generally on par with AF3 Benchmark (Figure 3A). For certain challenging design problems, such as IL-17A, we found through our own retrospective analysis that inputting a multiple sequence alignment of the target sequence to AF2M, in lieu of a structural template, increases the true positive rate of AF2M Benchmark (Appendix D.5). We suspect that this may be due to the lack of multimeric template support in AF2M. Consequently, for challenging design problems such as IL-17A and TNF*α* (next section), we input multiple sequence alignments instead of templates of target structures.

We note that all evaluation pipelines exhibit low precision (maximum of 12%) and completely fail to identify binders against H3, TGF*β*, and TIE2. While this limitation is not specific to our approach, it does place a low ceiling on the efficiency of binder design using *in silico* oracles as filters. These filters are also insensitive to whether binding occurs at the user-specified interface or not. In Figure 3B, two designed binders technically pass the AF2M Benchmark but bind an off-target site. This issue is addressable and we tackle it here by introducing an additional constraint that 80% of user-specified interface residues must be within 5Å of the binder when the interface comprises 4 or more residues (100% for interfaces of 3 residues or less). Distances are computed based on the closest heavy atoms. We term the resulting pipeline “AF2M+ Benchmark” and use it for all downstream analyses.

### 4.3. Binder design

We assess the binder design capabilities of Genie 3, RFDiffusion, BoltzGen, Proteina-Complexa, and BindCraft, the last being the current state-of-the-art method for binder design. We challenge these models with 10 design problems curated by Zambaldi et al. (2024) that range from relatively easy targets such as BH3 helix (Procko et al., 2014) to very difficult ones such as tumor necrosis factor alpha (TNF*α*) (Hu et al., 2013). The problem set consists of 6 monomeric targets and 4 multimeric targets.

#### Fixed sampling budget

We first assess the quality of the underlying sampling distribution for each model by fixing the total number of generated structures to 200 for every model/problem combination and disabling inference-time search for Proteina-Complexa. We inverse fold 8 sequences per generated structure using ProteinMPNN and treat a generated structure as successful if any of its 8 designed sequence-structure pairs pass the AF2M+ Benchmark. We note that without inference-time search, Proteina-Complexa performs better using ProteinMPNN-based sequence redesign than using its own sequence co-generation feature (Appendix E.3).

Genie 3 produces the highest number of successful designs for 7 out of the 10 problems, followed by BindCraft, which yields the most solutions for the remaining 3 problems (Figure 3C; left). Despite training only on dimers, Genie 3 solves multimeric design problems as well as it does monomeric ones, suggesting generalization beyond its training set. Both Genie 3 and BindCraft substantially outperform RFDiffusion, BoltzGen, and Proteina-Complexa. Genie 3’s success indicates that a generative model, an aspect it shares with RFDiffusion, BoltzGen, and Proteina-Complexa, can outperform a far more expensive hallucination-based model.

Indeed, when we normalize model success rate by persample runtime, we find that the gap between Genie 3 and all other models persists (Figure 3C; right). As computational sampling is often a bottleneck in *de novo* protein design, this is a major practical benefit. Here, we compute per-sample runtime as the sum of:

- Generation time: time taken by a generative model to produce the initial structure (Appendix E.9.1).
- Inverse folding time: product of per-structure inverse folding time and number of inverse-folded sequences per structure. Per-structure inverse folding time is computed by problem and thus consistent across generative models (Appendix E.9.2). The number of inverse-folded sequences per structure is fixed at 8.
- Folding time: product of per-sequence structure prediction time and number of generated sequences for a structural design. Per-sequence structure prediction time is done using ColabFold with 5 AF2M models and 20 recycles with no early termination (Appendix E.9.3). This ensures equal structure prediction times across models. The number of generated sequences for a structural design is fixed at 8.

Genie 3 is also more faithful to user-provided hotspot specifications; for H1 and IL-17A, a sizable proportion of otherwise successful BindCraft designs fail to bind the specified interface (Appendix E.4). This may be due to an underlying difference between hallucination and generation: in the former, compliance with interface hotspots is implemented as an additional *ad hoc* loss term that is optimized during hallucination; in the latter, hotspots are provided as inputs to the model during training and generation, resulting in a shift in the underlying distribution of the generative process towards compliant designs.

Beyond raw capability, we sought to better understand the overlap in the structural distribution of designs proposed by BindCraft and Genie 3. We combined the successful designs proposed by both methods then structurally clustered them using a TM-score threshold of 0.6. In Figure 3D, we plot the number of designs falling into the same cluster and different clusters for each design method. Consistently across all tasks, few designs are produced simultaneously by the two methods, indicating that they identify complementary solutions to the same problems.

#### Inference-time scaling

We next enabled inference-time search for Proteina-Complexa, to assess its performance in its (native) inference-time scaling setting where the number of unique successes is normalized per GPU-hour. We did not reassess BindCraft since Genie 3 outperforms it under a fixed sampling budget, and per-sample generation times for Genie 3 are substantially shorter than for BindCraft. We run Genie 3 with one pass of ProteinMPNN-based sequence redesign, as this optimizes its inference-time scaling behavior (Appendix E.5). We run Proteina-Complexa with beam search, which uses AF2M-based interface predicted Aligned Error (ipAE) as the reward to jointly optimize the generated sequence-structure pair. We use Proteina-Complexa’s own co-generated sequences as they are the best performing in the beam search setting (Appendix E.3). We compute per-sample runtime as before, except for dropping inverse-folding time for Proteina-Complexa and setting the number of generated sequences per structure to 1.

Comparing the inference-time scaling behaviors of Genie 3 and Proteina-Complexa across 10 binder design problems, we observe that Proteina-Complexa outperforms Genie 3 on 1 problem, the two are tied on 2 problems, and Genie 3 outperforms Protein-Complexa on the remaining 7 problems (Figure 4A). In 4 instances, Proteina-Complexa exhibits no scaling behavior while Genie 3’s performance robustly improves with increased compute. These qualitative gains are largely concentrated among multimeric targets (3 out of 4), suggesting that they remain a substantial challenge for Proteina-Complexa and other protein design methods.

**Figure 4.**
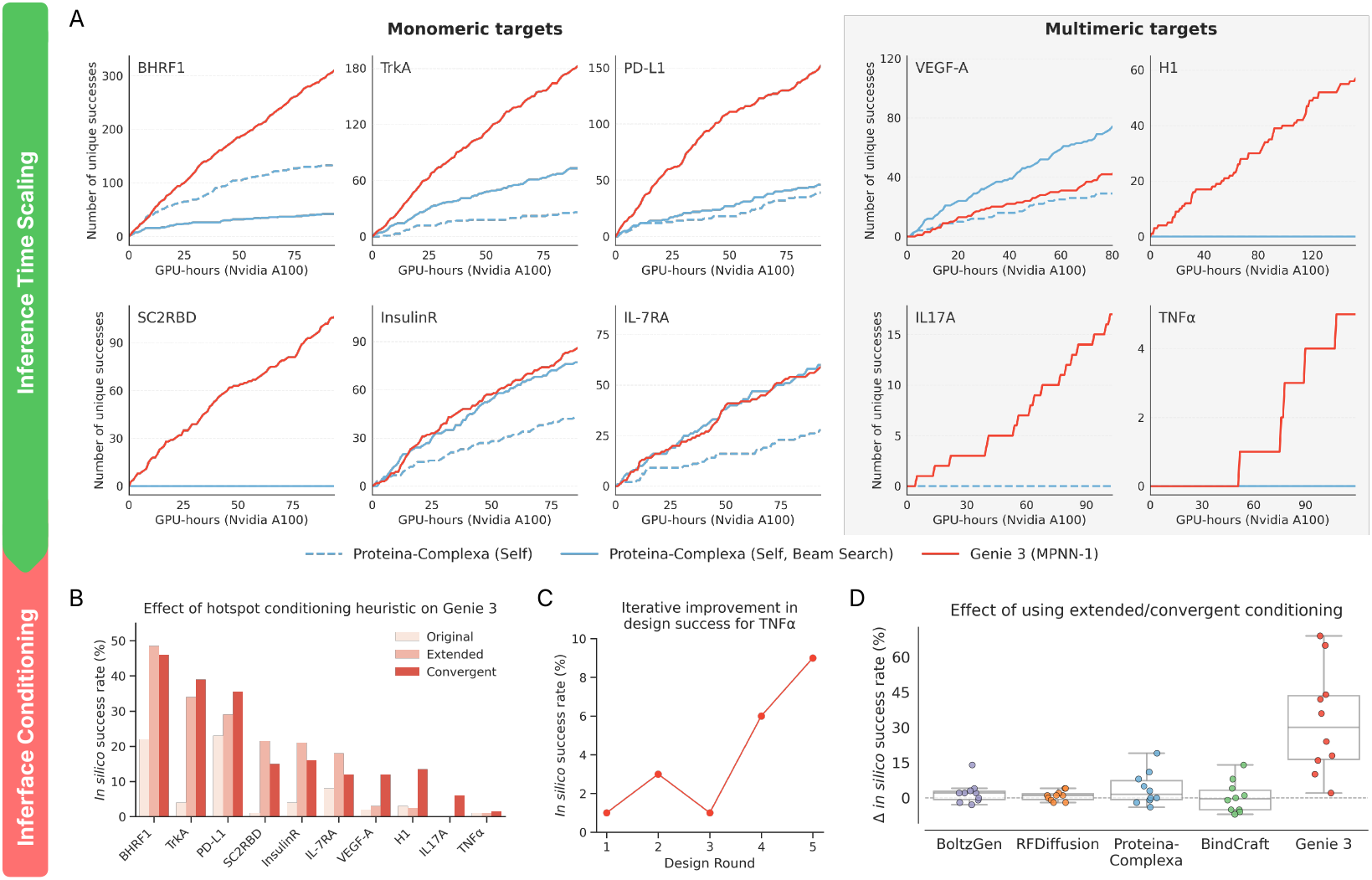
Impact of inference-time scaling and hotspot conditioning strategies on binder design performance. (A) Number of unique (TM-score *≤* 0.6) and successful (as assessed by AF2M+ Benchmark) designs as a function of Nvidia A100 GPU-hours (including generation and evaluation times) for 6 monomeric and 4 multimeric binder design problems. (B-C) Fractional rate of generating unique and successful Genie 3 designs when (B) using different hotspot conditioning heuristics and (C) using convergent hotspots to design TNF*α* binders as a function of number of design rounds (see Appendix E.8; 100 designs are sampled and evaluated per round). (D) Box plots of change in fractional rate of generating unique and successful designs when using extended/convergent hotspots versus original hotspots for Genie 3 and other leading methods.

### 4.4. Impact of binding interface expansion heuristics

To better understand how our binding interface heuristics impact Genie 3’s performance, we assessed its *in silico* success rates on the design problems of Zambaldi et al. (2024) by conditioning only on starting hotspot residues as well as on extended and convergent hotspots. Both expansion heuristics improve success rates, in some cases considerably (Figure 4B). Either approach works well for monomeric targets but convergent hotspots are substantially better for multimeric targets, including VEGF-A, H1, IL-17A, and TNF*α*. To further test this observation, we applied the convergent hotspots heuristic in an iterative manner to design a binder against TNF*α*, the most challenging target in our task set. We reasoned that repeated redefinition of hotspot residues may lead to further improvements. Indeed, we observed a nearly monotone climb in success rate as the number of design rounds increased, demonstrating inference-time scaling behavior in Genie 3 (Figure 4C). This is likely because prior design rounds inform the interfaces used in subsequent design rounds.

We note that this inference-time scaling behavior differs from that of Proteina-Complexa and what is shown in Figure 4A. In Genie 3, model conditioning transforms the underlying sampling distribution at every scaling stage, by injecting learned information from prior stages. Proteina-Complexa’s inference-time scaling behavior, on the other hand, results exclusively from repeated sampling, as it does not exploit accumulated knowledge across scaling stages, owing to its fixed sampling distribution. While *a priori*, the use of extended or convergent hotspots may have been expected to overly constrain generation and rigidify the binding pose, in practice it appears that such constraints help more than they hurt, perhaps through better alignment with the training data distribution.

Since the choice of hotspot residues is independent of design method, we next sought to determine if other models would similarly benefit from our interface expansion heuristics. We assessed RFDiffusion, BoltzGen, Proteina-Complexa and BindCraft using the same methodology as before and examined the differences in their success rates relative to using the original hotspots (Figure 4D). Unlike Genie 3, the other models show small or negligible improvements (Appendix E.7). We hypothesize that this may be due to the differing formulations of the binder design training task in other models. For instance, RFDiffusion is trained by providing at most 20% of hotspot residues, versus 50% for Genie 3, rendering the model less constrained by the user’s specification. The balance between overly rigidifying the binding pose and providing useful priors may also play out differently for the other models.

### 4.5. Designing a novel binder against Nipah NiV-G

For a targeted demonstration of Genie 3’s design capabilities, we participated in AdaptyvBio’s Nipah binder design challenge, which was aimed at designing high-affinity binders against Nipah virus Glycoprotein G (NiV-G). NiV-G binds the Ephrin-B2 and Ephrin-B3 cell surface receptors of human cells, initiating attachment to host cells and subsequent pathogenesis; a competitive binder of NiV-G could thus act to inhibit viral infection. The challenge was focused in scope and did not broadly test for selectivity or any *in vivo* properties, but the identification of high-affinity *in vitro* binders does provide a starting point for further therapeutic development. Up to 10 designed sequences could be submitted per team, from which the top designs were chosen based on their score on the Boltz-2 ipSAE metric (Passaro et al., 2025; Dunbrack Jr, 2025). Selected designs were engineered with a C-terminal Twin-Strep-tag and synthesized using a call-free protein expression system. Following synthesis, the resulting crude protein mixtures were used for the site-specific immobilization of the binders onto Strep-Tactin XT-functionalized chips. Binding kinetics were characterized via surface plasmon resonance (SPR) by exposing the functionalized sensors to a concentration gradient of purified recombinant NiV-G analyte (residues 71-620) ordered from Twist Biosciences.

To design new binders, we leveraged a publicly available crystal structure of NiV-G in complex with human Ephrin-B2 (PDB: 2VSM) to constrain the target interface region to the set of residues within 8Å of Ephrin-B2. We used Genie 3 to generate 200 designs then filtered them using our AF2M+ Benchmark to obtain 34 successful designs spanning 33 unique structural clusters (TM-score *>* 0.6). Since selection of binder candidates is based on the Boltz-2 ipSAE metric, we computed it for all our designs and selected the top 8 for submission. All 8 expressed and one had a measurable *K*_*D*_ of 92nM (Figure 3E and 3F), corresponding to a 1/8 hit rate. For reference, submissions made using RFDiffusion, BindCraft, and BoltzGen had aggregate hit rates of 3/60, 1/100, and 2/288, respectively (more details in Appendix E.10). We note that these hit rates encompass diverse generation and filtering procedures performed by groups with varying levels of expertise, and may thus not be representative of their corresponding models’ performance in a controlled experimental setting.

### 4.6. Architectural and sampling ablations

Genie 3 introduces architectural as well as sampling innovations over its predecessor. To better understand the individual contributions of these changes, we built ablated variants of Genie 3 and assessed their performance on unconditional monomer generation, which we consider to be an indicator of a model’s raw generative capacity. We compared Genie 3 to Genie 2 (baseline model without modifications) as well as two Genie 3 variants, one without the sampling modifications described in Section 3.5, and one without the architectural and training modifications described in Sections 3.1 and 3.3. As shown in Figure 5, ablating these modifications leads to a performance drop, with sampling modifications responsible for the greatest gains across all sequence lengths. The architectural and training modifications are also important, but only impact designs longer than 700 residues in length. Combined with the fact that Genie 3’s multimeric training dataset is restricted to complexes shorter than 256 residues, our architectural changes appear to be critical for generalization to larger complexes.

**Figure 5.**
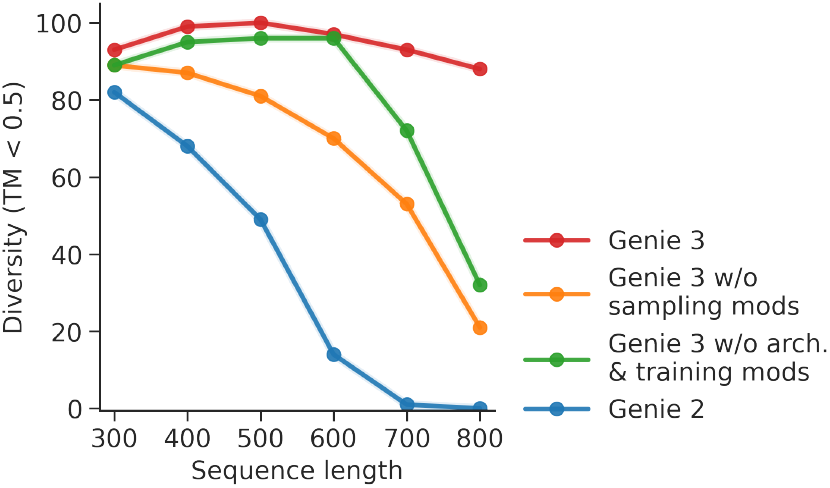
Comparative evaluation of unconditional generation performance using Genie variants. Performance is assessed as a function of protein length (x-axis) using the number of structurally unique designed clusters (per 100 trials) as the evaluation criterion (y-axis).

## 5 Conclusions

Genie 3 introduces architectural, sampling, and interface conditioning innovations to achieve state-of-the-art performance on unconditional generation, motif scaffolding, and binder design. It demonstrates that generative models can equal or even surpass hallucination-based approaches on difficult design problems, while being orders of magnitude faster in generation. In doing so, it illustrates that SE(3)-equivariance remains a powerful and expressive inductive bias for atomistic molecular models.

Nonetheless, limitations remain. Our assessments were dependent on *in silico* evaluation pipelines that in turn depend on imperfect protein structure prediction models. The inability of such models to accurately score protein-protein complexes and produce alternate protein conformations is well known, and likely limits their discriminative utility in protein design. A potential future direction for Genie is to integrate experimental feedback into its design process to steer generation directly towards consistency with experimental observables. Genie 3 and other generative binder design models also assume that target structures accurately reflect the conformation of the bound state, which ignores induced fit effects. The ability to design binders against unknown conformations is thus a key future direction, one that may leverage conformational sampling methods such as MSA subsampling (Wayment-Steele et al., 2024), BioEmu (Lewis et al., 2025) and ConforNets (Lee et al., 2026).

## Acknowledgments

We thank Tony Culos for insightful discussions and feedback. Research reported in this publication was supported by the National Institutes of Health under award number R35GM150546. This work used resources of the National Energy Research Scientific Computing Center (NERSC), a Department of Energy User Facility using NERSC award DDR-ERCAP0037622. We also acknowledged the EuroHPC Joint Undertaking for funding and access to Karolina at IT4Innovations, Czech Republic (project ID: EHPC-REG-2023R03-016), and the Ministry of Education, Youth and Sports of the Czech Republic (EPICURE, ID: MC2402) for additional support.

## A. Additional Information

### A.1. Features

### A.2. Improved latent reasoning via LatentTransformer

In Genie 3, we replace the PairTransformNet of Genie 2 with an improved latent reasoning module we term LatentTransformer.

In this section, we provide the detailed algorithm for LatentTransformer. Here,

- PairTransition, TriangleMultiplicationOutgoing, and TriangleMultiplicationIncoming follow from AlphaFold 3 (Abramson et al., 2024) Algorithms 11, 12 and 13 respectively.
- AddTrailingTokens adds global tokens, whose single and pair representations are initialized with zeros, while RemoveTrailingTokens drops global tokens post latent transformation.
- AttentionPairBiasUpdate is adapted from the Invariant Point Attention algorithm from AlphaFold 2 (Jumper et al., 2021) (Algorithm 22), without the integration of frame information.

#### Algorithm 1

LatentTransformer

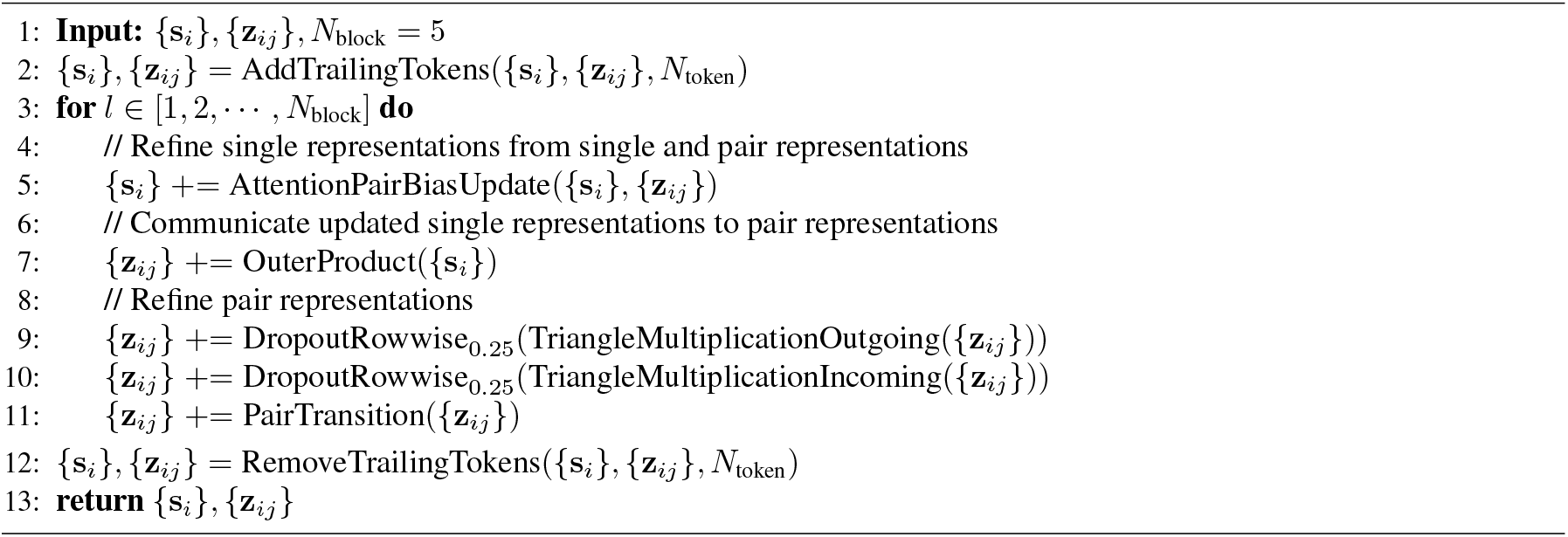

#### Algorithm 2

AttentionPairBiasUpdate

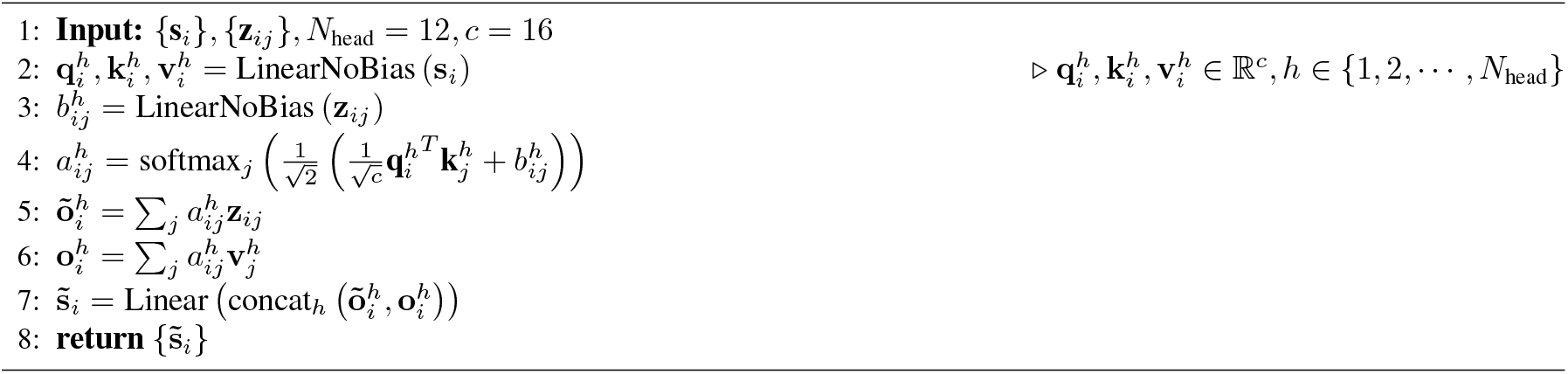

#### Algorithm 3

OuterProduct

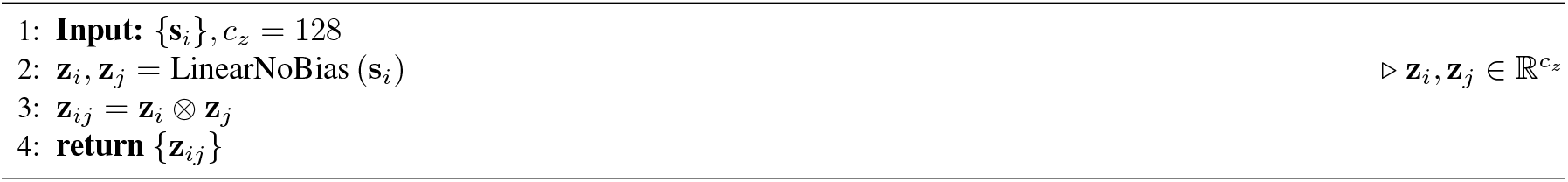

### A.3. Training details

We train Genie 3 using a multi-task training setup, with a 9:1 ratio between motif scaffolding training tasks (Appendix A.3.1) and binder design tasks (Appendix A.3.2). We train Genie 3 on 48 Nvidia A100 40GB GPUs for ~15 days, totaling 600,000 training steps. Per-GPU batch size is set to 1, resulting in an effective batch size of 48. Gradient updates are performed using an Adam optimizer with a learning rate of 0.0001.

#### A.3.1. Motif scaffolding tasks

We use the same training dataset as Genie 2, which consists of FoldSeek-clustered AlphaFold Database representative clusters filtered to sequences ranging in length between 20 and 256 residues (inclusive) and a minimum pLDDT of 80. This results in 587,479 structures. During training, we use the same motif selection algorithm detailed in Algorithm 1 of Genie 2 but further consider three different types of motif scaffolding training tasks depending on the amount of motif information provided as conditions to the model. These tasks are summarized in the table below, with the corresponding ratio of each task used during training.

|  | Task 1 | Task 2 | Task 3 |
| --- | --- | --- | --- |
| **Motif Information** |  |  |  |
| sequence | ✓ | ✓ | ✓ |
| backbone structure | ✓ | ✓ |  |
| side-chain structure | ✓ |  |  |
| Task Ratio | 0.4 | 0.4 | 0.2 |

##### A.3.2. Binder design tasks

We train on the training split of the 2024-02 version of the Pinder dataset. We further filter out structures with missing residues and atoms and cap the maximum number of residues in complexes to 256. The resulting Pinder training dataset consists of 135,745 structures with 5,331 interface clusters. During training, we first sample a groundtruth complex based on interface clusters before sampling within the selected interface cluster. For each sample, we randomly consider one chain as the target and the other as the groundtruth binder. Interface is computed with a Cb-Cb threshold of 8Å and the interface mask is randomly masked at a 50% ratio. We consider five different types of binder design training tasks depending on the amount of target and/or binder information provided as conditions to the model. These tasks are summarized in the table below, with the corresponding ratio of each task used during training.

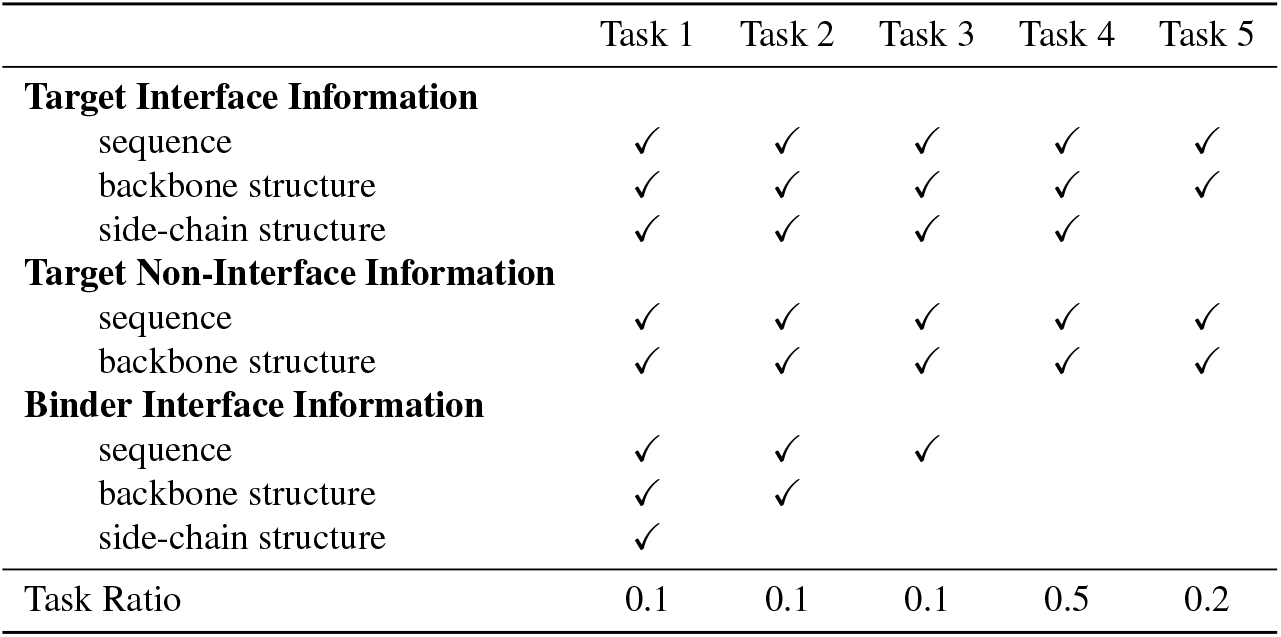

|  | Task 1 | Task 2 | Task 3 | Task 4 | Task 5 |
| --- | --- | --- | --- | --- | --- |
| **Target Interface Information** |  |  |  |  |  |
| sequence | ✓ | ✓ | ✓ | ✓ | ✓ |
| backbone structure | ✓ | ✓ | ✓ | ✓ | ✓ |
| side-chain structure | ✓ | ✓ | ✓ | ✓ |  |
| **Target Non-Interface Information** |  |  |  |  |  |
| sequence | ✓ | ✓ | ✓ | ✓ | ✓ |
| backbone structure | ✓ | ✓ | ✓ | ✓ | ✓ |
| **Binder Interface Information** |  |  |  |  |  |
| sequence | ✓ | ✓ | ✓ |  |  |
| backbone structure | ✓ | ✓ |  |  |  |
| side-chain structure | ✓ |  |  |  |  |
| Task Ratio | 0.1 | 0.1 | 0.1 | 0.5 | 0.2 |

## B. Unconditional Generation

### B.1 Evaluation metrics

Following existing literature (Watson et al., 2023; Lin et al., 2024; Geffner et al., 2025a) on protein generative model evaluations, we use a self-consistency pipeline to assess the quality of generated structures. Given a generated structure, we run ProteinMPNN (Dauparas et al., 2022) to generate a sequence and fold the sequence using ESMFold (Lin et al., 2023). We measure the quality in terms of consistency between the generated structure and the ESMFold-predicted structure, which is measured in terms of Kabsch-aligned Root Mean Squared Distance (RMSD) on C_*α*_ atoms. This metric is commonly known as self-consistency RMSD (scRMSD).

#### Designability

A generated structure is considered designable if its minimum scRMSD *<* 2Å (across 8 ProteinMPNN-predicted sequences). Designability is defined as the percentage of designable structures.

#### Diversity

We use the following FoldSeek (Van Kempen et al., 2024) command to cluster designable structures and diversity is reported as the number of unique clusters at a given TM threshold. We used FoldSeek release 10 (January 19, 2025); results may vary with different versions. In particular, we observed an approximately 0.05 change in designability (*e.g*., 370 vs. 345 clusters out of 500 structures) when using a more recent release, likely due to bug fixes in the TM-align parallelization pipeline. To ensure consistency and fairness, all evaluations were conducted using the exact same version.

~~~
foldseek easy-cluster \
    $INPUT_DIR \
    $OUTPUT_DIR/diversity \
    $TEMP_DIR \
    --cov-mode 0 \
    --alignment-type 1 \
    --min-seq-id 0 \
    --tmscore-threshold $TM_THRESHOLD
~~~

#### Novelty

We use the following FoldSeek command to compute the closest structure in the reference database 𝒟_ref_ for each designable structure *x ∈ 𝒟*_g_. This skips the prefiltering stage and uses TMAlign to compute structural similarity between each designable structure and every structure in the reference dataset (denoted as function *f*).

~~~
foldseek easy-search
    $INPUT_DIR
    $DATABASE_DIR
    $OUTPUT_FILEPATH
    $TEMP_DIR
    --alignment-type 1
    --exhaustive-search
    --tmscore-threshold 0.0
    --max-seqs 10000000000
    --format-output query,target,qtmscore,lddt
~~~

For reference databases, we use the built-in PDB database from FoldSeek and our clustered AFDB training dataset. Novelty is computed as

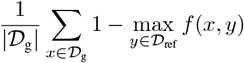

One caveat is that even with exhaustive search, FoldSeek might not find local alignments between a designable structure and a structure in the reference database. In these case, we assume the maximum novelty score of 1.

#### Secondary structure assessment

We assign secondary structures using the DSSP algorithm. We report the overall percentage of helices and strands by averaging the observed percentages of helices and strands across all designable structures, respectively.

### B.2 Reproduction of baseline methods

We provide details on the reproduction of baseline methods for unconditional generation below.

#### RFDiffusion

We followed instructions from the Github repository^1^ using the pretrained weights located at http://files.ipd.uw.edu/pub/RFdiffusion/. We use the provided default configuration, which sets the noise scale to 1.

#### FrameFlow

We followed the instructions from the Github repository^2^ and used the provided pretrained weights located at the checkpoint weights/pdb/published.ckpt for sampling.

#### Proteus

We followed the instructions from the Github repository^3^ and used the pretrained weight provided under ./weights/-paper weights.pt, utilizing the default parameters in the repository for unconditional generations.

#### FoldFlow 2

We followed the instructions from the Github repository^4^ and evaluated its unconditional generation performance with both ff2 base and ff2 reft checkpoints, where ff2 reft is finetuned for secondary diversity.

#### Proteina

We followed the instructions given in the Github repository^5^. For proteins shorter than 300 residues, we use *M*_FS_ with a noise scale of 0.5. For longer proteins with at least 300 residues, we use *M*_long_ with a noise scale of 0.35.

#### La-Proteina

We followed the instructions given in the Github repository^6^ and used the provided pretrained weights for sampling. For proteins shorter than 300 residues, we used the checkpoints of LD1 (no triangular layers) and LD2 (triangular layers) with noise scales of 0.1 for both the alpha carbon atoms and for the latent variables, combined with the AE1 checkpoint for the autoencoder. For longer proteins with a length of at least 300 residues, we used the checkpoint at LD3 with noise scales of 0.15 for the alpha carbon atoms and 0.05 for the latent variables, combined with the AE2 checkpoint for the autoencoder.

#### Protpardelle-1c

We followed the instructions given in the GitHub repository^7^ and used the provided pretrained weights and scripts for sampling. For unconditional generation, we used the configuration under examples/sampling/00 unconditional.yaml and provided the performance across 5 different versions. A summary of these 5 different versions is provided in the table below.

#### Genie 2

We followed instructions from the Github repository^8^, with a default noise scale of 0.6 for unconditional generation.

#### Ambient

We followed the provided instructions in the Github repository^9^. Specifically, for protein shorter than 300 residues, we use the model version ambient short checkpointed at epoch 130 and sample with a noise scale of 0.55. For proteins longer than 300 residues (inclusive), we use ambient long checkpointed at epoch 19 and sample with a noise scale of 0.6.

### B.3. Performance of short monomer generation

Unconditional short monomer generation (with sequence lengths fewer than 300) is an important first-step for validation since the majority of downstream design tasks require generation of short monomers for reasons such as easier expression and cost of screening. The ability to model the structural space of short monomers provides a better foundation for downstream design applications. Table 3 compares Genie 3 with other methods on unconditional short monomer generation using the set of metrics reported in Appendix B.1. For each method in consideration, we sample 100 structures per length between 50 and 250 (inclusive) with an increment of 50 in length. We observe that Genie 3 is capable of generating designable, diverse and novel structures, providing a foundation for downstream protein design tasks such as motif scaffolding and binder design. Genie 3 achieves better diversity and novelty than existing non-Genie-based protein diffusion models such as RFDiffusion and Proteina and its performance is on par with Genie 2. Genie 3 performs slightly behind Ambient on short monomer generation, suggesting the importance of better training data clustering, increasing model complexity and better handling of low confidence structures.

**Table 1.** Input feature dictionary to the model, where *N* is the number of tokens. Note that a token in our case corresponds to an atom, either C_*α*_ atom or a side chain heavy atom.

| Feature | Shape | Description |
| --- | --- | --- |
| residue_index | $[N]$ | Residue number in the input chain, renumbered per chain to start from 1. |
| token_index | $[N]$ | Token number which starts from 0, increases monotonically and does not restart for new chains. |
| token_mask | $[N]$ | Token mask indicating padding. |
| asym_id | $[N]$ | Unique integer for each distinct chain. |
| entity_id | $[N]$ | Unique integer for each distinct sequence. |
| sym_id | $[N]$ | Unique integer for each chain within the sequence. |
| is_atomized | $[N]$ | Mask indicating the token comes from an atomized residue, which is either a motif or an interface residue. |
| gt_restype | $[N, 20]$ | One-hot encoding of groundtruth sequence, where unknown residue is assigned with zeros. |
| atom_type | $[N, 37]$ | One-hot encoding of atom type in atom37 definition. |
| atom_symbol | $[N, 4]$ | One-hot encoding of atom symbol (N, C, O, S). |
| gt_atom_mask | $[N]$ | Mask indicating whether groundtruth atom position is available. |
| gt_atom_position | $[N, 3]$ | Groundtruth atom positions, initialized to zeros for invalid positions. |
| plddt | $[N]$ | Per-residue pLDDT for AF2-predicted structures; assigned with ones for experimentally validated structures such as PDB and Pinder structures. |
| token_frame_lindex | $[N]$ | Token index for left atom used for Frenet-Serret frame construction before assigning adjacent frames for terminal $C_\alpha$ and sidechain atoms (-1 if left atom is missing). |
| token_frame_rindex | $[N]$ | Token index for right atom used for Frenet-Serret frame construction before assigning adjacent frames for terminal $C_\alpha$ and sidechain atoms (-1 if right atom is missing). |
| token_frame_lindex_adj | $[N]$ | Token index for left atom used for Frenet-Serret frame construction after assigning adjacent frames for terminal $C_\alpha$ and sidechain atoms. |
| token_frame_cindex_adj | $[N]$ | Token index for center atom used for Frenet-Serret frame construction after assigning adjacent frames for terminal $C_\alpha$ and sidechain atoms. |
| token_frame_rindex_adj | $[N]$ | Token index for right atom used for Frenet-Serret frame construction after assigning adjacent frames for terminal $C_\alpha$ and sidechain atoms. |
| token_struct_mask | $[N]$ | Mask indicating whether the token has valid atom positions (equivalent to gt_atom_mask). |
| token_struct_frame_mask | $[N]$ | Mask indicating whether the token has valid frames after assigning adjacent frames for terminal $C_\alpha$ and sidechain atoms. |
| cond_seq_mask | $[N]$ | Mask indicating whether amino acid type is provided for the token as condition. |
| cond_group | $[N]$ | Unique integer for each conditional group, where inter-group positions and orientations are preserved as conditions while intra-group are not. |
| cond_struct_mask | $[N]$ | Mask indicating whether atom position is provided for the token as condition. |
| cond_struct_frame_mask | $[N]$ | Mask indicating whether frame information is provided for the token as condition. |
| cond_interface_mask | $[N]$ | Mask indicating whether the token is involved in the binding interface (for binder design). |

**Table 2.** Summary of different Protpardelle-1c versions.

| Version | Epoch | Description |
| --- | --- | --- |
| cc58 | 416 | single chain model, backbone-only |
| cc83 | 2616 | binder generation |
| cc91 | 383 | single chain model, all-atom |
| cc94 | 3100 | multichain model, all-atom |
| cc95 | 3490 | multichain model, backbone-only |

**Table 3.** Comparison of unconditional short monomer generation. For each method, we sample 100 structures per length between 50 and 250 (inclusive) with an increment of 50 in length, totaling 500 samples. To reduce sampling variance, we repeat each evaluation run 3 times and report the average metrics across the runs. Details on each evaluation run are included in Table 4 and Table 5, showing limited variations in metrics across runs.

| Method | Version | Designability | Diversity |  | Novelty |  | Sec. Struct. % |  |
| --- | --- | --- | --- | --- | --- | --- | --- | --- |
| | | | TM < 0.5 | TM < 0.6 | PDB | AFDB | $\alpha$ | $\beta$ |
| RFDiffusion |  | 0.93 | 0.42 | 0.69 | 0.36 | 0.35 | 61.9 | 17.9 |
| FrameFlow |  | 0.90 | 0.50 | 0.70 | 0.34 | 0.33 | 53.2 | 19.5 |
| Proteus |  | 0.93 | 0.24 | 0.40 | 0.37 | 0.37 | 71.1 | 9.0 |
| FoldFlow 2 | base | 0.97 | 0.43 | 0.77 | 0.34 | 0.32 | 83.7 | 0.8 |
|  | reft | 0.83 | 0.44 | 0.59 | 0.36 | 0.36 | 43.4 | 26.1 |
| Proteina | $\mathcal{M}_{FS}$ | 0.92 | 0.58 | 0.78 | 0.32 | 0.32 | 65.5 | 9.8 |
| La-Proteina | LD1 | 0.99 | 0.44 | 0.69 | 0.30 | 0.30 | 61.8 | 11.4 |
|  | LD2 | 0.99 | 0.29 | 0.50 | 0.27 | 0.27 | 62.4 | 11.0 |
| Protpardelle-1c | cc58 | 1.00 | 0.14 | 0.25 | 0.23 | 0.23 | 45.9 | 23.9 |
|  | cc94 | 0.97 | 0.13 | 0.34 | 0.28 | 0.28 | 68.1 | 9.1 |
| Genie 2 |  | 0.94 | 0.62 | 0.89 | 0.39 | 0.37 | 68.2 | 6.5 |
| Ambient | short | 0.97 | 0.76 | 0.93 | 0.41 | 0.39 | 76.7 | 2.2 |
| Genie 3 |  | 0.97 | 0.69 | 0.85 | 0.37 | 0.36 | 73.9 | 6.6 |

**Table 4.**
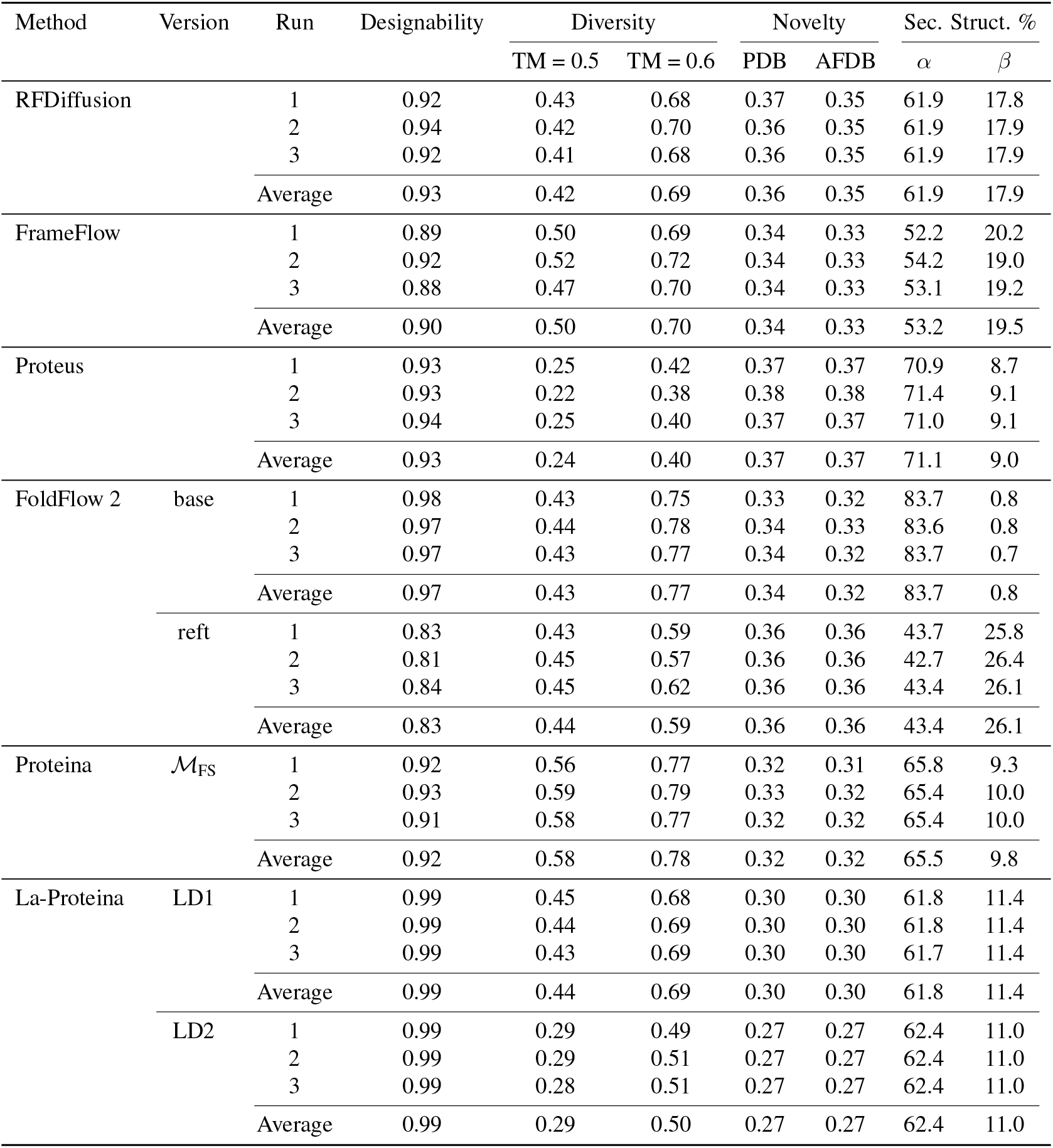
Unconditional generation performance on short monomers across runs (Part 1), which consists of 100 samples per length between 50 and 250 with an increment in length of 50 (500 structures in total).

| Method | Version | Run | Designability | Diversity |  | Novelty |  | Sec. Struct. % |  |
| --- | --- | --- | --- | --- | --- | --- | --- | --- | --- |
| | | | | TM = 0.5 | TM = 0.6 | PDB | AFDB | $\alpha$ | $\beta$ |
| RFDiffusion |  | 1 | 0.92 | 0.43 | 0.68 | 0.37 | 0.35 | 61.9 | 17.8 |
|  |  | 2 | 0.94 | 0.42 | 0.70 | 0.36 | 0.35 | 61.9 | 17.9 |
|  |  | 3 | 0.92 | 0.41 | 0.68 | 0.36 | 0.35 | 61.9 | 17.9 |
|  |  | Average | 0.93 | 0.42 | 0.69 | 0.36 | 0.35 | 61.9 | 17.9 |
| FrameFlow |  | 1 | 0.89 | 0.50 | 0.69 | 0.34 | 0.33 | 52.2 | 20.2 |
|  |  | 2 | 0.92 | 0.52 | 0.72 | 0.34 | 0.33 | 54.2 | 19.0 |
|  |  | 3 | 0.88 | 0.47 | 0.70 | 0.34 | 0.33 | 53.1 | 19.2 |
|  |  | Average | 0.90 | 0.50 | 0.70 | 0.34 | 0.33 | 53.2 | 19.5 |
| Proteus |  | 1 | 0.93 | 0.25 | 0.42 | 0.37 | 0.37 | 70.9 | 8.7 |
|  |  | 2 | 0.93 | 0.22 | 0.38 | 0.38 | 0.38 | 71.4 | 9.1 |
|  |  | 3 | 0.94 | 0.25 | 0.40 | 0.37 | 0.37 | 71.0 | 9.1 |
|  |  | Average | 0.93 | 0.24 | 0.40 | 0.37 | 0.37 | 71.1 | 9.0 |
| FoldFlow 2 | base | 1 | 0.98 | 0.43 | 0.75 | 0.33 | 0.32 | 83.7 | 0.8 |
|  |  | 2 | 0.97 | 0.44 | 0.78 | 0.34 | 0.33 | 83.6 | 0.8 |
|  |  | 3 | 0.97 | 0.43 | 0.77 | 0.34 | 0.32 | 83.7 | 0.7 |
|  |  | Average | 0.97 | 0.43 | 0.77 | 0.34 | 0.32 | 83.7 | 0.8 |
|  | reft | 1 | 0.83 | 0.43 | 0.59 | 0.36 | 0.36 | 43.7 | 25.8 |
|  |  | 2 | 0.81 | 0.45 | 0.57 | 0.36 | 0.36 | 42.7 | 26.4 |
|  |  | 3 | 0.84 | 0.45 | 0.62 | 0.36 | 0.36 | 43.4 | 26.1 |
|  |  | Average | 0.83 | 0.44 | 0.59 | 0.36 | 0.36 | 43.4 | 26.1 |
| Proteina | $\mathcal{M}_{FS}$ | 1 | 0.92 | 0.56 | 0.77 | 0.32 | 0.31 | 65.8 | 9.3 |
|  |  | 2 | 0.93 | 0.59 | 0.79 | 0.33 | 0.32 | 65.4 | 10.0 |
|  |  | 3 | 0.91 | 0.58 | 0.77 | 0.32 | 0.32 | 65.4 | 10.0 |
|  |  | Average | 0.92 | 0.58 | 0.78 | 0.32 | 0.32 | 65.5 | 9.8 |
| La-Proteina | LD1 | 1 | 0.99 | 0.45 | 0.68 | 0.30 | 0.30 | 61.8 | 11.4 |
|  |  | 2 | 0.99 | 0.44 | 0.69 | 0.30 | 0.30 | 61.8 | 11.4 |
|  |  | 3 | 0.99 | 0.43 | 0.69 | 0.30 | 0.30 | 61.7 | 11.4 |
|  |  | Average | 0.99 | 0.44 | 0.69 | 0.30 | 0.30 | 61.8 | 11.4 |
|  | LD2 | 1 | 0.99 | 0.29 | 0.49 | 0.27 | 0.27 | 62.4 | 11.0 |
|  |  | 2 | 0.99 | 0.29 | 0.51 | 0.27 | 0.27 | 62.4 | 11.0 |
|  |  | 3 | 0.99 | 0.28 | 0.51 | 0.27 | 0.27 | 62.4 | 11.0 |
|  |  | Average | 0.99 | 0.29 | 0.50 | 0.27 | 0.27 | 62.4 | 11.0 |

**Table 5.** Unconditional generation performance on short monomers across runs (Part 2), which consists of 100 samples per length between 50 and 250 with an increment in length of 50 (500 structures in total).

| Method | Version | Run | Designability | Diversity |  | Novelty |  | Sec. Struct. % |  |
| --- | --- | --- | --- | --- | --- | --- | --- | --- | --- |
| | | | | TM = 0.5 | TM = 0.6 | PDB | AFDB | $\alpha$ | $\beta$ |
| Protpardelle-1c | cc58 | 1 | 1.00 | 0.13 | 0.24 | 0.23 | 0.23 | 45.3 | 24.3 |
|  |  | 2 | 0.99 | 0.16 | 0.27 | 0.23 | 0.23 | 46.1 | 23.8 |
|  |  | 3 | 1.00 | 0.12 | 0.25 | 0.23 | 0.23 | 46.2 | 23.5 |
|  |  | Average | 1.00 | 0.14 | 0.25 | 0.23 | 0.23 | 45.9 | 23.9 |
|  | cc83 | 1 | 1.00 | 0.11 | 0.21 | 0.25 | 0.26 | 56.6 | 18.8 |
|  |  | 2 | 0.99 | 0.12 | 0.23 | 0.25 | 0.26 | 57.2 | 18.2 |
|  |  | 3 | 1.00 | 0.11 | 0.23 | 0.24 | 0.25 | 57.8 | 17.9 |
|  |  | Average | 1.00 | 0.11 | 0.22 | 0.25 | 0.26 | 57.2 | 18.3 |
|  | cc91 | 1 | 0.99 | 0.07 | 0.18 | 0.24 | 0.24 | 62.3 | 12.8 |
|  |  | 2 | 0.99 | 0.09 | 0.21 | 0.25 | 0.25 | 61.2 | 13.8 |
|  |  | 3 | 0.98 | 0.10 | 0.21 | 0.25 | 0.25 | 61.7 | 13.1 |
|  |  | Average | 0.99 | 0.09 | 0.20 | 0.25 | 0.25 | 61.7 | 13.2 |
|  | cc94 | 1 | 0.96 | 0.13 | 0.33 | 0.28 | 0.28 | 67.5 | 9.5 |
|  |  | 2 | 0.97 | 0.13 | 0.34 | 0.28 | 0.28 | 69.0 | 8.5 |
|  |  | 3 | 0.97 | 0.13 | 0.34 | 0.28 | 0.27 | 67.9 | 9.4 |
|  |  | Average | 0.97 | 0.13 | 0.34 | 0.28 | 0.28 | 68.1 | 9.1 |
|  | cc95 | 1 | 0.99 | 0.15 | 0.25 | 0.25 | 0.26 | 47.9 | 24.9 |
|  |  | 2 | 0.99 | 0.16 | 0.25 | 0.25 | 0.26 | 48.5 | 24.7 |
|  |  | 3 | 0.99 | 0.14 | 0.26 | 0.25 | 0.26 | 48.1 | 25.0 |
|  |  | Average | 0.99 | 0.15 | 0.25 | 0.25 | 0.26 | 48.2 | 24.9 |
| Genie 2 |  | 1 | 0.94 | 0.61 | 0.89 | 0.39 | 0.37 | 67.3 | 7.1 |
|  |  | 2 | 0.95 | 0.60 | 0.90 | 0.38 | 0.37 | 68.7 | 6.2 |
|  |  | 3 | 0.94 | 0.66 | 0.88 | 0.39 | 0.38 | 68.5 | 6.2 |
|  |  | Average | 0.94 | 0.62 | 0.89 | 0.39 | 0.37 | 68.2 | 6.5 |
| Ambient | short | 1 | 0.96 | 0.74 | 0.91 | 0.41 | 0.39 | 77.4 | 1.8 |
|  |  | 2 | 0.97 | 0.74 | 0.93 | 0.41 | 0.39 | 76.7 | 2.1 |
|  |  | 3 | 0.98 | 0.81 | 0.95 | 0.41 | 0.39 | 74.9 | 2.7 |
|  |  | Average | 0.97 | 0.76 | 0.93 | 0.41 | 0.39 | 76.7 | 2.2 |
| Genie 3 |  | 1 | 0.96 | 0.69 | 0.86 | 0.37 | 0.36 | 73.5 | 6.8 |
|  |  | 2 | 0.97 | 0.71 | 0.85 | 0.37 | 0.36 | 73.7 | 6.6 |
|  |  | 3 | 0.97 | 0.67 | 0.84 | 0.37 | 0.36 | 74.4 | 6.4 |
|  |  | Average | 0.97 | 0.69 | 0.85 | 0.37 | 0.36 | 73.9 | 6.6 |

### B.4. Additional details on long monomer generations

### B.5. Sampling time

We compare the sampling times of different models on unconditional generation. For fair comparison, we profile sampling time using a single A100 GPU (40GB memory) and average the inference time of a single sample over 10 runs.

## C. Motif Scaffolding

### C.1 Evaluation metrics

We use the same designability metric described in Section B.1 with an additional constraint-satisfaction metric to ensure the successful integration of the desired motif. Specifically, we consider a generated structure as a (backbone) success if it satisfies scRMSD *<* 2Å and motif backbone RMSD *<* 1Å, which is consistent with existing literature (Zheng et al., 2025; Lin et al., 2024). Here, we consider all N, Ca, C and O atoms for motif backbone RMSD computation and motif alignment is computed using the Kabsch algorithm. A generated structure is an all-atom success if it additionally passes motif all-atom RMSD *<* 2Å. Here, we consider all heavy atoms for motif all-atom RMSD computation. Motif alignment is also computed using the Kabsch algorithm. We also used the same FoldSeek (Van Kempen et al., 2024) clustering command described in Section B.1 to compute the number of unique successes, using a TM threshold of 0.6. Success rate is computed as the percentage of successful generated structures and unique success rate is computed as the percentage of successful generated clusters.

### C.2 Reproduction of baseline methods

We provide details on the reproduction of baseline methods on motif scaffolding below.

#### RFDiffusion

We followed instructions from the Github repository^10^ using the pretrained weights located at http://files.ipd.uw.edu/pub/RFdiffusion/. We use the provided default configuration, which sets the noise scale to 1.

#### FrameFlow

We followed instructions from the Github repository^11^ and used the provided pretrained weights for motif scaffolding, located at checkpoint weights/pdb amortization/published.ckpt for sampling.

#### Genie 2

We followed instructions from the Github repository^12^ and used the default noise scale of 0.4 for motif scaffolding.

#### Proteina

We followed the instructions given in the Github repository^13^ for sampling. We used the motif scaffolding model *M*_motif_ with a noise scale of 0.5.

#### La-Proteina

We followed the instructions given in the Github repository^14^ and used provided pretrained weights for motif scaffolding. We used the checkpoint of LD4 (indexed atomistic motif scaffolding) with noise scales of 0.1 for both the alpha carbon atoms and for the latent variables, combined with the AE3 checkpoint for the autoencoder.

#### Protpardelle-1c

We followed the instructions given in the GitHub repository^15^ and used the provided pretrained weights and scripts for sampling. For motif scaffolding, we used the configuration under examples/sampling/03 motifbench.yaml, which uses the backbone model (cc58) with side chain conditioning (see Table 2).

### C.3. Discussion on motif scaffolding performance

We used a recently published motif scaffolding benchmark named MotifBench consisting of 30 problems. This problem set builds upon the original set of motif scaffolding problems curated by Watson et al. (2023) and introduces more challenging motif scaffolding tasks to better assess the performance of protein generative models. Figure 2B compares the performance of Genie 3 and other recent protein diffusion models, demonstrating that Genie 3 is capable of solving more problems with greater diversity. One caveat is the difference between indexed and unindexed generation in the context of motif scaffolding: for indexed generation, the sequence index of the motif is fixed and provided to the model as a condition. We consider indexed generation in this work. We note that the selection of motif placements (in other words, the number of residues in between motif segments, as well as the sequential order of motif segments) can significantly affect model performance. To ensure consistent evaluations, we use the default motif configuration provided in MotifBench GitHub repository^16^ (under examples/contig specifications.csv) when evaluating.

To better understand the performance of Genie 3, we further compare the individual task performance of Genie 3 with other models in Figures 7 and 8. We highlight problems that Genie 3 performs better on in blue, problems that competing model performs better on in orange, and problems that both models achieve the same performance on in grey. For each problem-model pair, we sample and evaluate 100 designs per run and report the average across three runs. Detailed statistics for each run are provided in Table 8. For Genie 3, we also test with different directional scales and select *θ* = 0.1 since it gives the best performance (Figure 6A). Detailed statistics for runs with different directional scales are provided in Table 9. When comparing to leading methods Protpardelle-1c and La-Proteina, we observe that Genie 3 generates more diverse clusters than Protpardelle-1c for Group 1 and Group 2 (motifs with 1-2 segments), but lags in performance for Group 3 problems (motifs with at least 3 segments).

**Table 6.** Unconditional generation performance on long monomers. For each length between 300 and 800 (with a length step of 100), we sample 100 structures from each model and compare their performance based on diversity, clustered at a TM threshold of 0.5.

| Method | Version | Run | Designability |  |  |  |  |  | Diversity (TM = 0.5) |  |  |  |  |  |
| --- | --- | --- | --- | --- | --- | --- | --- | --- | --- | --- | --- | --- | --- | --- |
|  |  |  | L = 300 | 400 | 500 | 600 | 700 | 800 | L = 300 | 400 | 500 | 600 | 700 | 800 |
| RFDiffusion |  | 1 | 0.74 | 0.50 | 0.32 | 0.13 | 0.04 | 0.03 | 0.49 | 0.44 | 0.32 | 0.13 | 0.04 | 0.03 |
|  |  | 2 | 0.68 | 0.57 | 0.32 | 0.15 | 0.10 | 0.01 | 0.41 | 0.48 | 0.27 | 0.13 | 0.10 | 0.01 |
|  |  | 3 | 0.79 | 0.55 | 0.33 | 0.13 | 0.03 | 0.02 | 0.54 | 0.47 | 0.28 | 0.13 | 0.03 | 0.02 |
|  |  | Average | 0.74 | 0.54 | 0.32 | 0.14 | 0.06 | 0.02 | 0.48 | 0.46 | 0.29 | 0.13 | 0.06 | 0.02 |
| FrameFlow |  | 1 | 0.70 | 0.61 | 0.37 | 0.01 | 0.00 | 0.00 | 0.46 | 0.51 | 0.36 | 0.01 | 0.00 | 0.00 |
|  |  | 2 | 0.74 | 0.67 | 0.42 | 0.04 | 0.00 | 0.00 | 0.61 | 0.57 | 0.38 | 0.04 | 0.00 | 0.00 |
|  |  | 3 | 0.71 | 0.64 | 0.42 | 0.04 | 0.00 | 0.00 | 0.60 | 0.61 | 0.41 | 0.04 | 0.00 | 0.00 |
|  |  | Average | 0.72 | 0.64 | 0.4 | 0.03 | 0.00 | 0.00 | 0.56 | 0.56 | 0.38 | 0.03 | 0.00 | 0.00 |
| Proteus |  | 1 | 0.93 | 0.80 | 0.76 | 0.65 | 0.44 | 0.28 | 0.21 | 0.30 | 0.31 | 0.21 | 0.13 | 0.08 |
|  |  | 2 | 0.92 | 0.90 | 0.78 | 0.65 | 0.42 | 0.27 | 0.22 | 0.37 | 0.27 | 0.18 | 0.07 | 0.07 |
|  |  | 3 | 0.94 | 0.88 | 0.84 | 0.54 | 0.38 | 0.23 | 0.20 | 0.40 | 0.36 | 0.17 | 0.08 | 0.05 |
|  |  | Average | 0.93 | 0.86 | 0.79 | 0.61 | 0.41 | 0.26 | 0.21 | 0.36 | 0.31 | 0.19 | 0.09 | 0.07 |
| Proteina | $\mathcal{M}_{\text{long}}$ | 1 | 0.93 | 0.86 | 0.84 | 0.74 | 0.74 | 0.46 | 0.46 | 0.44 | 0.54 | 0.58 | 0.60 | 0.41 |
|  |  | 2 | 0.94 | 0.87 | 0.77 | 0.77 | 0.73 | 0.54 | 0.48 | 0.43 | 0.50 | 0.60 | 0.58 | 0.44 |
|  |  | 3 | 0.93 | 0.88 | 0.75 | 0.78 | 0.74 | 0.56 | 0.48 | 0.47 | 0.49 | 0.58 | 0.59 | 0.49 |
|  |  | Average | 0.93 | 0.87 | 0.79 | 0.76 | 0.74 | 0.52 | 0.47 | 0.45 | 0.51 | 0.59 | 0.59 | 0.45 |
| La-Proteina | LD3 | 1 | 0.90 | 0.81 | 0.60 | 0.50 | 0.34 | 0.25 | 0.22 | 0.25 | 0.26 | 0.29 | 0.26 | 0.22 |
|  |  | 2 | 0.90 | 0.82 | 0.59 | 0.51 | 0.27 | 0.24 | 0.22 | 0.26 | 0.24 | 0.29 | 0.18 | 0.22 |
|  |  | 3 | 0.91 | 0.79 | 0.59 | 0.51 | 0.29 | 0.23 | 0.22 | 0.26 | 0.25 | 0.31 | 0.19 | 0.21 |
|  |  | Average | 0.9 | 0.81 | 0.59 | 0.51 | 0.3 | 0.24 | 0.22 | 0.26 | 0.25 | 0.30 | 0.21 | 0.22 |
| Genie 2 |  | 1 | 0.87 | 0.75 | 0.50 | 0.13 | 0.02 | 0.00 | 0.82 | 0.72 | 0.50 | 0.13 | 0.02 | 0.00 |
|  |  | 2 | 0.90 | 0.77 | 0.55 | 0.17 | 0.00 | 0.00 | 0.82 | 0.74 | 0.55 | 0.17 | 0.00 | 0.00 |
|  |  | 3 | 0.90 | 0.69 | 0.49 | 0.14 | 0.01 | 0.00 | 0.82 | 0.68 | 0.49 | 0.14 | 0.01 | 0.00 |
|  |  | Average | 0.89 | 0.74 | 0.51 | 0.15 | 0.01 | 0.00 | 0.82 | 0.71 | 0.51 | 0.15 | 0.01 | 0.00 |
| Ambient | long | 1 | 0.96 | 0.91 | 0.93 | 0.98 | 0.85 | 0.68 | 0.95 | 0.90 | 0.92 | 0.98 | 0.85 | 0.68 |
|  |  | 2 | 0.94 | 0.95 | 0.94 | 0.93 | 0.88 | 0.68 | 0.94 | 0.95 | 0.94 | 0.92 | 0.88 | 0.68 |
|  |  | 3 | 0.97 | 0.93 | 0.95 | 0.87 | 0.89 | 0.70 | 0.96 | 0.93 | 0.95 | 0.86 | 0.89 | 0.70 |
|  |  | Average | 0.96 | 0.93 | 0.94 | 0.93 | 0.87 | 0.69 | 0.95 | 0.93 | 0.94 | 0.92 | 0.87 | 0.69 |
| Genie 3 |  | 1 | 0.92 | 1.00 | 1.00 | 1.00 | 0.94 | 0.77 | 0.92 | 1.00 | 1.00 | 1.00 | 0.94 | 0.77 |
|  |  | 2 | 0.94 | 1.00 | 1.00 | 0.97 | 0.92 | 0.83 | 0.94 | 1.00 | 1.00 | 0.97 | 0.92 | 0.83 |
|  |  | 3 | 0.93 | 1.00 | 1.00 | 0.97 | 0.93 | 0.88 | 0.93 | 0.99 | 1.00 | 0.97 | 0.93 | 0.88 |
|  |  | Average | 0.93 | 1.00 | 1.00 | 0.98 | 0.93 | 0.83 | 0.93 | 1.00 | 1.00 | 0.98 | 0.93 | 0.83 |

**Table 7.**
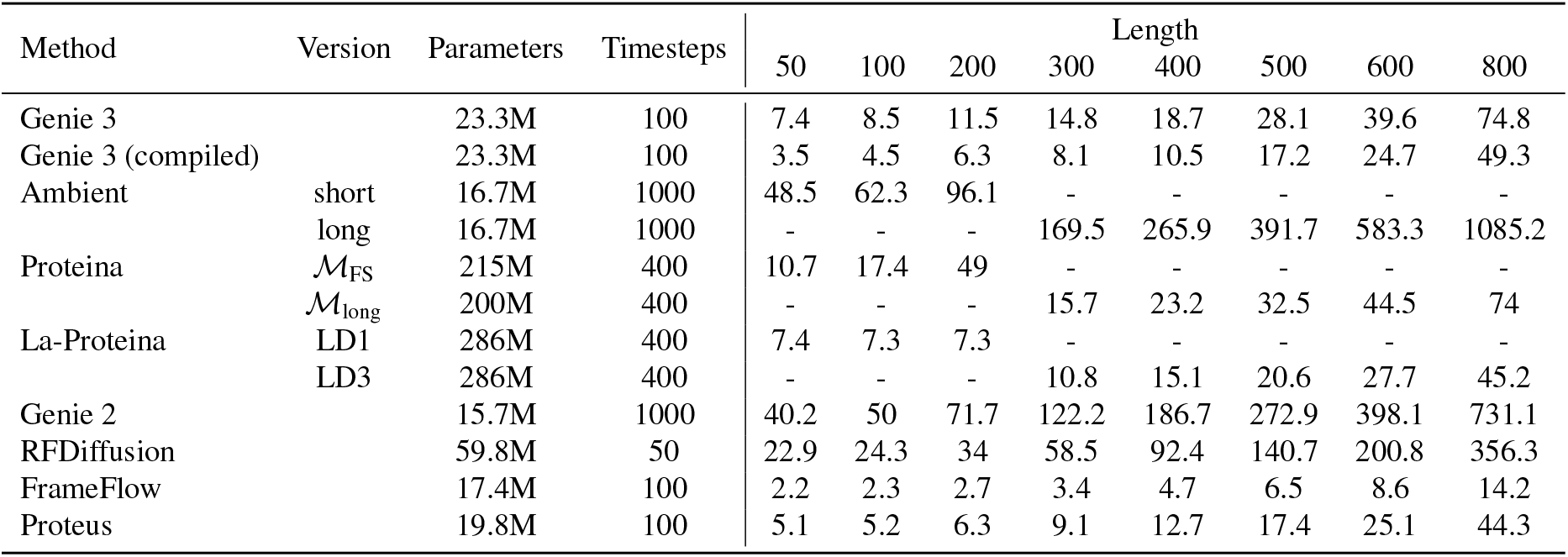
Sampling time of different methods for proteins of different lengths.

**Table 8.**
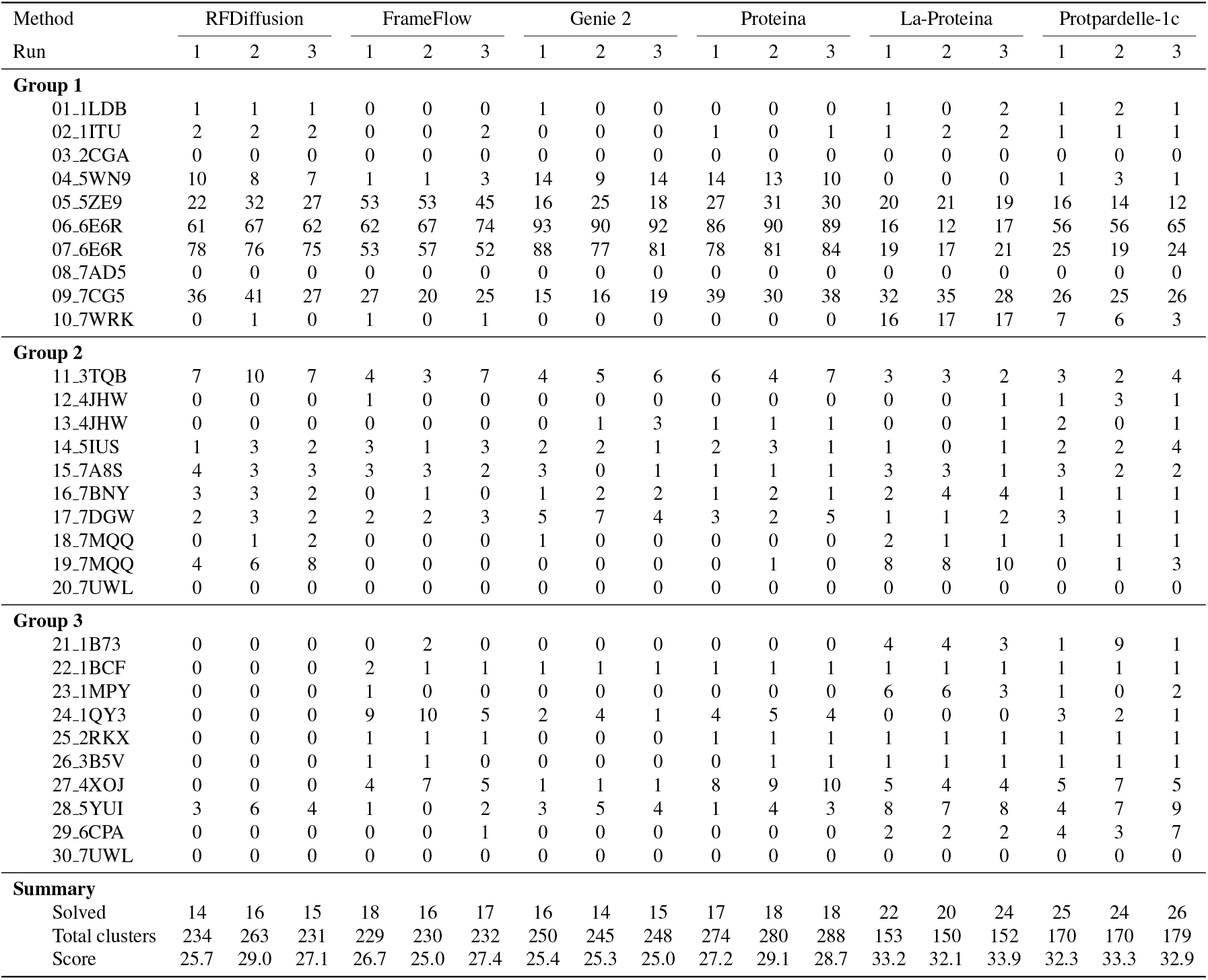
Performance of different protein diffusion models on MotifBench, measured in diversity clustered at TM = 0.6. We sample 100 structures per problem per model and repeat 3 times per model to ensure statistical significance.

| Method | RFDiffusion |  |  | FrameFlow |  |  | Genie 2 |  |  | Proteina |  |  | La-Proteina |  |  | Protpardelle-1c |  |  |
| --- | --- | --- | --- | --- | --- | --- | --- | --- | --- | --- | --- | --- | --- | --- | --- | --- | --- | --- |
| Run | 1 | 2 | 3 | 1 | 2 | 3 | 1 | 2 | 3 | 1 | 2 | 3 | 1 | 2 | 3 | 1 | 2 | 3 |
| **Group 1** |  |  |  |  |  |  |  |  |  |  |  |  |  |  |  |  |  |  |
| 01_1LDB | 1 | 1 | 1 | 0 | 0 | 0 | 1 | 0 | 0 | 0 | 0 | 0 | 1 | 0 | 2 | 1 | 2 | 1 |
| 02_1ITU | 2 | 2 | 2 | 0 | 0 | 2 | 0 | 0 | 0 | 1 | 0 | 1 | 1 | 2 | 2 | 1 | 1 | 1 |
| 03_2CGA | 0 | 0 | 0 | 0 | 0 | 0 | 0 | 0 | 0 | 0 | 0 | 0 | 0 | 0 | 0 | 0 | 0 | 0 |
| 04_5WN9 | 10 | 8 | 7 | 1 | 1 | 3 | 14 | 9 | 14 | 14 | 13 | 10 | 0 | 0 | 0 | 1 | 3 | 1 |
| 05_5ZE9 | 22 | 32 | 27 | 53 | 53 | 45 | 16 | 25 | 18 | 27 | 31 | 30 | 20 | 21 | 19 | 16 | 14 | 12 |
| 06_6E6R | 61 | 67 | 62 | 62 | 67 | 74 | 93 | 90 | 92 | 86 | 90 | 89 | 16 | 12 | 17 | 56 | 56 | 65 |
| 07_6E6R | 78 | 76 | 75 | 53 | 57 | 52 | 88 | 77 | 81 | 78 | 81 | 84 | 19 | 17 | 21 | 25 | 19 | 24 |
| 08_7AD5 | 0 | 0 | 0 | 0 | 0 | 0 | 0 | 0 | 0 | 0 | 0 | 0 | 0 | 0 | 0 | 0 | 0 | 0 |
| 09_7CG5 | 36 | 41 | 27 | 27 | 20 | 25 | 15 | 16 | 19 | 39 | 30 | 38 | 32 | 35 | 28 | 26 | 25 | 26 |
| 10_7WRK | 0 | 1 | 0 | 1 | 0 | 1 | 0 | 0 | 0 | 0 | 0 | 0 | 16 | 17 | 17 | 7 | 6 | 3 |
| **Group 2** |  |  |  |  |  |  |  |  |  |  |  |  |  |  |  |  |  |  |
| 11_3TQB | 7 | 10 | 7 | 4 | 3 | 7 | 4 | 5 | 6 | 6 | 4 | 7 | 3 | 3 | 2 | 3 | 2 | 4 |
| 12_4JHW | 0 | 0 | 0 | 1 | 0 | 0 | 0 | 0 | 0 | 0 | 0 | 0 | 0 | 0 | 1 | 1 | 3 | 1 |
| 13_4JHW | 0 | 0 | 0 | 0 | 0 | 0 | 0 | 1 | 3 | 1 | 1 | 1 | 0 | 0 | 1 | 2 | 0 | 1 |
| 14_5IUS | 1 | 3 | 2 | 3 | 1 | 3 | 2 | 2 | 1 | 2 | 3 | 1 | 1 | 0 | 1 | 2 | 2 | 4 |
| 15_7A8S | 4 | 3 | 3 | 3 | 3 | 2 | 3 | 0 | 1 | 1 | 1 | 1 | 3 | 3 | 1 | 3 | 2 | 2 |
| 16_7BNY | 3 | 3 | 2 | 0 | 1 | 0 | 1 | 2 | 2 | 1 | 2 | 1 | 2 | 4 | 4 | 1 | 1 | 1 |
| 17_7DGW | 2 | 3 | 2 | 2 | 2 | 3 | 5 | 7 | 4 | 3 | 2 | 5 | 1 | 1 | 2 | 3 | 1 | 1 |
| 18_7MQQ | 0 | 1 | 2 | 0 | 0 | 0 | 1 | 0 | 0 | 0 | 0 | 0 | 2 | 1 | 1 | 1 | 1 | 1 |
| 19_7MQQ | 4 | 6 | 8 | 0 | 0 | 0 | 0 | 0 | 0 | 0 | 1 | 0 | 8 | 8 | 10 | 0 | 1 | 3 |
| 20_7UWL | 0 | 0 | 0 | 0 | 0 | 0 | 0 | 0 | 0 | 0 | 0 | 0 | 0 | 0 | 0 | 0 | 0 | 0 |
| **Group 3** |  |  |  |  |  |  |  |  |  |  |  |  |  |  |  |  |  |  |
| 21_1B73 | 0 | 0 | 0 | 0 | 2 | 0 | 0 | 0 | 0 | 0 | 0 | 0 | 4 | 4 | 3 | 1 | 9 | 1 |
| 22_1BCF | 0 | 0 | 0 | 2 | 1 | 1 | 1 | 1 | 1 | 1 | 1 | 1 | 1 | 1 | 1 | 1 | 1 | 1 |
| 23_1MPY | 0 | 0 | 0 | 1 | 0 | 0 | 0 | 0 | 0 | 0 | 0 | 0 | 6 | 6 | 3 | 1 | 0 | 2 |
| 24_1QY3 | 0 | 0 | 0 | 9 | 10 | 5 | 2 | 4 | 1 | 4 | 5 | 4 | 0 | 0 | 0 | 3 | 2 | 1 |
| 25_2RKX | 0 | 0 | 0 | 1 | 1 | 1 | 0 | 0 | 0 | 1 | 1 | 1 | 1 | 1 | 1 | 1 | 1 | 1 |
| 26_3B5V | 0 | 0 | 0 | 1 | 1 | 0 | 0 | 0 | 0 | 0 | 1 | 1 | 1 | 1 | 1 | 1 | 1 | 1 |
| 27_4XOJ | 0 | 0 | 0 | 4 | 7 | 5 | 1 | 1 | 1 | 8 | 9 | 10 | 5 | 4 | 4 | 5 | 7 | 5 |
| 28_5YUI | 3 | 6 | 4 | 1 | 0 | 2 | 3 | 5 | 4 | 1 | 4 | 3 | 8 | 7 | 8 | 4 | 7 | 9 |
| 29_6CPA | 0 | 0 | 0 | 0 | 0 | 1 | 0 | 0 | 0 | 0 | 0 | 0 | 2 | 2 | 2 | 4 | 3 | 7 |
| 30_7UWL | 0 | 0 | 0 | 0 | 0 | 0 | 0 | 0 | 0 | 0 | 0 | 0 | 0 | 0 | 0 | 0 | 0 | 0 |
| **Summary** |  |  |  |  |  |  |  |  |  |  |  |  |  |  |  |  |  |  |
| Solved | 14 | 16 | 15 | 18 | 16 | 17 | 16 | 14 | 15 | 17 | 18 | 18 | 22 | 20 | 24 | 25 | 24 | 26 |
| Total clusters | 234 | 263 | 231 | 229 | 230 | 232 | 250 | 245 | 248 | 274 | 280 | 288 | 153 | 150 | 152 | 170 | 170 | 179 |
| Score | 25.7 | 29.0 | 27.1 | 26.7 | 25.0 | 27.4 | 25.4 | 25.3 | 25.0 | 27.2 | 29.1 | 28.7 | 33.2 | 32.1 | 33.9 | 32.3 | 33.3 | 32.9 |

**Table 9.** Performance of Genie 3 on MotifBench across different directional scales, measured in diversity clustered at TM = 0.6. We sample 100 structures per problem per directional scale and repeat 3 times for each directional scale to ensure statistical significance.

| Directional Scale | $\eta = 0.0$ | | | $\eta = 0.1$ | | | $\eta = 0.2$ | | | $\eta = 0.4$ | | | $\eta = 0.6$ | | | $\eta = 0.8$ | | |
| --- | --- | --- | --- | --- | --- | --- | --- | --- | --- | --- | --- | --- | --- | --- | --- | --- | --- | --- |
| Run | 1 | 2 | 3 | 1 | 2 | 3 | 1 | 2 | 3 | 1 | 2 | 3 | 1 | 2 | 3 | 1 | 2 | 3 |
| **Group 1** |  |  |  |  |  |  |  |  |  |  |  |  |  |  |  |  |  |  |
| 01_1LDB | 4 | 5 | 4 | 1 | 1 | 2 | 1 | 2 | 1 | 1 | 1 | 2 | 0 | 1 | 1 | 2 | 0 | 0 |
| 02_1ITU | 3 | 2 | 3 | 5 | 2 | 4 | 4 | 5 | 4 | 2 | 2 | 1 | 0 | 2 | 0 | 0 | 0 | 1 |
| 03_2CGA | 0 | 0 | 0 | 0 | 1 | 1 | 0 | 0 | 0 | 0 | 0 | 0 | 0 | 0 | 0 | 0 | 0 | 0 |
| 04_5WN9 | 21 | 16 | 21 | 22 | 31 | 20 | 18 | 18 | 14 | 21 | 16 | 19 | 14 | 18 | 22 | 7 | 8 | 13 |
| 05_5ZE9 | 17 | 23 | 19 | 14 | 18 | 17 | 17 | 19 | 21 | 22 | 13 | 18 | 21 | 22 | 22 | 22 | 18 | 15 |
| 06_6E6R | 69 | 75 | 72 | 74 | 74 | 74 | 74 | 82 | 82 | 73 | 72 | 72 | 82 | 74 | 76 | 78 | 78 | 72 |
| 07_6E6R | 98 | 93 | 97 | 95 | 95 | 95 | 96 | 99 | 98 | 92 | 95 | 96 | 92 | 88 | 93 | 83 | 81 | 88 |
| 08_7AD5 | 0 | 0 | 0 | 1 | 1 | 0 | 1 | 0 | 1 | 0 | 0 | 0 | 0 | 0 | 0 | 0 | 0 | 0 |
| 09_7CG5 | 69 | 61 | 68 | 66 | 71 | 67 | 74 | 69 | 67 | 62 | 70 | 67 | 66 | 66 | 56 | 44 | 47 | 43 |
| 10_7WRK | 3 | 4 | 2 | 1 | 2 | 2 | 5 | 3 | 0 | 0 | 4 | 4 | 0 | 0 | 4 | 0 | 0 | 0 |
| **Group 2** |  |  |  |  |  |  |  |  |  |  |  |  |  |  |  |  |  |  |
| 11_3TQB | 8 | 8 | 5 | 7 | 11 | 6 | 9 | 9 | 5 | 8 | 12 | 11 | 10 | 9 | 10 | 11 | 10 | 13 |
| 12_4JHW | 1 | 0 | 0 | 1 | 1 | 0 | 0 | 1 | 1 | 0 | 0 | 0 | 0 | 1 | 0 | 0 | 0 | 1 |
| 13_4JHW | 2 | 3 | 4 | 6 | 4 | 5 | 6 | 4 | 9 | 3 | 1 | 1 | 3 | 1 | 0 | 3 | 3 | 1 |
| 14_5IUS | 2 | 2 | 1 | 2 | 4 | 4 | 3 | 4 | 4 | 6 | 2 | 3 | 3 | 5 | 4 | 3 | 4 | 3 |
| 15_7A8S | 1 | 2 | 3 | 1 | 1 | 2 | 2 | 2 | 2 | 0 | 2 | 1 | 3 | 1 | 1 | 1 | 1 | 2 |
| 16_7BNY | 3 | 2 | 3 | 3 | 3 | 6 | 2 | 4 | 3 | 3 | 5 | 3 | 1 | 4 | 3 | 1 | 1 | 2 |
| 17_7DGW | 2 | 4 | 4 | 5 | 4 | 5 | 5 | 4 | 3 | 3 | 3 | 3 | 9 | 6 | 4 | 4 | 4 | 6 |
| 18_7MQQ | 1 | 2 | 1 | 1 | 1 | 2 | 0 | 1 | 1 | 0 | 0 | 1 | 2 | 0 | 0 | 1 | 1 | 1 |
| 19_7MQQ | 14 | 8 | 15 | 9 | 6 | 8 | 17 | 7 | 6 | 7 | 6 | 6 | 4 | 9 | 7 | 2 | 2 | 3 |
| 20_7UWL | 0 | 0 | 0 | 0 | 0 | 0 | 0 | 0 | 0 | 0 | 0 | 0 | 0 | 0 | 0 | 0 | 0 | 0 |
| **Group 3** |  |  |  |  |  |  |  |  |  |  |  |  |  |  |  |  |  |  |
| 21_1B73 | 1 | 2 | 0 | 1 | 1 | 2 | 0 | 3 | 1 | 0 | 2 | 1 | 1 | 1 | 1 | 1 | 3 | 0 |
| 22_1BCF | 1 | 1 | 1 | 1 | 1 | 1 | 1 | 1 | 1 | 1 | 1 | 1 | 1 | 1 | 1 | 1 | 1 | 1 |
| 23_1MPY | 0 | 0 | 0 | 0 | 0 | 0 | 0 | 0 | 0 | 0 | 0 | 0 | 0 | 0 | 0 | 0 | 0 | 0 |
| 24_1QY3 | 33 | 29 | 29 | 27 | 25 | 26 | 31 | 21 | 28 | 29 | 32 | 22 | 19 | 17 | 14 | 13 | 9 | 15 |
| 25_2RKX | 1 | 3 | 1 | 1 | 2 | 1 | 2 | 2 | 3 | 1 | 2 | 3 | 0 | 2 | 1 | 2 | 1 | 0 |
| 26_3B5V | 0 | 0 | 0 | 0 | 0 | 0 | 0 | 0 | 0 | 0 | 0 | 1 | 0 | 0 | 1 | 0 | 0 | 0 |
| 27_4XOJ | 0 | 2 | 1 | 1 | 1 | 0 | 0 | 1 | 2 | 0 | 1 | 2 | 1 | 1 | 2 | 1 | 0 | 0 |
| 28_5YUI | 5 | 8 | 8 | 5 | 3 | 6 | 10 | 5 | 7 | 7 | 9 | 10 | 9 | 7 | 4 | 3 | 7 | 2 |
| 29_6CPA | 2 | 2 | 1 | 2 | 1 | 2 | 1 | 5 | 3 | 3 | 2 | 1 | 0 | 0 | 3 | 0 | 3 | 1 |
| 30_7UWL | 0 | 0 | 0 | 0 | 0 | 0 | 0 | 0 | 0 | 0 | 0 | 0 | 0 | 0 | 0 | 0 | 0 | 0 |
| **Summary** |  |  |  |  |  |  |  |  |  |  |  |  |  |  |  |  |  |  |
| Solved | 23 | 23 | 22 | 25 | 26 | 23 | 21 | 24 | 24 | 18 | 22 | 24 | 18 | 21 | 21 | 20 | 19 | 19 |
| Total clusters | 361 | 357 | 363 | 352 | 365 | 358 | 379 | 371 | 367 | 344 | 353 | 349 | 341 | 336 | 330 | 283 | 282 | 283 |
| Score | 37.9 | 40.9 | 38.1 | 38.8 | 39.0 | 40.5 | 39.2 | 42.0 | 40.1 | 34.6 | 37.7 | 38.3 | 33.7 | 36.0 | 35.8 | 31.6 | 32.2 | 31.1 |

**Figure 6.**
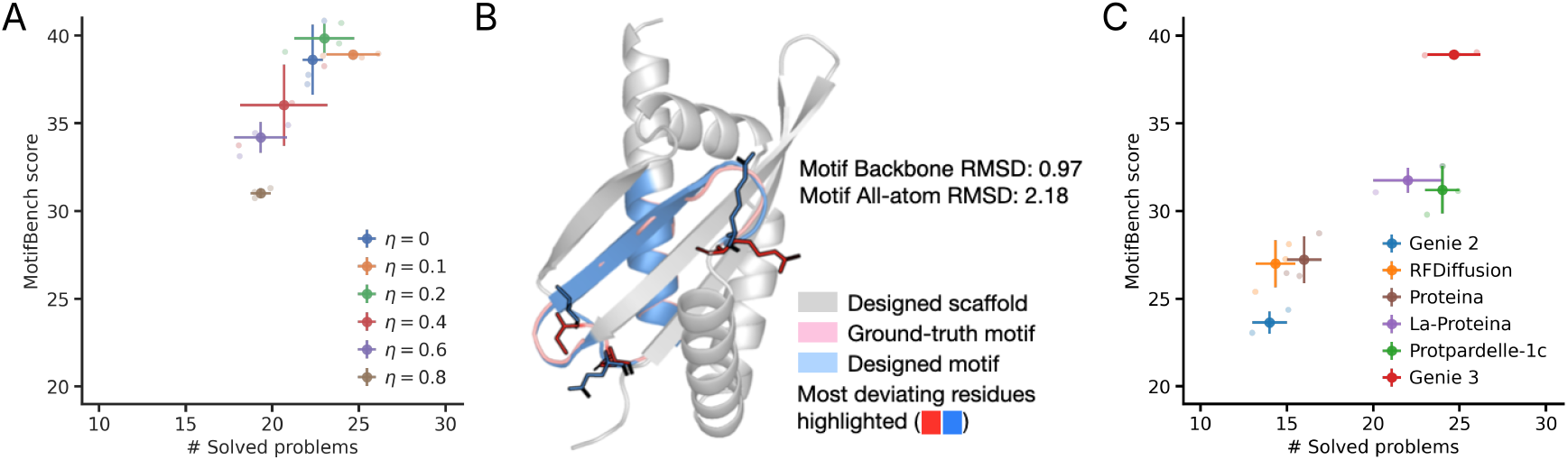
Additional information on MotifBench performance. (A) Performance of Genie 3 on MotifBench using different directional scales *η*, assessed using MotifBench score (y-axis) as a function of the number of solved MotifBench problems (x-axis). Dark dots correspond to mean diversity of three repeated samplings (light dots) with crosses indicating the extent of one standard deviation. (B) Example of motif scaffolding design with valid backbone (motif backbone RMSD *<* 1Å) but incorrect side chain arrangements (motif all-atom RMSD *>* 2Å). Here, we see close backbone alignment between the target motif (pink) and design motif (blue). However, there are significant deviations in sidechain arrangements for several motif residues (shown in sticks). (C) MotifBench performance with an additional constraint of motif all-atom RMSD *<* 2Å, which enforces motif sidechain agreement. Performance is assessed using MotifBench score (y-axis) as a function of the number of solved MotifBench problems (x-axis). Dark dots correspond to mean diversity of three repeated samplings (light dots) with crosses indicating the extent of one standard deviation.

**Figure 7.**
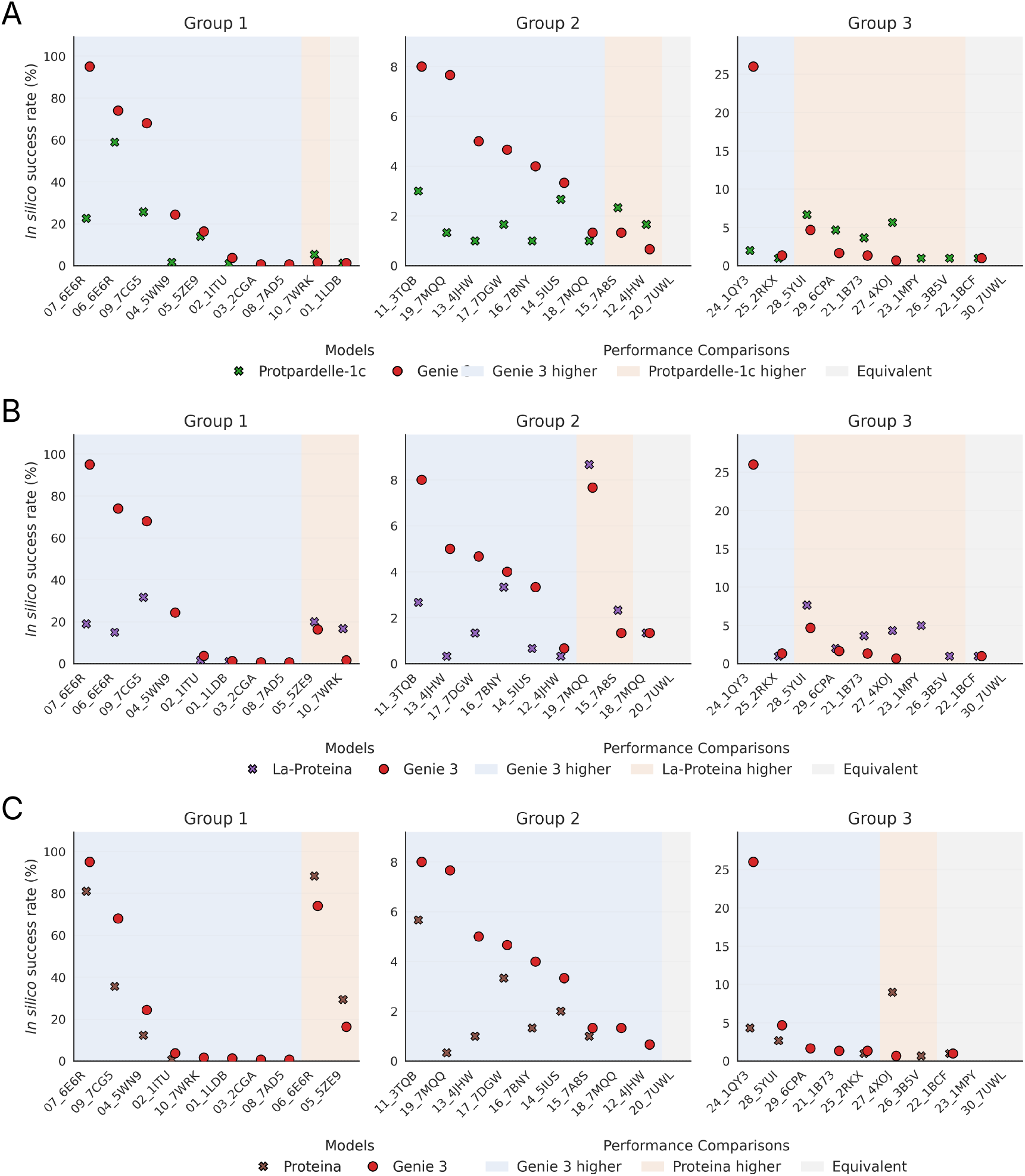
Comparison of Genie 3 with Protpardelle-1c (A), La-Proteina (B) and Proteina (C) on MotifBench performance. Here, we report the number of unique (TM-score *≤* 0.6) successful designs out of 100 design attempts. We omit data points if the model is unable to generate any successful design for a given problem. Problems that Genie 3 performs best on, problems that a competing method performs best on, and problems that Genie 3 and a competing method show equivalent performance on, are colored blue, orange, and grey, respectively.

**Figure 8.**
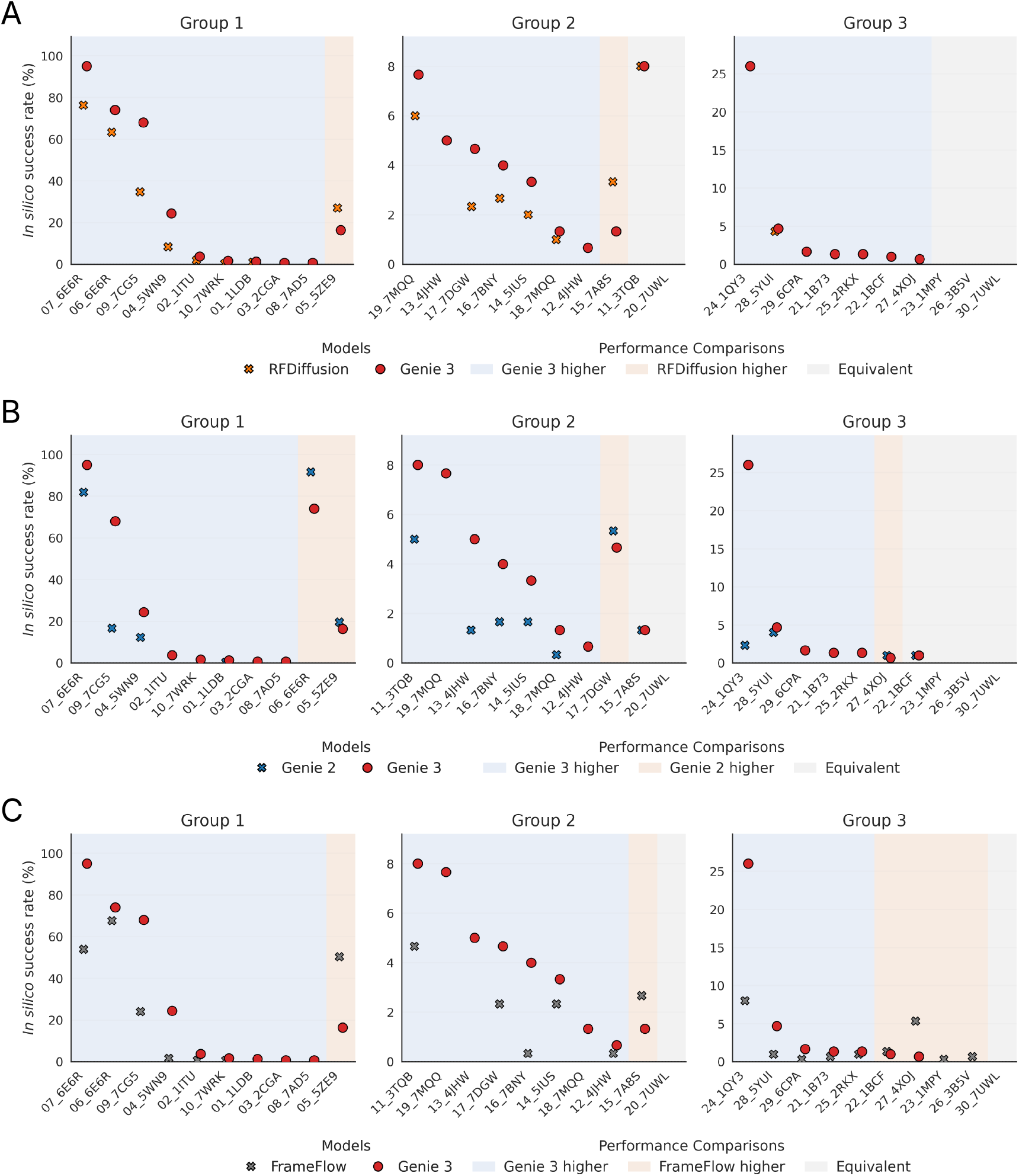
Comparison of Genie 3 with RFDiffusion (A), Genie 2 (B) and FrameFlow (C) on MotifBench performance. Here, we report the number of unique (TM-score *≤* 0.6) successful designs out of 100 design attempts. We omit data points if the model is unable to generate any successful design for a given problem. Problems that Genie 3 performs best on, problems that a competing method performs best on, and problems that Genie 3 and a competing method show equivalent performance on, are colored blue, orange, and grey, respectively.

It is worth noting that when computing motif agreements and evaluating successes for MotifBench, the aligned RMSD is computed based on motif backbone atoms (N, Ca, C and O). This poses an issue since a design with validated backbone structure could have incorrect side-chain configurations, inhibiting its ability to perform its desired function (Figure 6B). Thus, similar to other works on motif scaffolding (Geffner et al., 2025a; Lu et al., 2025), we introduce an additional all-atom constraint on motifs, which requires motif all-atom RMSD to be below 2Å, and call the corresponding successes “all-atom successes” (more details in Appendix C.1). Figure 6C visualizes motif scaffolding performance on MotifBench with an additional all-atom success criteria, which is consistent with the performance on MotifBench reported in Figure 2B.

## D. Assessment of *In silico* Binder Evaluation Pipeline

In this section, we provide further details on the retrospective analysis of the Cao dataset, including dataset information (Appendix D.1), summary of different binder design evaluation pipelines (Appendix D.2), detailed statistics of AF2M benchmark on Cao dataset (Appendix D.3), and limitations of the *in silico* binder evaluation pipeline (Appendix D.4). We also perform a retrospective analysis on the set of experimental binders reported by Zambaldi et al. (2024) and analyze the effect of different target conditions on the filtering power of the AF2M+ Benchmark that we use in this work (Appendix D.5).

### D.1. Cao dataset

Binder candidates were generated by placing individual amino acid residues predicted to favorably interact with the target hotspot—identified using RifGen—onto miniprotein scaffold backbones. This initial set was then refined through a targeted search procedure, where high-quality interface motifs were identified, grouped based on structural similarity, and reinserted into scaffold frameworks to preferentially sample effective binding configurations. Candidate designs were ranked using Rosetta-based scoring criteria and subsequently validated experimentally by expressing the corresponding synthetic genes on yeast cell surfaces, followed by enrichment of binders using fluorescence-activated cell sorting (FACS). Binding affinity was estimated using the midpoint concentration (SC50) of the binding response, and binders were classified using a threshold of SC50 *<* 4*μ*M.

### D.2. etails on different binder design evaluation pipelines

An *in silico* binder design evaluation pipeline takes a designed binder with sequence and structure information and determines if it is an *in silico* success based on various selected filters. Specifically, the sequence of the designed binder is refolded using a structure prediction model and success is determined based on confidences from the prediction model (binder quality) and the consistency between the predicted and designed structures (model consistency). There are three key components in each *in silico* binder design evaluation pipeline, including the use of structure prediction model, the use of target structure information, and the selection of filters. We compare different *in silico* binder design evaluation pipelines that are commonly used in the literature in Table 11, with details on selection filters summarized in Table 12. Note that since AF2 does not natively support multimers, the binder and target are inputted as a monomer with a gap of 200 residues in between.

**Table 10.** Summary statistics for binder design problems in the Cao dataset, consisting of experimentally tested binders and non-binders across 11 binder design problems.

| Problem | Target PDB | Target Chain | Confirmed Binders | Confirmed Non-binders | Total | Target Description |
| --- | --- | --- | --- | --- | --- | --- |
| InsulinR | 4OGA | E | 259 | 59,714 | 59,973 | Insulin receptor domains L1-CR |
| FGFR2 | 1DJS | A | 604 | 59,395 | 59,999 | Fibroblast growth factor receptor 2 |
| SARS-CoV-2 | 6M0J | E | 18 | 99,980 | 99,998 | SARS-CoV-2 spike protein |
| H3 | 3ZTJ | B | 50 | 59,891 | 59,941 | Hemagglutinin HA2 chain |
| IL-7RA | 3DI3 | B | 22 | 14,977 | 14,999 | Interleukin-7 receptor subunit alpha |
| PDGFR | 3MJG | X | 284 | 99,708 | 99,992 | Beta-type platelet-derived growth factor receptor |
| EGFR | 1MOX | A | 15 | 99,788 | 99,803 | Epidermal Growth Factor Receptor |
| TGFβ | 3KFD | A | 100 | 14,900 | 15,000 | Transforming growth factor beta-1 |
| TIE2 | 2GY7 | B | 5 | 99,995 | 100,000 | Angiopoietin-1 receptor |
| TrkA | 2IFG | A | 10 | 14,990 | 15,000 | Nerve growth factor receptor |
| VirB8 | 4O3V | A | 72 | 14,939 | 15,011 | VirB8-like protein of type IV secretion system |

**Table 11.** Comparison of different *in silico* binder evaluation pipeline.

|  | Structure Prediction Model | Use of Target Structure | Selection Filters |
| --- | --- | --- | --- |
| AF2 Benchmark | AlphaFold 2 | Template and initialization | Level 1 |
| Optimized AF2 Benchmark | AlphaFold 2 | Template and initialization | Level 2 |
| AF3 Benchmark | AlphaFold 3 | Template | Level 3 |
| AF2M Benchmark | AlphaFold Multimer | Template and initialization | Level 3 |

**Table 12.** Comparison of different selection filters.

| Filter Group | Binder Quality | Model Consistency |
| --- | --- | --- |
| Level 1 | Average complex pLDDT > 80<br>Average interaction pAE < 10Å | Binder RMSD < 1Å |
| Level 2 | Average complex pLDDT > 90<br>Average interaction pAE < 7Å | Binder RMSD < 1.5Å |
| Level 3 | Binder pTM > 0.8<br>Minimum interaction pAE < 1.5Å | Complex RMSD < 2.5Å |

### D.3. Additional data on the performance of the AF2M benchmark on the Cao dataset

### D.4. Limitations of the *in silico* binder evaluation pipeline

Figure 9 provides a detailed comparison of *in silico* filtering power across benchmarks. While *in silico* binder evaluation pipelines help enrich for experimental successes, their precision is generally low across problems (maximum ~12%) and pipelines completely fail for several problems such as H3, TGF*β* and TIE2. The best performing combination of structure prediction models and filters also vary significantly by problem, although the extent is small due to low filtering power. In addition, strict filters generally improve pipeline filtering power at the cost of low recall (Figure 10), suggesting that a high percentage of successful binders are undetected by the *in silico* evaluation pipelines, and thus using any *in silico* evaluation pipeline to compare different generative models could be inherently problematic. It is also worth noting that potential distributional differences in sequence and structure space between the Cao dataset (generated through iterative searching and docking) and one induced by a generative model could further impact the reliability of the filtering power computed based on the Cao dataset. Despite these limitations, *in silico* binder evaluation pipelines are commonly used since they help enrich for experimental successes and cut down on experimental costs when validating thousands of potential binders.

**Figure 9.**
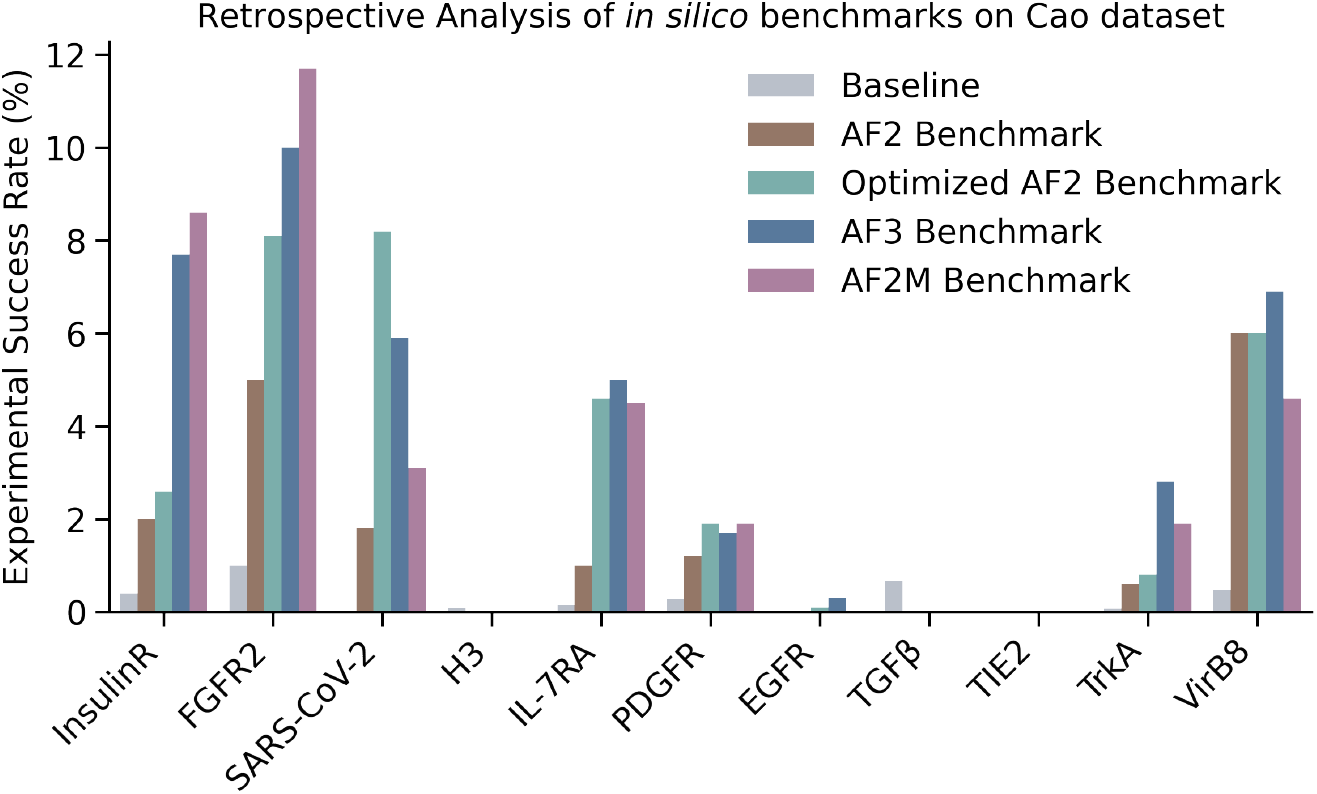
Comparisons of different *in silico* binder design evaluation pipelines on the Cao dataset, which consists of experimentally validated binders and non-binders across 11 different binder design problems. We report the percentage of true binders among *in silico* successes determined by each evaluation pipeline (corresponding to precision). Details on different pipelines are provided in Appendix D.2.

**Figure 10.**
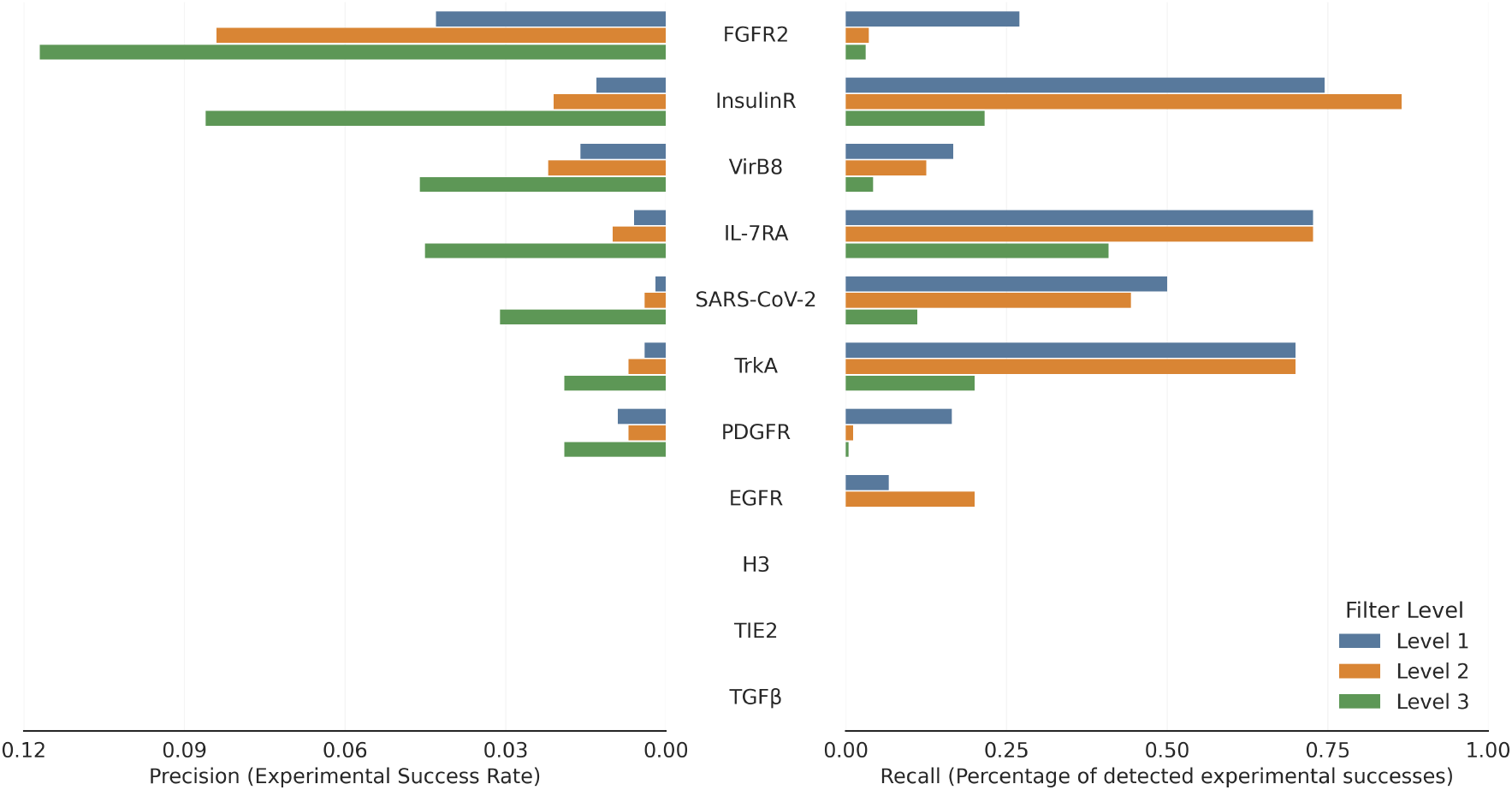
Comparisons of AF2M Benchmark with different sets of filters on the Cao dataset. The left bar plot visualizes precision, which is the percentage of true binders among *in silico* successes determined by the AF2M-based evaluation pipeline. The right bar plot visualizes the recall, which is the percentage of detected true binders among all true binders. Full retrospective analysis data for AF2M Benchmark is provided in Table 13.

### D.5. Effect of conditioning using different target information in AF2M on *in silico* filtering

We conduct additional retrospective analysis on the set of experimental successes reported by Zambaldi et al. (2024), which consists of 3-6 experimentally validated binders (with only sequence information) across a subset of binder design problems that we used for evaluation. Though this evaluation set is small, it provides us with useful insights on the effect of using different target information as conditions in AF2M on *in silico* filtering, mainly because it consists of challenging multimeric targets such as VEGF-A and IL17A. We compare between conditioning target information using MSAs and using templates to AF2M. While both conditioning strategies perform similarly across most cases, we find that the template-based approach fails to discover all experimental binders using binder quality filters in AF2M Benchmark (specifically binder pTM *>* 0.8 and minimum interaction pAE *<* 1.5Å). We suspect that this could be due to the lack of multimeric template support in AF2M and the richness of information in MSAs compared to monomeric templates.

**Table 13.** Retrospective performance of AF2M-based binder evaluation pipeline on the Cao dataset.

| Problem | Binders | Non-binders | Level | TP | FP | TN | FN | Precision | Recall | F1 |
| --- | --- | --- | --- | --- | --- | --- | --- | --- | --- | --- |
| InsulinR | 259 | 59,714 | L1 | 193 | 14226 | 45488 | 66 | 0.013 | 0.745 | 0.026 |
|  |  |  | L2 | 224 | 10614 | 49100 | 35 | 0.021 | 0.865 | 0.040 |
|  |  |  | L3 | 56 | 595 | 59119 | 203 | 0.086 | 0.216 | 0.123 |
| FGFR2 | 604 | 59,395 | L1 | 163 | 3628 | 55767 | 441 | 0.043 | 0.270 | 0.074 |
|  |  |  | L2 | 22 | 241 | 59154 | 258 | 0.084 | 0.036 | 0.051 |
|  |  |  | L3 | 19 | 144 | 59251 | 585 | 0.117 | 0.031 | 0.050 |
| SARS-CoV-2 | 18 | 99,980 | L1 | 9 | 5712 | 94268 | 9 | 0.002 | 0.500 | 0.003 |
|  |  |  | L2 | 8 | 2011 | 97969 | 10 | 0.004 | 0.444 | 0.008 |
|  |  |  | L3 | 2 | 63 | 99917 | 16 | 0.031 | 0.111 | 0.048 |
| H3 | 50 | 59,891 | L1 | 0 | 0 | 59891 | 50 | 0.000 | 0.000 | 0.000 |
|  |  |  | L2 | 0 | 0 | 59891 | 50 | 0.000 | 0.000 | 0.000 |
|  |  |  | L3 | 0 | 0 | 59891 | 50 | 0.000 | 0.000 | 0.000 |
| IL-7RA | 22 | 14,977 | L1 | 16 | 2553 | 12424 | 6 | 0.006 | 0.727 | 0.012 |
|  |  |  | L2 | 16 | 1584 | 13393 | 6 | 0.010 | 0.727 | 0.020 |
|  |  |  | L3 | 9 | 192 | 14785 | 13 | 0.045 | 0.409 | 0.081 |
| PDGFR | 284 | 99,708 | L1 | 47 | 5234 | 94474 | 237 | 0.009 | 0.165 | 0.017 |
|  |  |  | L2 | 3 | 441 | 99267 | 281 | 0.007 | 0.011 | 0.008 |
|  |  |  | L3 | 1 | 51 | 99657 | 283 | 0.019 | 0.004 | 0.006 |
| EGFR | 15 | 99,788 | L1 | 1 | 17784 | 82004 | 14 | 0.000 | 0.067 | 0.000 |
|  |  |  | L2 | 3 | 12715 | 87073 | 12 | 0.000 | 0.200 | 0.000 |
|  |  |  | L3 | 0 | 1852 | 97936 | 15 | 0.000 | 0.000 | 0.000 |
| TGFβ | 100 | 14,900 | L1 | 0 | 0 | 14900 | 100 | 0.000 | 0.000 | 0.000 |
|  |  |  | L2 | 0 | 0 | 14900 | 100 | 0.000 | 0.000 | 0.000 |
|  |  |  | L3 | 0 | 0 | 14900 | 100 | 0.000 | 0.000 | 0.000 |
| TIE2 | 5 | 99,995 | L1 | 0 | 907 | 95017 | 5 | 0.000 | 0.000 | 0.000 |
|  |  |  | L2 | 0 | 25 | 95899 | 5 | 0.000 | 0.000 | 0.000 |
|  |  |  | L3 | 0 | 64 | 95860 | 5 | 0.000 | 0.000 | 0.000 |
| TrkA | 10 | 14,990 | L1 | 7 | 1909 | 13081 | 3 | 0.004 | 0.700 | 0.007 |
|  |  |  | L2 | 7 | 953 | 14037 | 3 | 0.007 | 0.700 | 0.014 |
|  |  |  | L3 | 2 | 104 | 14886 | 8 | 0.019 | 0.200 | 0.034 |
| VirB8 | 72 | 14,939 | L1 | 12 | 758 | 14181 | 60 | 0.016 | 0.167 | 0.029 |
|  |  |  | L2 | 9 | 398 | 14541 | 63 | 0.022 | 0.125 | 0.038 |
|  |  |  | L3 | 3 | 62 | 14877 | 69 | 0.046 | 0.042 | 0.044 |

## E. Binder Design

### E.1. Evaluation dataset

We use the set of binder design problems curated by Zambaldi et al. (2024) to assess different generative models, with details summarized in Table 14 and visualizations on target structure and hotspot residues provided in Figure 12. Two minor adjustments are made: for H1, we use the full chain definitions available in the PDB file; for IL-17A, we truncate a few (2-4) residues in each terminal end of chain A to ensure both chains of the homodimer target have the same length, making it easier for chain permutations in the evaluation pipeline.

**Table 14.**
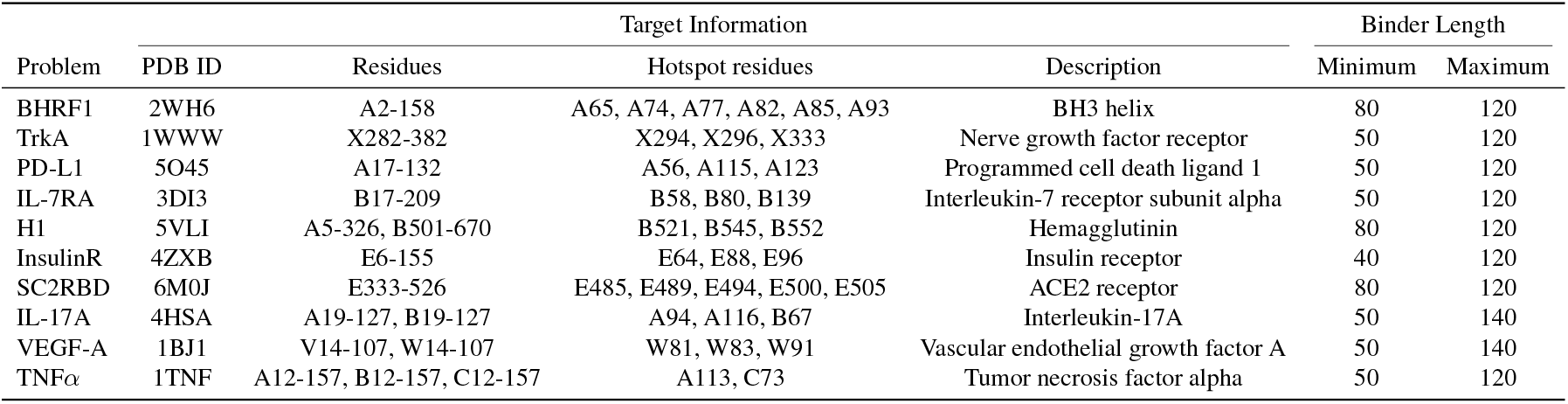
Set of binder design problems adapted from AlphaProteo.

**Figure 11.**
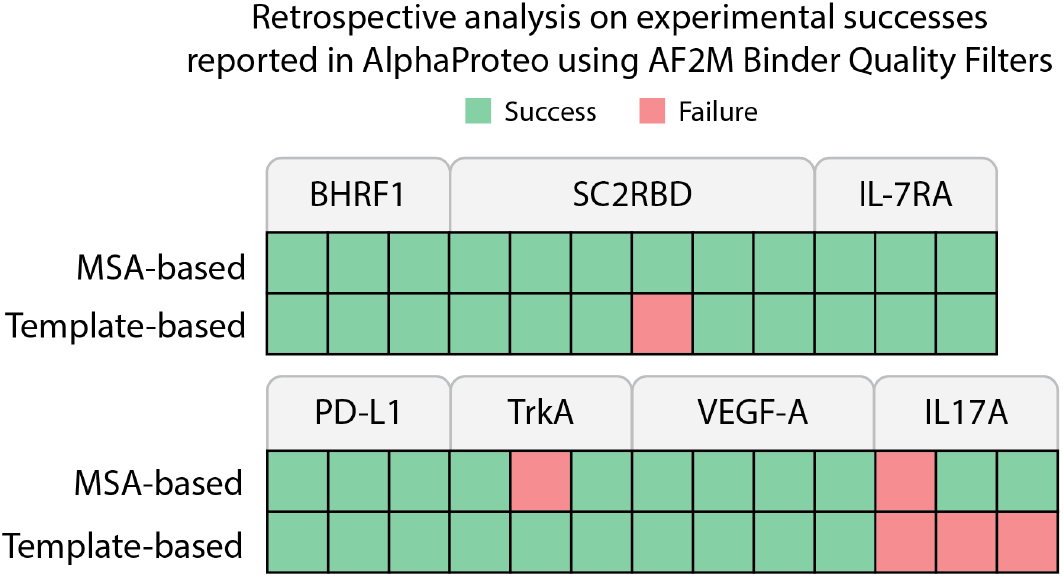
Comparisons of different target conditioning information to AF2M on the set of experimental binders reported by Zambaldi et al. (2024), demonstrating the importance of conditioning via MSAs of target sequence on detection of experimental binders.

**Figure 12.**
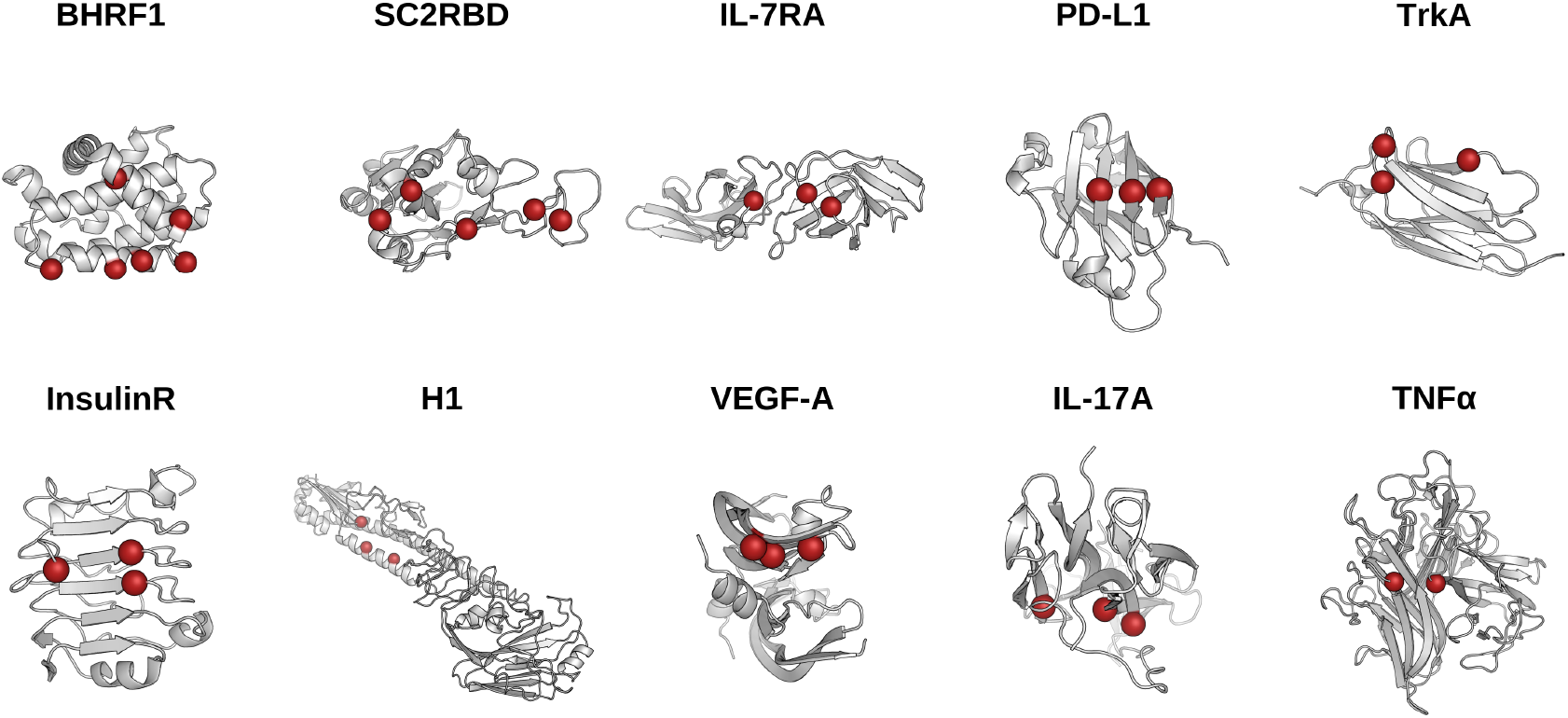
Visualization of target structures in the binder design problem set, with target hotspot residues highlighted in red.

### E.2 Reproduction of baseline methods

We provide details on the reproduction of baseline binder design methods below.

#### RFDiffusion

We follow the instructions from RFDiffusion GitHub repository^17^. Figure 13 shows the unique success rate clustered at TM = 0.6 using hotspot residues as conditioning, demonstrating that sampling with a noise scale of 0 achieves better binder design performance than sampling with a noise scale of 1. This is consistent with the observation by Zambaldi et al. (2024). We thus sample with a noise scale of 0 when reproducing the performance of RFDiffusion.

**Figure 13.**
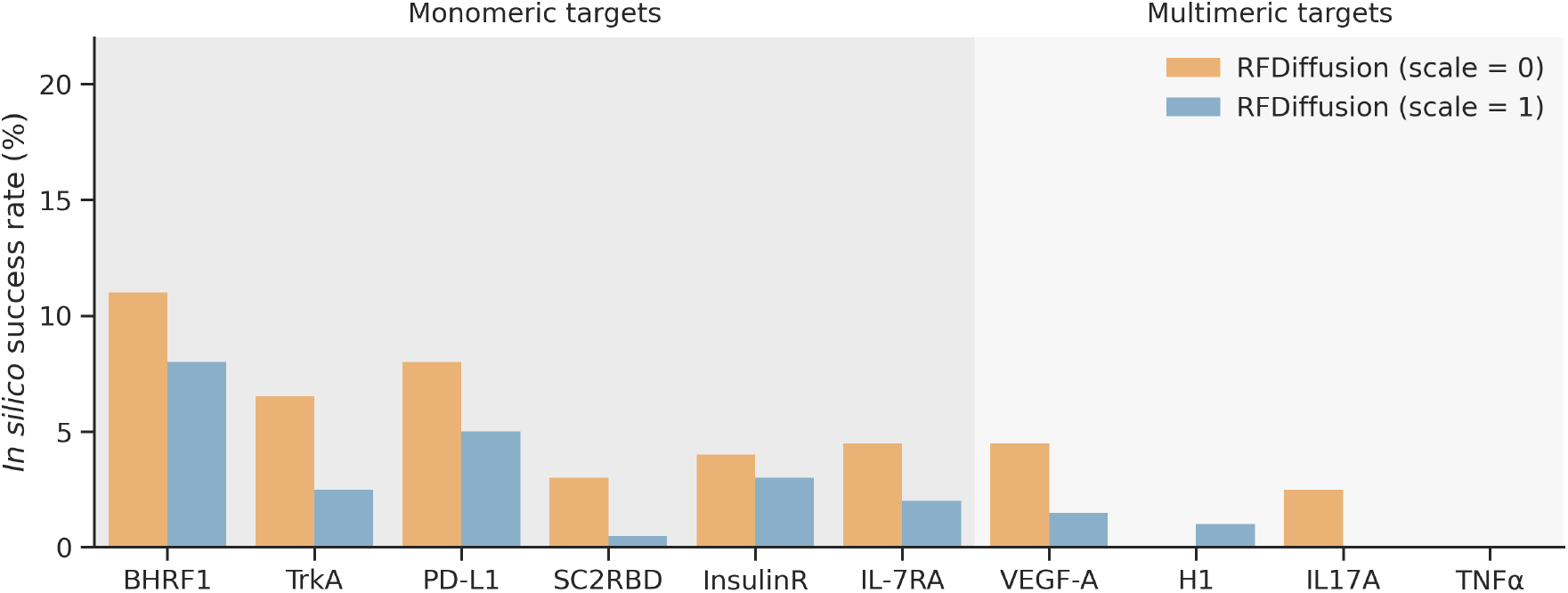
Fractional rate of generating unique (TM-score *≤* 0.6) successful designs (as assessed by AF2M+ Benchmark) out of 200 attempts per problem. Here, we use the hotspot residues as conditioning to the model since it does not show much performance difference between hotspot conditioning and extended/convergent conditioning for RFDiffusion.

**Figure 14.**
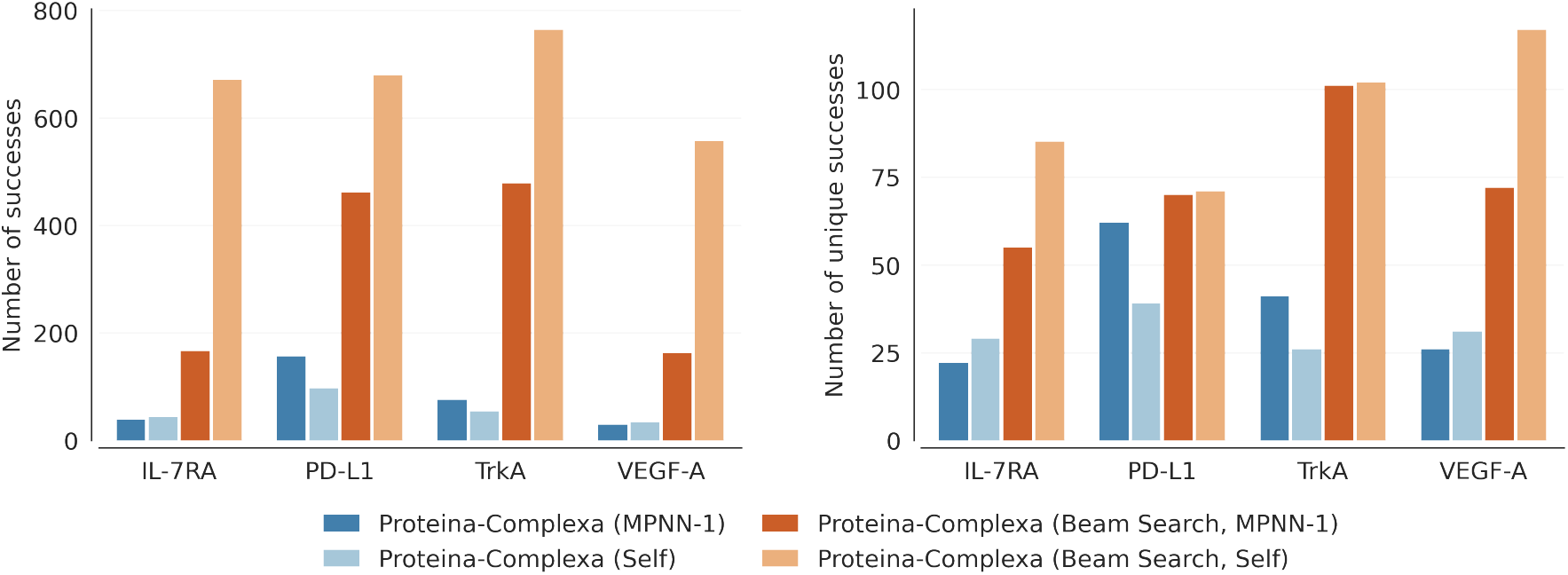
*In silico* binder design performance across four Proteina-Complexa model variants, measured in terms of number of total successes (left) and number of unique successes (right, clustered at a TM threshold of 0.6). It consists of a 2*×*2 factorial design crossing two structure sets (with and without beam search) against two sequence sources (self-generated and ProteinMPNN).

#### BoltzGen

We follow the instructions from the BoltzGen GitHub repository^18^ and use the default `protein-anything`setting. For consistent comparison with other methods, we only perform the design stage, omitting the inverse-folding and structure prediction stages.

#### Proteina-Complexa

We follow the instructions from the Complexa GitHub repository^19^ and use the default protein binder setting and checkpoints. Throughout the manuscript, self refers to using the Complexa-generated sequence without redesign, whereas MPNN refers to sequence redesign using ProteinMPNN. We take the generated structure and sequence and evaluate them using our standard evaluation pipeline rather than the Complexa evaluation pipeline. For beam search, we use the AF2Multimer reward model with standard settings, `beam_width`set to 4, and `n_branch`set to 4.

#### BindCraft

We follow the instructions from the BindCraft GitHub repository^20^ and use the default 4-stage multimer setting. For consistent comparison with other methods, we disable ProteinMPNN and only run the first stage (sequence optimization). Note that a sampling trajectory could be terminated early if its corresponding computational metrics are far above the filter thresholds. This means that to generate 200 designs using BindCraft, more than 200 trajectories are generated. In addition, BindCraft optimizes a sequence against computational metrics from AlphaFold Multimer (AF2M), which is also the “oracle” structure prediction model used in the *in silico* binder evaluation pipeline. This could potentially advantage BindCraft.

### E.3. Self-Generated versus ProteinMPNN Sequences for Proteina-Complexa

Didi et al. (2026) demonstrate that Proteina-Complexa’s co-generated sequences outperform ProteinMPNN sequences through high-throughput experimental validation. However, this comparison conflates two distinct sources of optimization. In the beam search setting, self-generated sequences are produced *jointly* with their input structures using AlphaFold-Multimer (AF2M) confidence as a reward signal — meaning both the sequence and the structure are optimized end-to-end. When ProteinMPNN is applied to the beam search outputs, it operates on the same optimized structures but without access to the AF2M signal during sequence generation, creating an asymmetric comparison.

To disentangle these contributions, we construct a 2*×* 2 factorial design crossing two structure sets (with and without beam search) against two sequence sources (self-generated and ProteinMPNN):

- **Proteina-Complexa (Self)**: structures without beam search, self-generated sequences
- **Proteina-Complexa (MPNN-1)**: structures without beam search, ProteinMPNN sequences
- **Proteina-Complexa (Beam Search, Self)**: structures with beam search, self-generated sequences
- **Proteina-Complexa (Beam Search, MPNN-1)**: structures with beam search, ProteinMPNN sequences

This design allows us to isolate contributions coming from sequence and structure optimization. Results across four binder design targets (IL-7RA, PD-L1, TrkA, VEGF-A) on which Proteina-Complexa performs best experimentally (Didi et al., 2026) yield three observations:

1. **Without beam search, ProteinMPNN sequences perform comparably or better than self-generated sequences** (Proteina-Complexa (MPNN-1) vs. Proteina-Complexa (Self)). Since neither condition benefits from AF2M-guided optimization, this provides a fair baseline comparison between the two sequence sources and suggests that self-generated sequences carry no inherent advantage over ProteinMPNN-generated sequences.
2. **Beam search independently improves performance regardless of sequence source** (Proteina-Complexa (Beam Search, MPNN-1) vs. Proteina-Complexa (MPNN-1)). Holding the sequence model fixed at ProteinMPNN, beam search increases total and unique success rates, demonstrating the effect of structural optimizations through beam search.
3. **Within the beam search setting, self-generated sequences outperform ProteinMPNN** (Proteina-Complexa (Beam Search, Self) vs. Proteina-Complexa (Beam Search, MPNN-1)). Since both conditions use identical beam-search-optimized structures and observation 1 rules out an inherent sequence-level advantage, this residual gap isolates a genuine benefit of sequence co-optimization with AF2M during beam search. In other words, the self-generated sequences benefit not because they are better sequences *per se*, but because they were optimized *together* with their structures under the same AF2M objective.

Taken together, these results reframe the original paper’s conclusion: the performance advantage of self-generated sequences is not a property of the sequence model itself, but a result of joint sequence-structure optimization under AF2M guidance. ProteinMPNN, applied *post hoc* to beam searched structures without access to the same optimization signal, is not a fair counterfactual. Nevertheless, this experiment establishes beam search with self-generated sequences as the optimal setting for running Proteina-Complexa and we thus use it for inference-time scaling comparisons with Genie 3.

### E.4. Performance without additional hotspot coverage constraints

Figure 15 shows that while all models generally respect the specified interface region, BindCraft struggles to conform to it on more challenging multimeric targets, where a sizeable fraction of otherwise successful designs fail to engage the correct interface. When hotspot expansion heuristics are applied, however, BindCraft recovers in some cases such as H1 and IL17A — producing more successful designs that satisfy the interface constraint — suggesting the model benefits from a more refined interface definition as prior knowledge. On the other hand, BindCraft performs worse on problems such as VEGF-A when using hotspot expansion heuristics, suggesting it is being overly constrained by a rigid binding pose.

**Figure 15.**
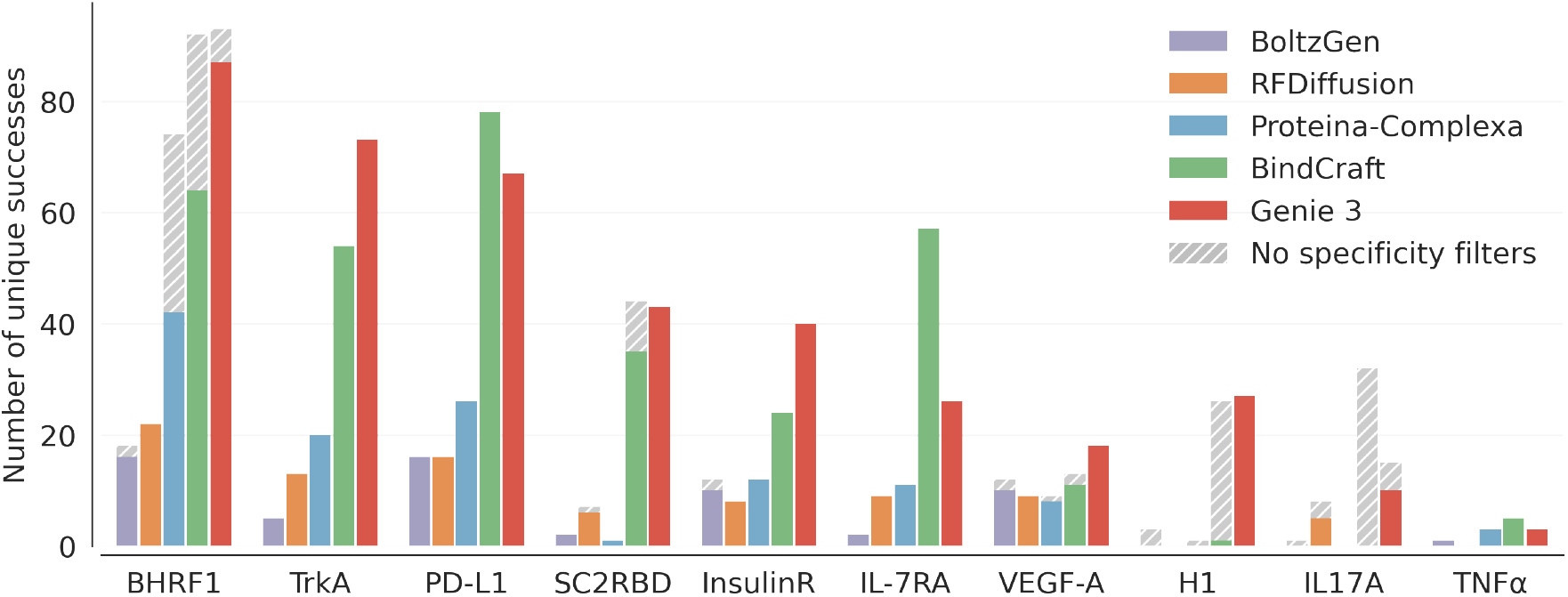
*In silico* binder design performance measured as the number of unique (TM-score *≤* 0.6) and successful (AF2M+ Benchmark) binder designs. Shaded regions indicate the fraction of otherwise successful designs that do not bind their user-specified interface.

**Figure 16.**
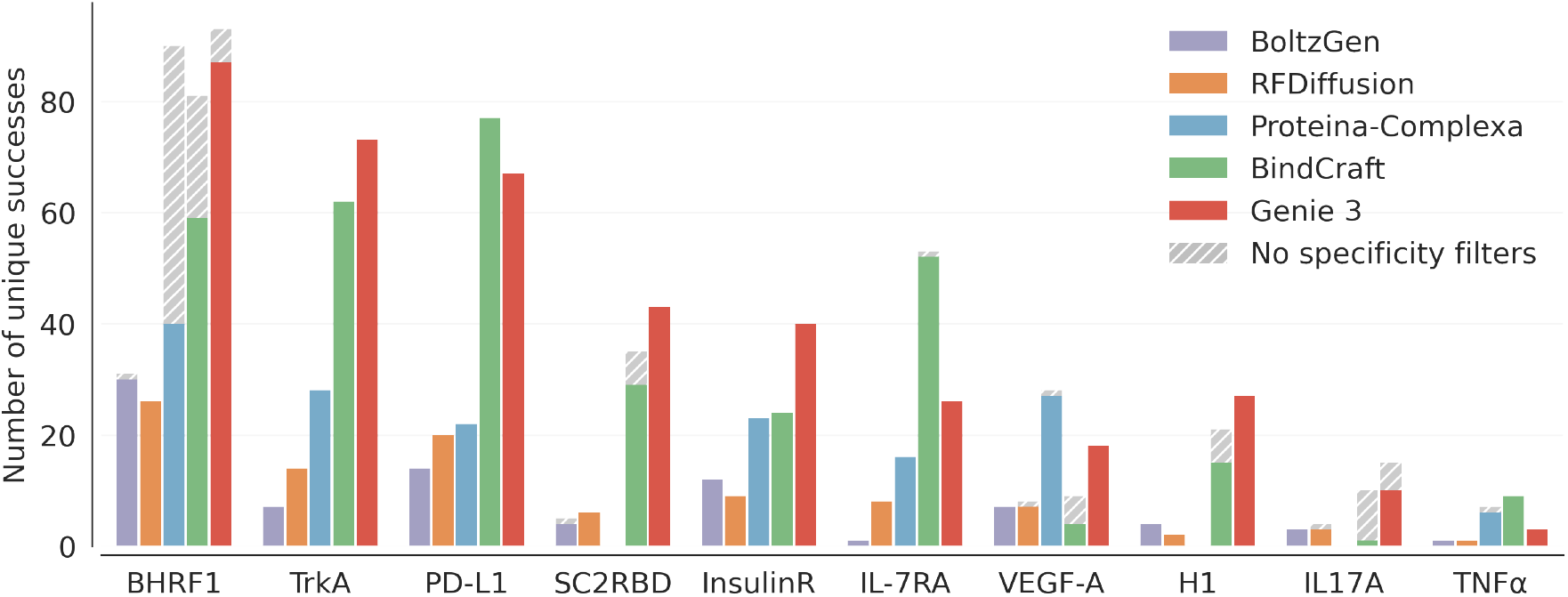
*In silico* binder design performance measured as the number of unique (TM-score *≤* 0.6) and successful (AF2M+ Benchmark) binder designs, where the same hotspot expansion heuristics (extended and convergent hotspots for monomeric and multimeric targets, respectively) are applied to all methods. Shaded regions indicate the fraction of otherwise successful designs that do not bind their user-specified interface.

**Figure 17.**
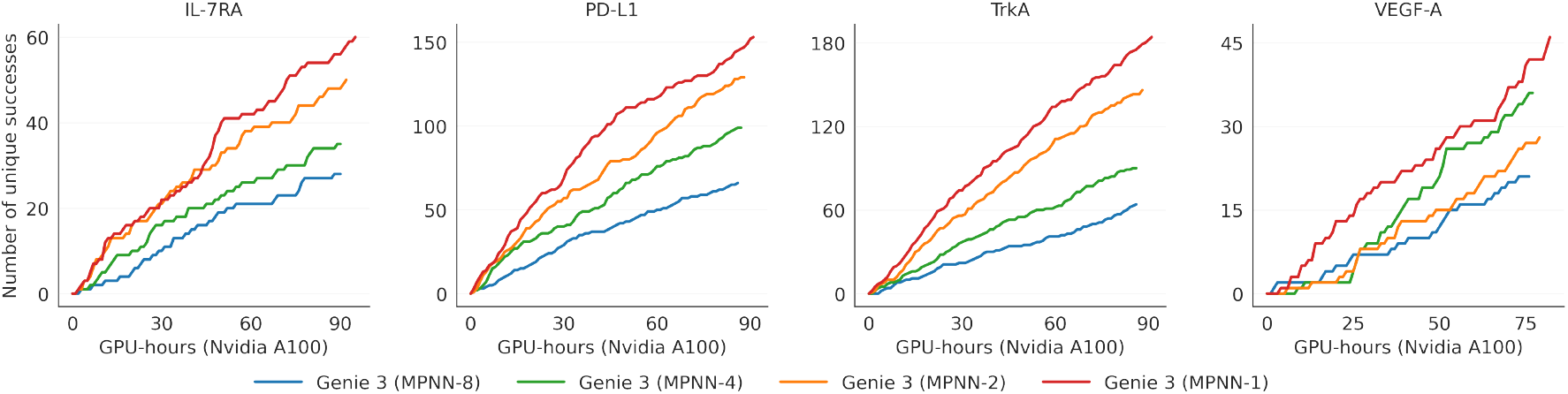
Tradeoff between structural and sequence diversity in Genie 3, visualized as the number of unique successes as a function of Nvidia A100 GPU-hours. Here, each model variant is given an evaluation budget of 1,600 ColabFold calls. Decreasing the number of ProteinMPNN sequences per structure trades off sequence diversity for structural diversity.

### E.5. Structural versus Sequence Diversity in Genie 3

Since total runtime is dominated by ColabFold evaluation (20 recycles, 5 models, *~* 250s per structure) rather than structure generation (*~* 20s) or inverse folding (*~* 1s), we investigate the optimal allocation of a fixed computational budget between structural and sequence diversity. Specifically, we fix the total number of ColabFold calls to 1,600 and vary the number of generated structures and ProteinMPNN calls per structure inversely:

- 200 generated structures, 8 ProteinMPNN calls each
- 400 generated structures, 4 ProteinMPNN calls each
- 800 generated structures, 2 ProteinMPNN calls each
- 1600 generated structures, 1 ProteinMPNN call each

We report unique successes as a function of total Nvidia A100 GPU-hours, including both generation and evaluation, across four binder design targets (3 monomeric, 1 multimeric). The trend is consistent across all targets: maximizing structural diversity — 1600 generated structures with a single ProteinMPNN call each — yields the best inference-time scaling at negligible additional runtime cost. This indicates that structural diversity is the primary driver of success rate for Genie 3, and that expanding sequence diversity through multiple ProteinMPNN calls offers diminishing returns relative to generating more structures.

### E.6. Visualizations of different interface conditioning heuristics at inference time

### E.7. Assessments based on using hotspot expansion heuristics for baseline models

Figure 19 visualizes the effect of applying hotspot expansion heuristics for baseline models by problem. We observe that hotspot expansion generally benefits other generative models such as BoltzGen, RFDiffusion and Proteina-Complexa, though the improvement is minimal with the largest gain observed in Proteina-Complexa. The effect of hotspot expansion on BindCraft is mixed. We next compare Genie 3 with baseline models, all of which use the same set of expanded hotspots as Genie 3. Here, we adopt the same setting as fixed sampling budget experiment, where 200 structures are sampled for each problem-model combination and each generated structure is inversely folded with ProteinMPNN (8 sequences per structure). We observe a similar trend to Figure 3C, where Genie 3 performs best in 6 out of 10 problems.

**Figure 18.**
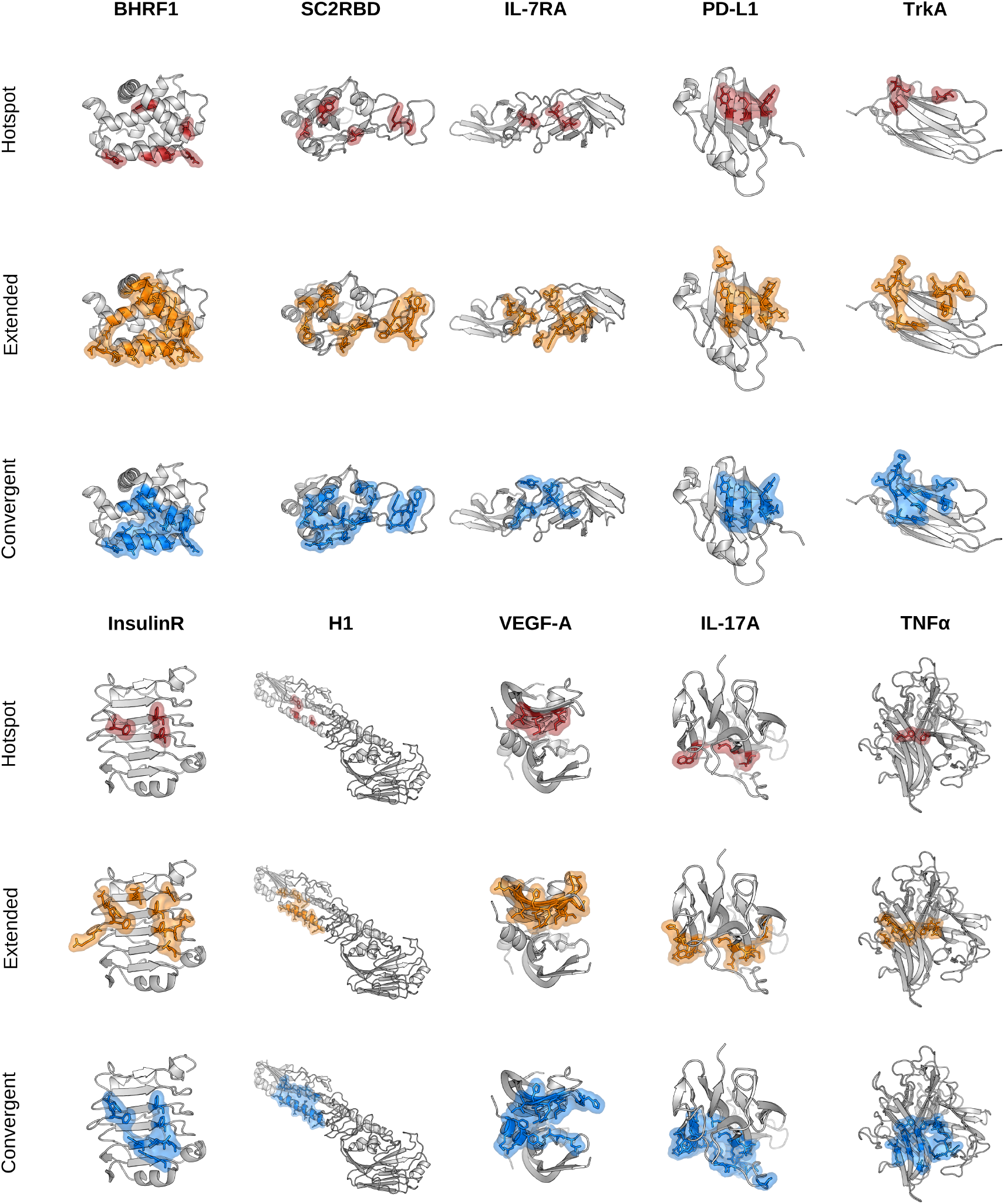
Visualization of the three interface definitions used across all targets: default **hotspot** residues, **extended** interfaces including nearby surface residues, and **convergent** interfaces derived from residues in successful designs.

**Figure 19.**
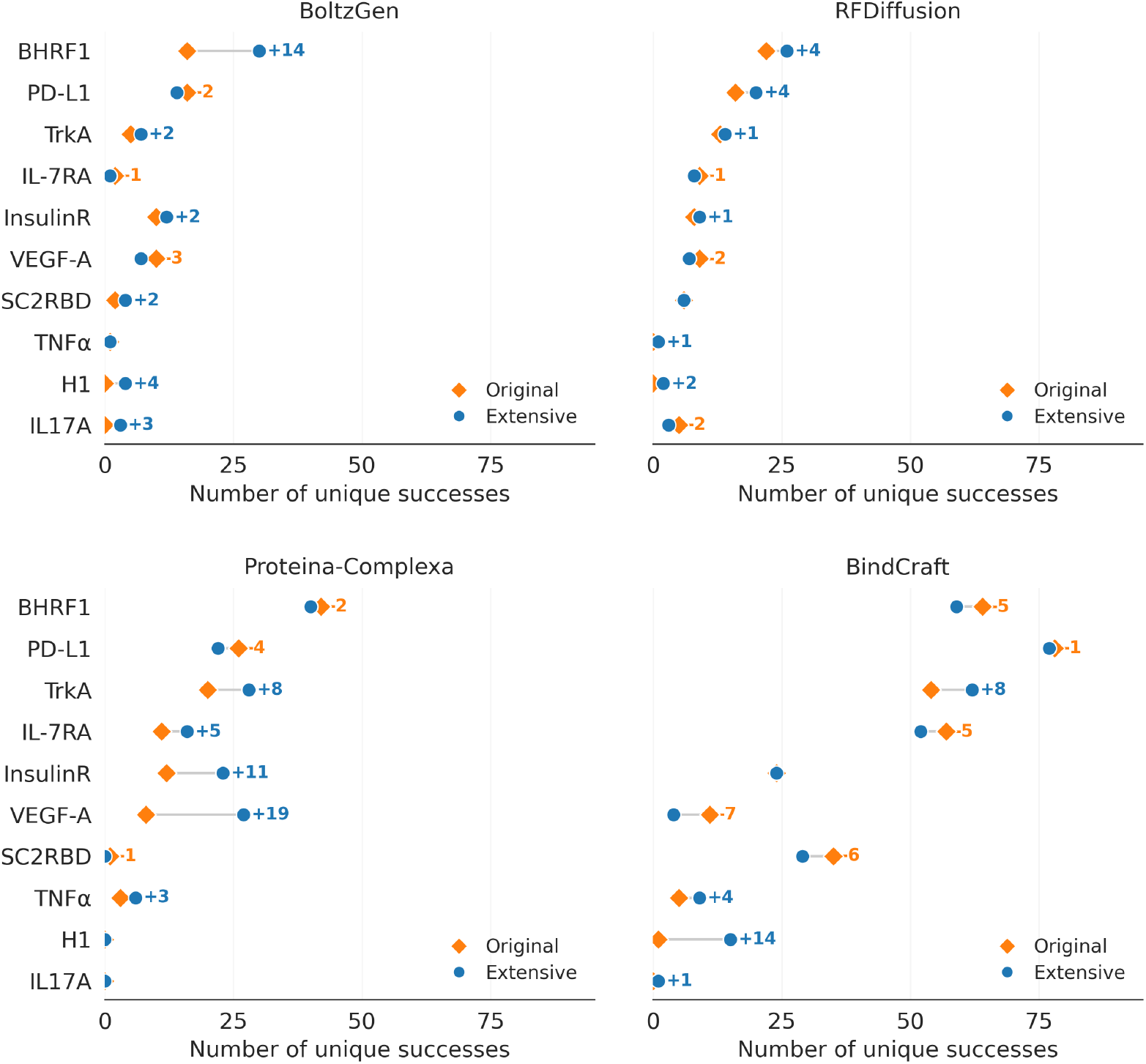
Effect of using hotspot expansion heuristics for baseline models. For each model-problem combination, we sample 200 structures and perform sequence redesign using ProteinMPNN (8 sequences per structure). Number of unique (TM-score *≤* 0.6) and successful (AF2M+ Benchmark) binder designs is reported and compared. Here, original hotspots corresponds to initial user-specified hotspots and extensive hotspots corresponds to hotspots computed through our expansion heuristics (extended and convergent for monomeric and multimeric targets, respectively).

### E.8. Iterative Interface Refinement with Convergent Heuristic

When extending the binding interface through multiple rounds of binder design, we aim to enrich residues associated with higher success rates while still allowing room for diverse binders that target slightly different hotspots. If we naively take the union of all interface residues observed in previous successful designs, the interface definition quickly becomes overspecified as new successes introduce additional target residues. Conversely, taking the intersection leads to underspecification: a single outlier success can eliminate genuinely important interface residues that do not appear universally. To address this, we introduce a probabilistic interface definition. We assign each target residue a probability of being specified a hotspot equal to the frequency with which it appears in the binding interface across all successful complexes accumulated up to the current round.

Algorithm 4 details our iterative interface refinement algorithm. In Figure 4D, we show the results of applying Algorithm 4 to binder generation against TNF*α*. The number of successful binders increases across rounds, reaching 9 successes by round 5, compared to only 1 success when the interface is not refined.

#### Algorithm 4

Iterative Interface Refinement

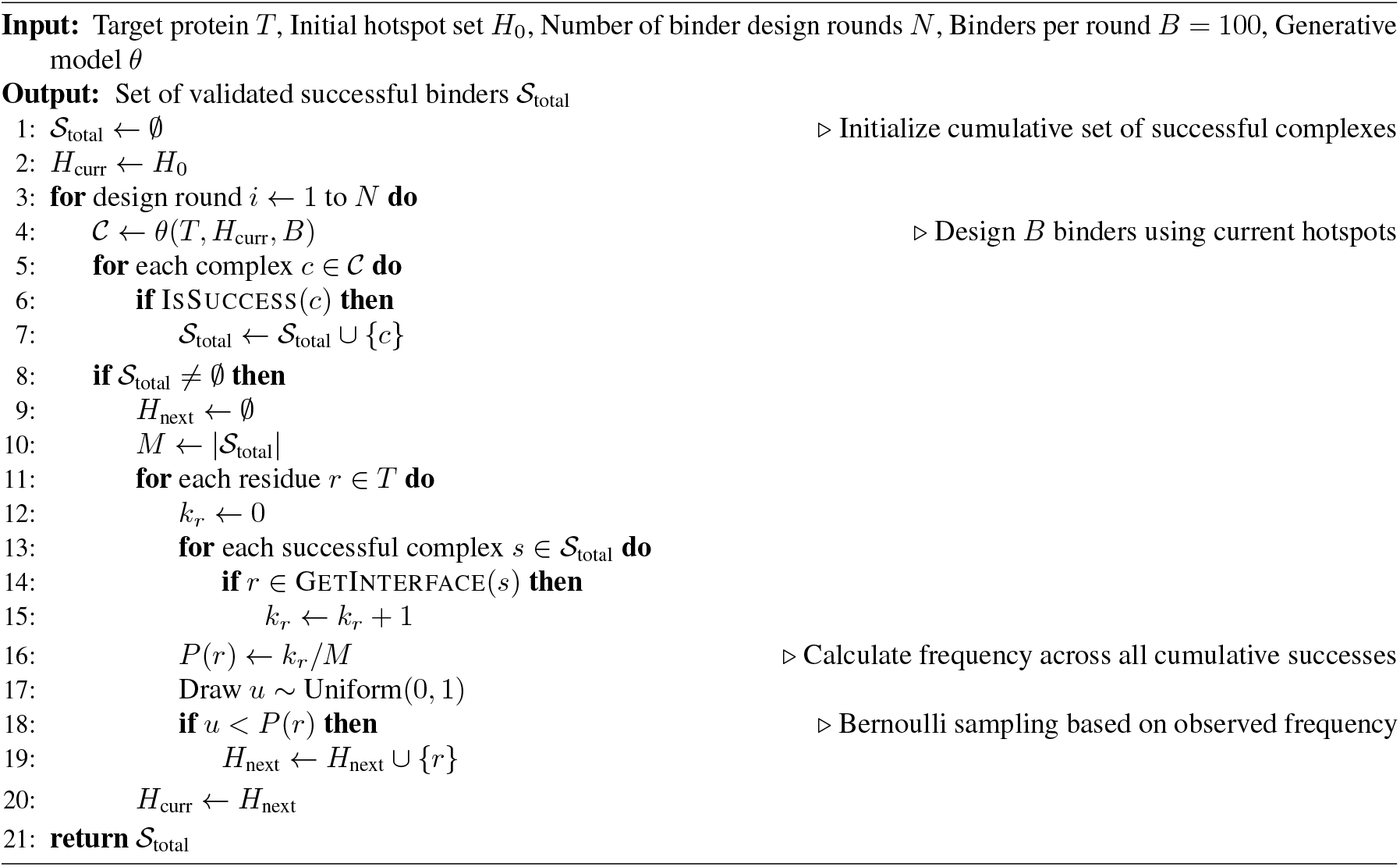

### E.9. Profiling of per-sample runtime

We consider per-sample runtime as a sum of generation time, inverse folding time (if applicable), and structure prediction time.

#### E.9.1. Generation time

We profile generation time on binder design using a single A100 GPU (40GB memory) and average the inference time of a single sample over 10 runs. Here, we report the generation time by problem since the length of target proteins varies by problems and the length of binder proteins varies by samples. For Genie 3, we disable model compilation since the variation in binder lengths cause dynamic shapes for model input and thus constant model recompilation once the input shape is changed. One potential way to fix this is to ensure constant input shape via padding. Figure 21 demonstrates that Genie 3 is reasonable among generative models, generating more slowly than Proteina-Complexa but faster than BoltzGen and RFDiffusion. Genie 3 is also significantly faster than BindCraft, a hallucination-based approach.

**Figure 20.**
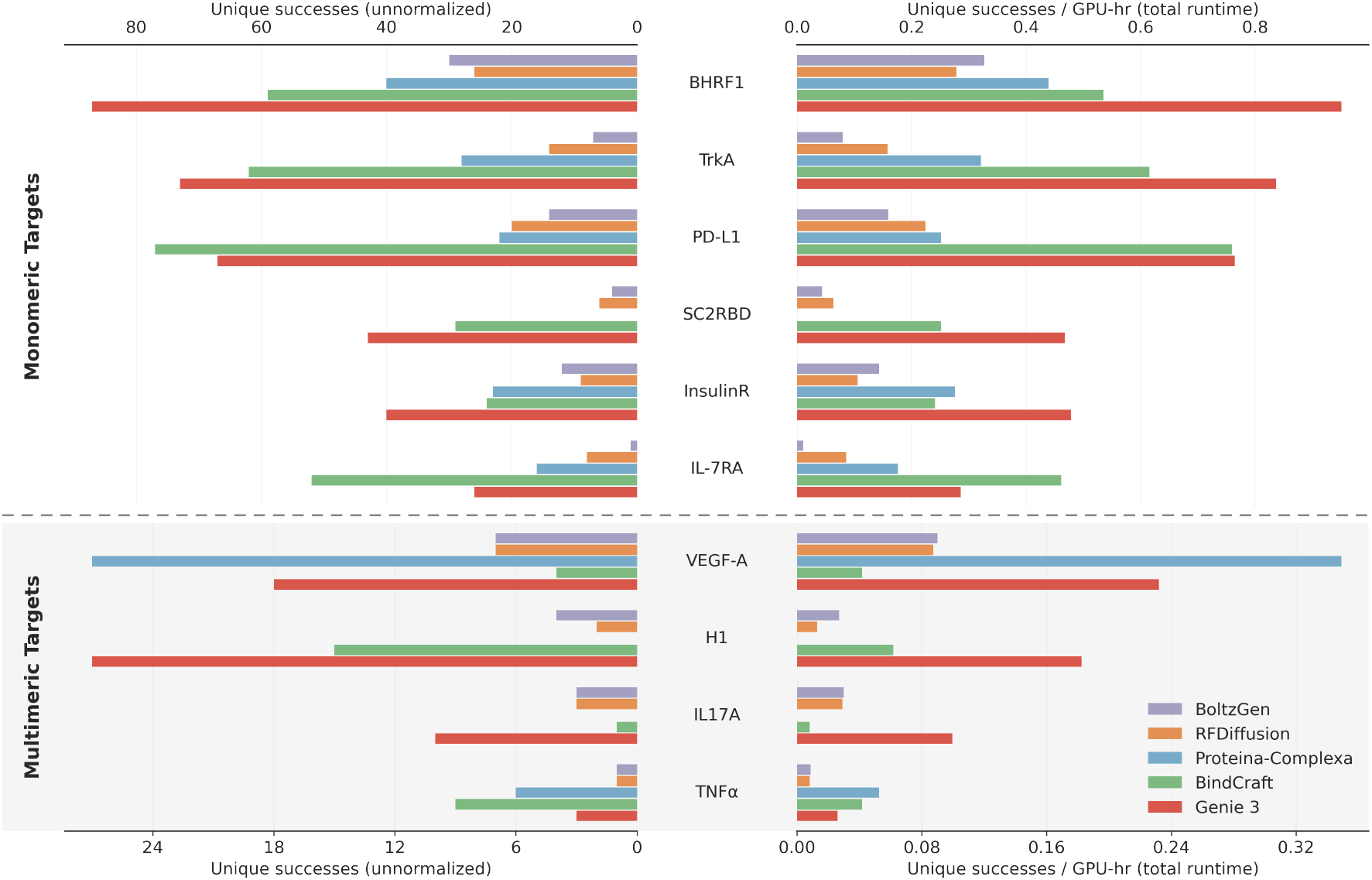
Number of unique (TM-score *≤* 0.6) successful designs (as assessed by AF2M+ Benchmark) out of 200 attempts per problem for Genie 3 and four leading binder design methods (left: unnormalized; right: normalized by number of Nivida A100 GPU hours used for generation and evaluation). Here, all models use the same set of expanded hotspots for generation (extended and convergent hotspots for monomeric and multimeric targets, respectively).

**Figure 21.**
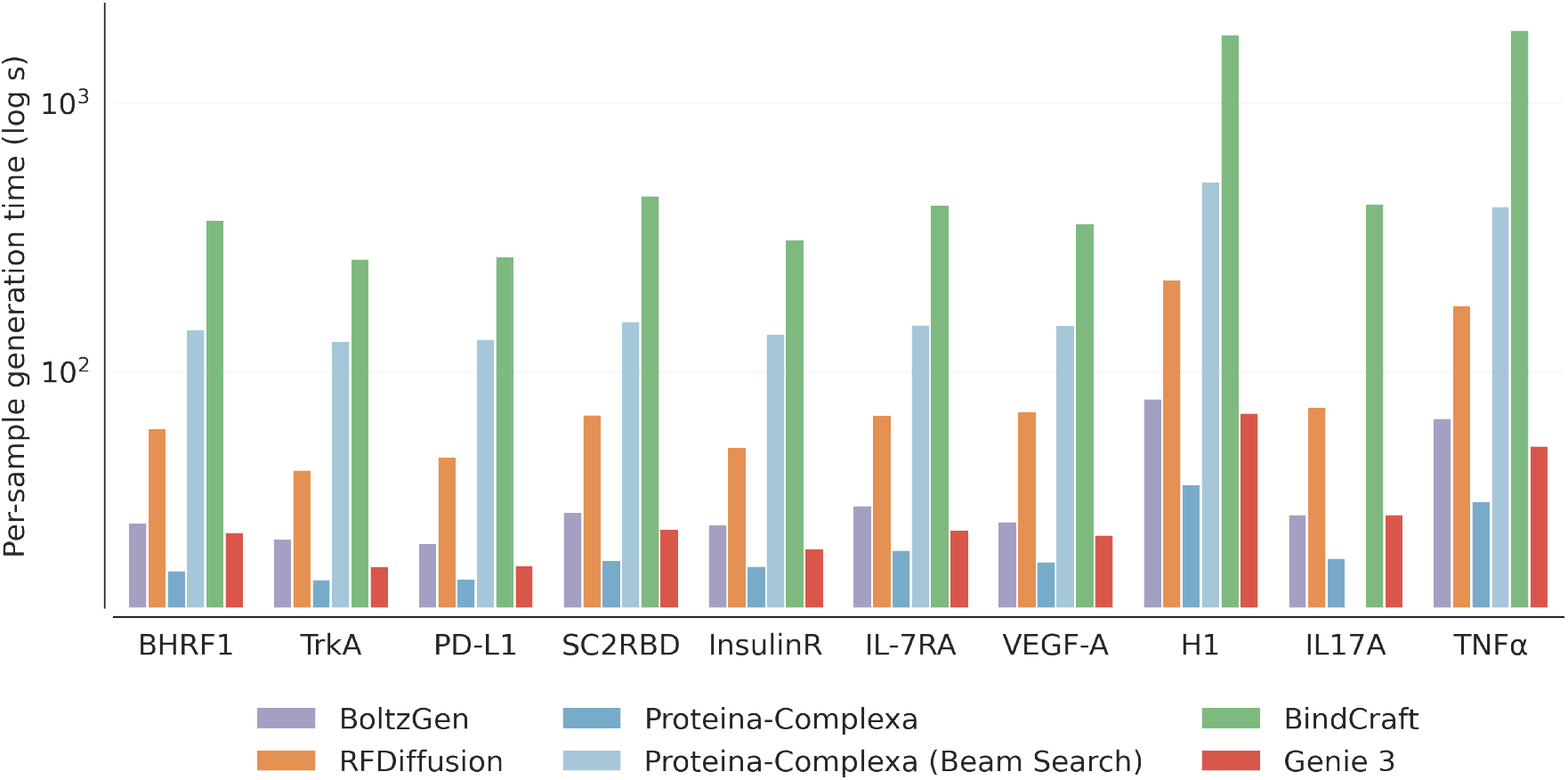
Genie 3 is superior in sampling speed, which is crucial for preparing a large-scale design pool for *in silico* filtering and screening. Here, we visualize the per-sample generation time in log scale, with full data summarized in Table 15.

#### E.9.2. Inverse folding time

We profile inverse folding time for binder design on a single 40GB A100 GPU and report the mean per-sample runtime for each target across 20 Genie 3-generated binders, with one sequence inverse folded per sample. All measurements are taken after 5 warmup runs.

**Table 15.** Comparison of sampling time on binder design (in seconds).

| Model | BHRF1 | TrkA | PD-L1 | SC2RBD | InsulinR | IL-7RA | VEGF-A | H1 | IL-17A | TNFα |
| --- | --- | --- | --- | --- | --- | --- | --- | --- | --- | --- |
| BindCraft | 356 | 256 | 260 | 442 | 301 | 404 | 345 | 1771 | 411 | 1844 |
| RFDiffusion | 54 | 37 | 42 | 61 | 44 | 58 | 62 | 202 | 65 | 160 |
| BoltzGen | 20 | 18 | 17 | 22 | 19 | 21 | 19 | 63 | 21 | 52 |
| Proteina-Complexa | 11 | 11 | 11 | 12 | 11 | 11 | 11 | 22 | 12 | 18 |
| with beam search | 142 | 129 | 131 | 153 | 137 | 148 | 147 | 504 | 153 | 409 |
| Genie 3 | 18 | 13 | 13 | 18 | 14 | 15 | 16 | 54 | 21 | 38 |

**Table 16.** Comparison of inverse folding time on binder design (mean per-sample time in seconds).

| Model | BHRF1 | TrkA | PD-L1 | SC2RBD | InsulinR | IL-7RA | VEGF-A | H1 | IL-17A | TNFα |
| --- | --- | --- | --- | --- | --- | --- | --- | --- | --- | --- |
| ProteinMPNN | 0.88 | 0.71 | 0.72 | 0.97 | 0.97 | 1.31 | 1.06 | 1.97 | 1.03 | 1.83 |

**Table 17.**
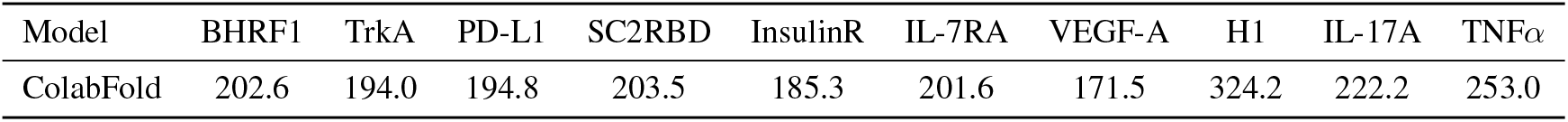
Comparison of structure prediction time on binder design (in seconds).

#### E.9.3. Structure prediction time

We profile structure prediction time on binder design by running ColabFold on the query consisting of target sequence(s) and a 100-residue binder with 20 recycles and 5 models, and report the runtime for each problem. Each measurement is taken once. Because AF2M uses early stopping, runtimes may vary across runs.

#### E.10. Additional details on binder design against NiV-G

The reported hit rates for submissions made using RFDiffusion, BindCraft, and BoltzGen are curated using the method tagging feature from the binder design competition website^21^. Note that there could potentially be designs generated using these methods that have missing method tags. In addition, these hit rates might not be accurate reflections of model performance due to differences and suboptimal choices in the use of generative approaches and filtering procedures.

## Footnotes

1 https://github.com/RosettaCommons/RFdiffusion

2 https://github.com/microsoft/protein-frame-flow

3 https://github.com/Wangchentong/Proteus

4 https://github.com/DreamFold/FoldFlow

5 https://github.com/NVIDIA-Digital-Bio/proteina

6 https://github.com/NVIDIA-Digital-Bio/la-proteina

7 https://github.com/ProteinDesignLab/protpardelle-1c

8 https://github.com/aqlaboratory/genie2

9 https://github.com/jozhang97/ambient-proteins

10 https://github.com/RosettaCommons/RFdiffusion

11 https://github.com/microsoft/protein-frame-flow

12 https://github.com/aqlaboratory/genie2

13 https://github.com/NVIDIA-Digital-Bio/proteina

14 https://github.com/NVIDIA-Digital-Bio/la-proteina

15 https://github.com/ProteinDesignLab/protpardelle-1c

16 https://github.com/blt2114/MotifBench

17 https://github.com/RosettaCommons/RFdiffusion

18 https://github.com/HannesStark/boltzgen

19 https://github.com/NVIDIA-Digital-Bio/Proteina-Complexa

20 http://github.com/martinpacesa/BindCraft

21 https://proteinbase.com/collections/nipah-binder-competition-results

